# Combinatorial and Inducible CRISPRa/i Enables Canalized hiPSC Forward Programming and Iterative Refinement *via* Single-Cell Genomics

**DOI:** 10.64898/2026.05.31.729073

**Authors:** Federica Sozza, Alberto Romano, Nicole D’Elia, Martina Terenzi, Maria Luisa Ratto, Emily R. Cliff, Gabrielle Nattenberg, Sara Bianchi, Silvia Becca, Heather Klug, Davide Cacchiarelli, Jesse G. Zalatan, Elisa Balmas, Alessandro Bertero

**Affiliations:** Molecular Biotechnology Center “Guido Tarone”, Department of Molecular Biotechnology and Health Sciences, University of Turin, Torino, Italy; Department of Chemistry, University of Washington, Seattle, WA, USA; Department of Laboratory Medicine & Pathology, University of Washington, Seattle WA USA; Telethon Institute of Genetics and Medicine (TIGEM), Pozzuoli; Department of Translational Medicine, University of Naples Federico II and Scuola Superiore Meridionale, Naples, Italy

## Abstract

Synthetic gene-regulation logic is established in immortalized cell lines but remains largely aspirational in human induced pluripotent stem cells (hiPSCs) and derivatives. This gap constrains both mechanistic discovery and translational engineering in physiologically relevant models. We developed **CIRI** (**C**ombinatorial **I**nducible C**R**ISPR in **I**PSCs), an isogenic, safe-harbor–engineered platform in which tetracycline-responsive single guide RNAs (sgRNAs) carry modular RNA aptamers that recruit RNA-binding proteins and effector domains. This design enables multimodal regulation from a single catalytically inactive Cas9 (dCas9), exemplified by orthogonal CRISPR activation and interference (CRISPRa/i). After optimizing sgRNA–aptamer architectures, we achieved robust CRISPRa and CRISPRi in hiPSCs and hiPSC-derived cardiac organoids. CIRI rapidly channels hiPSC forward programming into skeletal myocytes by activating *MYOD1* while repressing *NANOG*, *POU5F1/OCT4*, and *SOX2*. Combinatorial pooled dual-guide single-cell RNA sequencing screens identify *ID3* as a road-block and *KDM6B* and *SMARCD3* as synergistic enhancers of myogenic maturation. Together, CIRI establishes a programmable synthetic biology framework in human stem cell models.

**GRAPHICAL ABSTRACT:** 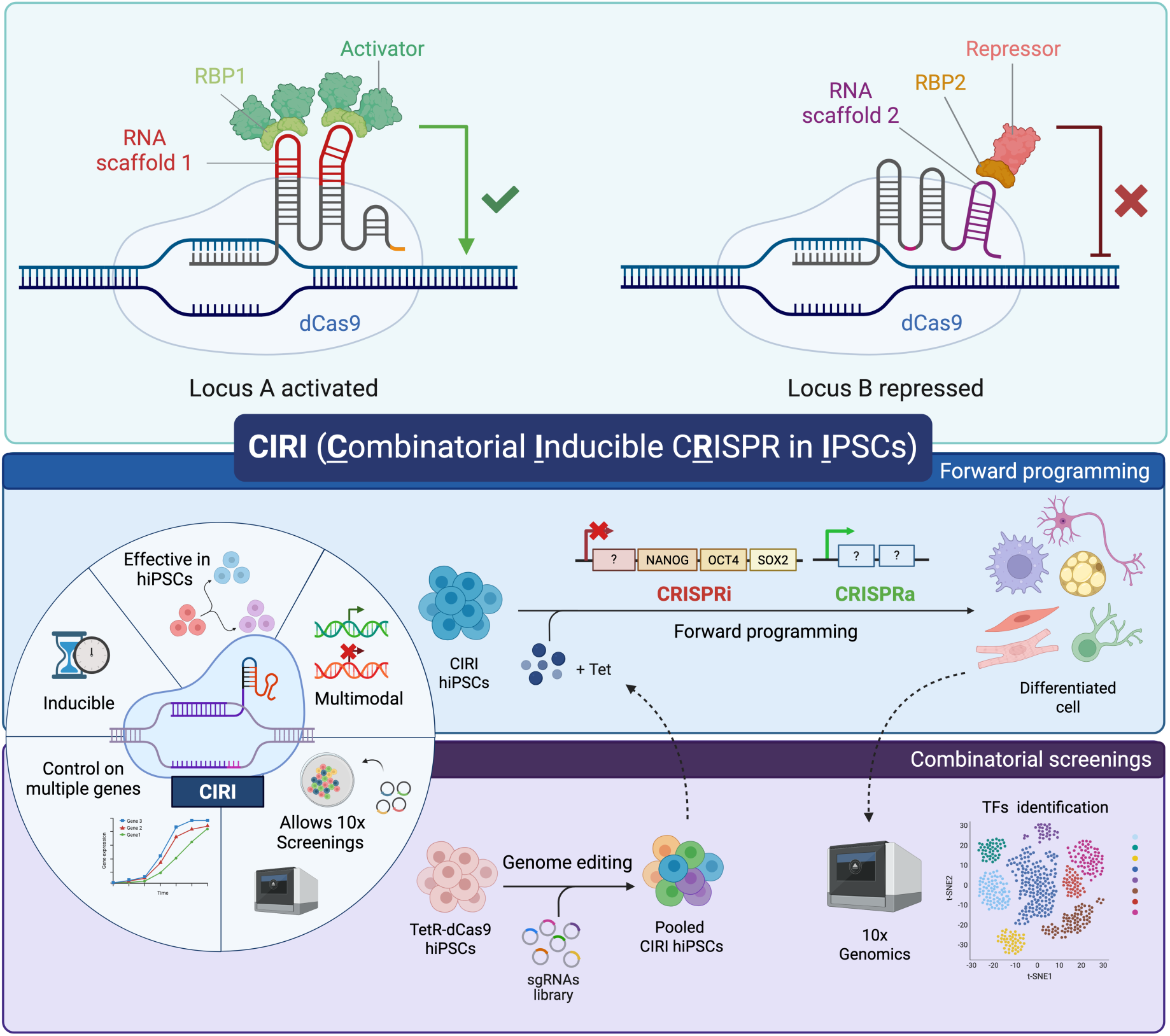

## INTRODUCTION

Synthetic biology aspires to recapitulate the remarkable precision, modularity, and adaptability that underlie natural gene regulatory programs. Among the tools driving this effort, Clustered Regularly Interspaced Short Palindromic Repeats (CRISPR)–Cas systems have endowed researchers with the ability to manipulate gene expression with unprecedented fidelity and versatility. Yet, despite their extraordinary success in immortalized cell lines^1–5^, these technologies have yet to fulfill their full potential when it comes to human induced pluripotent stem cells (hiPSCs)^6^, a model that uniquely captures human development and disease.

The advent of hiPSCs revolutionized biomedical research by providing a patient-specific, genetically matched source of human cell types without the ethical concerns associated with embryonic stem cells^7,8^. When combined with genome editing, hiPSCs offer extraordinary opportunities for precision medicine, disease modeling, and regenerative therapy^9^. However, the implementation of CRISPR technologies in hiPSCs has proven far more challenging than initially anticipated^10^. hiPSCs are intrinsically fragile, poorly clonogenic, and prone to silencing exogenous genetic material, making them unusually sensitive to the delivery, expression, and long-term stability requirements of CRISPR-based platforms. Moreover, standard CRISPR–Cas9 editing, which relies on double-strand DNA breaks, triggers p53-dependent apoptosis, thereby selecting for p53-deficient cells and compromising genomic stability and translational potential^11,12^. Efforts to circumvent these issues have focused on epigenetic editing strategies such as CRISPR activation (CRISPRa) or interference (CRISPRi)^4,13–20^. However, implementation in hiPSCs remains hindered by the poor expression and instability of large fusion proteins with catalytically inactive Cas9 (dCas9), as well as by epigenetic silencing of these components upon differentiation^21,22^. Robust temporal control of CRISPR systems is also pivotal in hiPSC models, both to prevent premature loss of stemness when perturbing cell fate-related genes, and to enable the study of gene function during specific developmental windows. However, widely used doxycycline-inducible DNA polymerase II promoters activate poorly after hiPSC differentiation, including when deployed in genomic safe harbors^23,24^.

All of these limitations compound when the goal moves from a simple gain/loss-of-function perturbation to simultaneous activation and repression of multiple genes. Such bidirectional, multiplexed control, which has been demonstrated in bacteria and in immortalized eukaryotic cells^25,26^, is essential, for instance, to reconstruct gene circuits that dictate lineage specification and dynamic transitions in cell fate. Multi-component designs exacerbate the challenges of hiPSC models by increasing cargo size, the number of exogenous elements to maintain, and the opportunities for variegated expression across clones and lineages. Overcoming these barriers requires rethinking how CRISPR components are assembled, delivered, and regulated. Here, we present **CIRI** (**C**ombinatorial **I**nducible C**R**ISPR in **I**PSCs), a flexible CRISPR/-dCas9 platform engineered to address these constraints. CIRI integrates tetracycline-inducible single-guide RNAs (sgRNAs) bearing modular RNA aptamers that recruit distinct regulatory effectors directly to their targets, enabling concurrent regulatory modalities (e.g., CRISPRa and CRISPRi) without the need for multiple dCas9 fusion proteins. Safe-harbor integration of silencing-resistant promoters ensures stable and homogeneous activity across differentiation. Finally, we demonstrate methods for single- and dual-guide combinatorial pooled single-cell RNA sequencing (scRNA-seq) screens to iteratively refine regulatory logic, applying them to achieve fast forward programming canalized toward skeletal muscle cell fate.

## RESULTS

### Engineering sgRNA scaffolds toward combinatorial CRISPRa/i in hiPSCs

We sought to develop a system that enables bidirectional and combinatorial gene regulation in hiPSCs while remaining practical to deploy across differentiation (with inducible control addressed in the next subsection). Accordingly, we first focused on identifying a flexible, modular, and multiplexable CRISPR architecture effective in hiPSCs. Several sgRNA scaffolds have been previously employed in CRISPR engineering. Notably, additional protein recruitment capabilities have been achieved through the usage of the well-characterized viral RNA sequences MS2, PP7, and com, which are recognized, respectively, by MCP, PCP, and Com RNA-binding proteins (RBPs)^25,27–29^. By fusing effectors to these RBPs (e.g., transcriptional activators or repressors), it is possible to directly recruit them to the desired genomic site. This strategy bypasses the need to engineer multiple large dCas9 effector fusions by recruiting compact RBP-effector modules to RNA hairpin-bearing sgRNAs, enabling multimodal, combinatorial regulation from a single dCas9 chassis and improving deployability in hiPSCs.

To select the optimal recruitment system for CRISPRa and CRISPRi applications in hiPSCs, we first tested various combinations of RNA hairpins and RBPs. To preserve compatibility with CRISPR-based screenings utilizing 3’ capture scRNA-seq (10X Genomics)^30^, each recruitment system was tested in combination with one of two possible “capture sequences” (cs1 and cs2) as well as in absence of any capture sequence. These tests were performed using a healthy donor male hiPSC line (WTC11) harboring a randomly integrated dCas9-2A-mCherry transgene (hereafter, “Cas-R”). This cell line was generated during an early attempt to integrate into the *CLYBL* genomic safe harbor (GSH) locus a bicistronic construct encoding dCas9 and mCherry as separate proteins linked by tandem self-cleaving 2A peptides. However, genomic analysis indicated that the plasmid was randomly integrated into the genome (Figure S1A). Despite this, the resulting cell line expressed moderate levels of the dCas9–mCherry transgene, as confirmed by flow cytometry and RT-qPCR (Figures 1A–1B).

**Figure 1.**
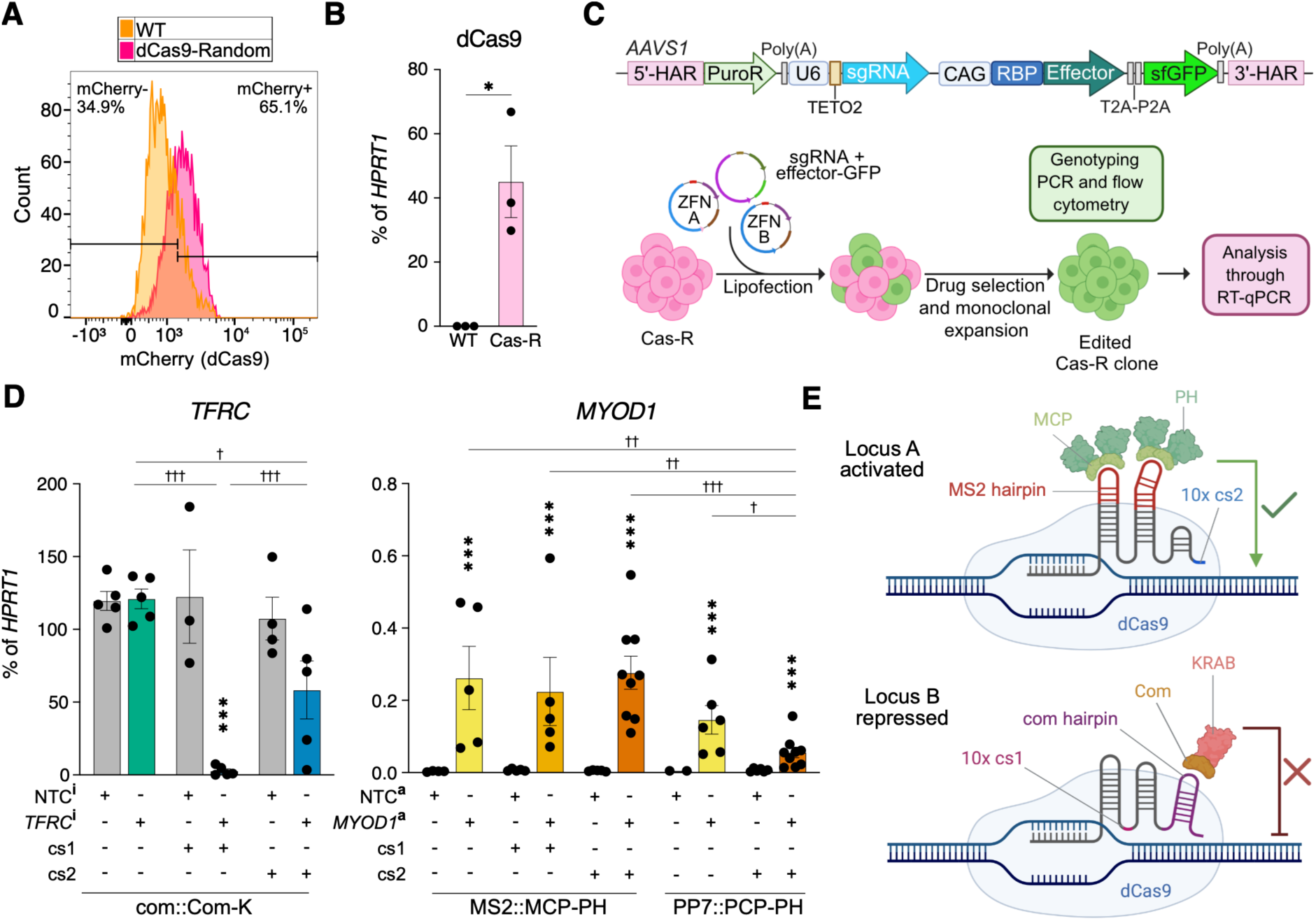
Optimization of sgRNA scaffolds for orthogonal CRISPRa/i in hiPSCs (A) Flow cytometry quantification of dCas9-2A-mCherry expression in WTC11 hiPSCs carrying a randomly integrated transgene, hereafter Cas9-R, compared with parental wild-type (WT) hiPSCs. (B) RT-qPCR quantification of dCas9 expression in Cas9-R hiPSCs. N = 3 successive passages; p = 0.0104 *versus* WT by unpaired two-tailed Welch’s t-test (*). (C) Workflow for sgRNA-scaffold optimization. Cas9-R hiPSCs were targeted at the *AAVS1* locus with cassettes encoding various sgRNA scaffolds and matched RBP–effector modules for CRISPRa or CRISPRi testing. HAR, homology arm; PuroR, puromycin N-acetyltransferase; Poly(A), bovine growth hormone polyadenylation signal; U6-TETO2, Pol III U6 promoter with a tet operator; CAG, cytomegalovirus immediate early enhancer/chicken beta-actin/rabbit beta-globin hybrid promoter; RBP, RNA-binding protein; T2A-P2A, tandem self-cleaving peptides; sfGFP, superfolder green fluorescent protein; ZFN, zinc-finger nuclease. Construct not shown to scale. (D) RT-qPCR analysis of target gene expression after CRISPRi of *TFRC* (*TFRC*^i^) or CRISPRa of *MYOD1* (*MYOD1*^a^), compared to matched controls (NTC^i/a^). cs1/2, 10X Genomics capture sequences 1/2; NTC, non-targeting control; K, KRAB; PH, p65-HSF1. N = 2–9 clones; two-way ANOVA with protospacer type and guide::effector architecture as factors, followed by Holm–Sidak’s multiple-comparison test for targeting *versus* NTC comparisons within each architecture (asterisks), and all pairwise architecture comparisons within the same guide type (daggers). *, † p < 0.05; **, †† p < 0.01; ***, ††† p < 0.001; p > 0.05 not reported. (E) Schematic of the selected orthogonal CRISPRa and CRISPRi modules. MS2-cs2 sgRNAs recruit MCP-PH for activation, whereas com-cs1 sgRNAs recruit Com-KRAB for repression, enabling distinct effectors to be recruited to different loci from the same dCas9 chassis. Note that MCP binds internal MS2 hairpins as a dimer, whereas Com binds as a monomer and requires the 3’-terminal com hairpin context, including the base of the stem.

WT-dCas9 hiPSCs were engineered at the *AAVS1* GSH with a cassette encoding a matched pair of sgRNA and RBP-effector-2A-sfGFP (superfolder GFP), both constitutively expressed. The RBP-effector cassette was driven by the CAG promoter, which we selected based on its established stable activity across hPSCs and multiple derivatives^23^. The effectors tested were p65-HSF1 (PH)^28^ and VP64-p65-Rta (VPR)^16^ as activators for CRISPRa, and KRAB (K)^13^ and KRAB-MeCP2 (KM)^31^ as repressors for CRISPRi. We compared sgRNA designs carrying either two internal MS2 or PP7 hairpins (MS2 tested for CRISPRa and CRISPRi; PP7 tested for CRISPRa only) or a 3’-terminal com hairpin (tested for CRISPRi only). Capture sequences were placed internally or terminally depending on hairpin position (Figure S1B).

The experimental workflow is summarized in Figure 1C. Genome-edited hiPSCs were selected with puromycin, monoclonally expanded, and screened by genotyping and flow cytometry to verify at least single-copy, on-target integration and sfGFP reporter expression. In exploratory experiments, we evaluated various designs and obtained two main conclusions. First, PH and VPR produced comparable CRISPRa activity (Figure S1C); however, VPR-expressing hiPSCs progressively lost typical undifferentiated morphology upon prolonged culture, prompting us to prioritize PH for subsequent optimization. Second, K and KM showed configuration-dependent activity: K was effective when recruited by 3’-terminal com, whereas KM performed only with internal MS2; in this context, K yielded more robust CRISPRi overall (Figure S1D).

Based on these results, we refined the platform in a second set of experiments focused on PH and K to drive activation and repression of *MYOD1* and *TFRC*, respectively, using validated sgRNAs^14,19^ compared to a non-targeting control (NTC). For this refined clone set, we prioritized homozygous *AAVS1*-edited clones with homogeneous sfGFP expression (>90%; representative flow cytometry in Figure S1E; summary in Table S1), resulting in comparable sfGFP mean fluorescence intensity (MFI) across the tested configurations (Figure S1F). Selected clones were then analyzed by RT-qPCR for the gene being targeted for repression or activation (Figure 1D). For CRISPRa, we observed activation of the myogenic master regulator *MYOD1* using both MS2::MCP and PP7::PCP, though the former proved marginally stronger, in contrast to what was observed in other cell types^26^. Specifically, MS2-cs2::MCP combination provided the strongest and most consistent activation, reaching up to ∼75fold induction over NTC (although absolute *MYOD1* expression remained modest at ∼0.2% of housekeeping gene levels), and was chosen for subsequent experiments. For CRISPRi, only com-cs1::Com achieved near-complete silencing of *TFRC* (∼98% knockdown); therefore, this combination was chosen for the subsequent experiments. Unexpectedly, we did not observe significant repression in the absence of cs1. Additional RT-qPCRs for dCas9 and the effectors demonstrated that the observed differences in CRISPRa/i efficiency did not correlate with changes in transgene expression; moreover, none of the conditions disrupted the expression of pluripotency markers (Figure S1G).

In all, we identified two effective strategies for CRISPRa and CRISPRi in hiPSCs that rely on modified sgRNAs harboring distinct recruitment strategies, streamlining their combined use for bidirectional gene regulation in hiPSCs. A schematic representation of the selected components is provided in Figure 1E.

### An isogenic tetracycline-inducible CRISPR hiPSC platform

Having identified efficient recruiting strategies, we next focused on inducibility to support perturbations that would otherwise compromise hiPSC maintenance. Because constitutive expression of dCas9 and the PH or K RBP-effector modules had no overt effects on hiPSC fitness, we implemented temporal control at the level of the sgRNA, thereby keeping the protein components constant while gating targeting and effector recruitment. We previously established a tetracycline (Tet)responsive Pol III promoter to drive inducible sgRNA expression for CRISPR/Cas9 nuclease-mediated gene knockout in hPSCs, and showed that this architecture retains robust inducibility across numerous differentiated lineages^23^. This strategy avoids the risk of differentiation-associated silencing of doxycycline-inducible dCas9 expression in hiPSC derivatives^24^. To enable analogous Tet-inducible regulation for CRISPRa/i, we generated a hiPSC line constitutively expressing a codon-optimized version of the Tet repressor (OPTtetR)^23^ and dCas9-2A-mCherry from two established GSHs: *hROSA26*^32^ and *CLYBL*^33^, respectively (Figure 2A). We used separate loci to minimize promoter interference, and leveraged the CAG promoter for both transgenes to support stable, stoichiometrically balanced expression of the core chassis across differentiation. As our ultimate goal was to employ the resulting cell line to perform screens for gene programs able to induce a myogenic fate in hiPSCs, we employed a WTC11 hiPSC line previously engineered to contain a fluorescent reporter knock in at a key sarcomeric locus (*TTN*-mEGFP)^34^.

**Figure 2.**
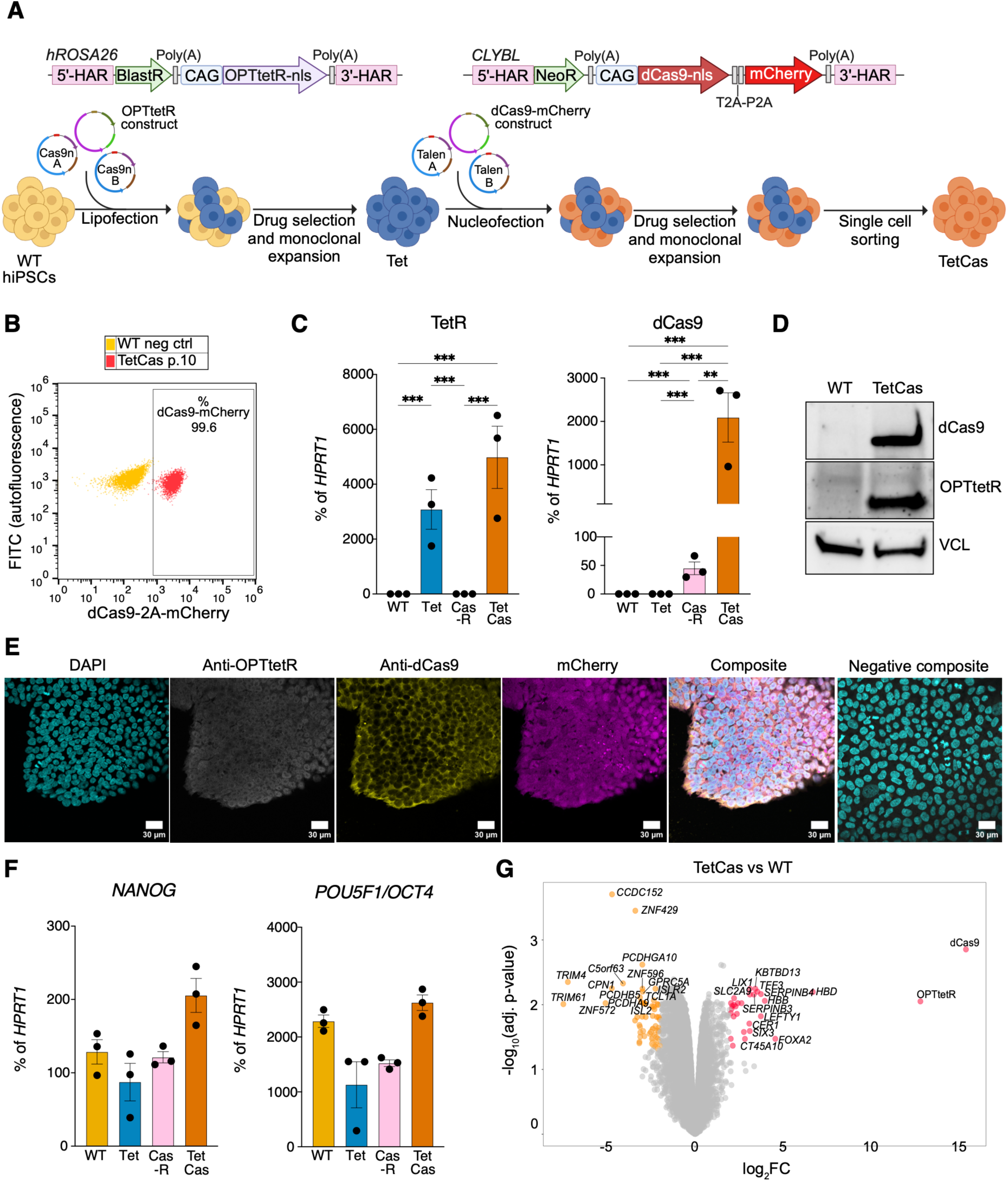
Generation of hiPSCs for tetracycline-inducible CRISPRa/i (A) Two-step engineering workflow. *TTN*-mEGFP reporter WTC11 hiPSCs were targeted with codon-optimized tetracycline repressor (OPTtetR) at *hROSA26* (Tet hiPSCs), then re-targeted with dCas9-2A-mCherry at *CLYBL* and subcloned to obtain TetCas hiPSCs. Cas9n, *S. pyogenes* D10A Cas9 nickase; TALEN, transcription activator-like effector nuclease; BlastR, blasticidin S deaminase; NeoR, neomycin phosphotransferase II; other abbreviations as in Figure 1C. Constructs not shown to scale. (B) Flow cytometry quantification of dCas9-2A-mCherry expression in TetCas hiPSCs after 10 passages (p.10). (C) RT-qPCR quantification of transgene mRNA expression in TetCas hiPSCs compared with Tet, Cas9-R, and WT hiPSCs. N = 3 cultures; one-way ANOVA followed by Holm–Sidak’s multiple-comparison test for all pairwise comparisons. ** = p < 0.01; *** = p < 0.001. (D) Western blot analysis of transgene protein expression in TetCas hiPSCs. VCL, vinculin loading control. (E) Confocal microscopy of multiplexed transgene immunostaining in TetCas hiPSCs. Scale bars, 30 µm. (F) As in panel C, but assessing pluripotency marker expression. p > 0.05 for all comparisons. (G) Volcano plot of differential gene expression between TetCas and WT hiPSCs by bulk RNA-seq. N = 3 cultures. Genes down- and upregulated in TetCas hiPSCs are shown in orange and red, respectively.

For the first editing step, we transfected WTC11 *TTN*-mEGFP hiPSCs with the targeting vector encoding a blasticidin resistance gene trap and OPTtetR, along with CRISPR/Cas9n constructs to facilitate *hROSA26* editing. Genome edited cells were selected with blasticidin S and monoclonal lines were screened by genomic PCR to verify site-specific targeting (Fig-ure S2A). We selected one clone expressing the transgene in homozygosity and showcasing no additional random integrations. We named this cell line “Tet”.

For the second editing step, we retargeted Tet cells by electroporation with a targeting vector encoding a neomycin resistance gene trap and dCas9-2A-mCherry, together with *CLYBL*-specific transcription activator-like effector nucleases (TALENs). We selected edited hiPSCs using geneticin before clonal expansion, genotyping, and flow cytometry to determine mCherry expression (Figure S2B and summary in Table S2). Since only a subset of genotype-positive hiPSCs showed mCherry expression, we further subcloned mCherry-positive cells by fluorescence-activated cell sorting (FACS; Figure S2C). Genotyping of the subclones confirmed that all were heterozygous without random integrations (Figure S2D).

Through this procedure, we isolated a line homogeneously expressing dCas9-2A-mCherry at high levels for at least ten low-density passages in culture (Figure 2B). We validated the expression of the two transgenes at RNA level by RT-qPCR (Figure 2C), and at protein level by western blot (Figure 2D) and immunofluorescence (Figure 2E). This line, hereafter “TetCas”, showed ∼500-fold higher expression of dCas9 compared to Cas-R hiPSCs, showcasing the advantage of relying on GSH for stable transgene expression. Importantly, sequential genome editing and constitutive expression of OPTtetR and dCas9-2A-mCherry did not alter expression of pluripotency markers *NANOG* and *POU5F1/OCT4* at the transcript (Figure 2F) and protein level (Figure S2E). To assess potential subtle, non-specific effects, we performed bulk RNA-seq comparing TetCas cells and parental WTC11 *TTN*-mEGFP (each analyzed in three sequential subcultures). Differential expression analysis identified a limited number of genes (29 up- and 70 down-regulated more than 2-fold; adjusted p < 0.05), indicating no major transcriptional differences (Figure 2G). Gene Set Enrichment Analysis (GSEA) detected no enrichment of the “response to DNA damage” or “signal transduction by p53 class mediator” gene sets, mitigating concerns of a potential genotoxic stress signature (Figure S2F). Consistent with this, comparative genomic hybridization (CGH) confirmed that TetCas hiPSCs retained genome-wide copy-number profiles indistinguishable from parental WTC11 *TTN*-mEGFP line (Figure S2G). These analyses collectively validated a *TTN* reporter hiPSC line stably expressing all transgenes required to implement inducible multimodal CRISPRa and CRISPRi.

### Inducible CRISPRa and CRISPRi in hiPSCs and differentiated derivatives

We employed TetCas cells to perform inducible CRISPRi and CRISPRa in hiPSCs and their differentiated derivatives. Using the same workflow as above, we engineered the *AAVS1* locus with cassettes encoding for an inducible sgRNA and constitutively expressed RBP-effector-2A-sfGFP. Based on the optimization results, we used the MS2-cs2::MCP-PH combination for CRISPRa and com-cs1::Com-K for CRISPRi. To express Tet-inducible sgRNAs, we leveraged a U6 promoter containing a single Tet operator (U6-TO)^35^, which is repressed by OPTtetR in standard culture media and rapidly activated upon Tet addition. We selected U6-TO over our previously established system based on an analogous Tet-inducible H1 promoter because U6 drives higher sgRNA levels and a more precisely defined 5’ transcript start^36,37^, improving on-target activity and reducing the likelihood of off-target effects. Monoclonal lines were screened by *AAVS1* genotyping and flow cytometry to confirm co-expression of dCas9-2A-mCherry and the effector-2A-sfGFP cassette (summary in Table S3); clones were then treated with Tet prior to RT-qPCR analysis. A schematic of the constructs is shown in Figure S3A.

In the inducible CRISPRi setting, repression of *TFRC* was weaker than in the configuration-matched constitutive CRISPRi system (Figure 3A). Nevertheless, we achieved effective inducible CRISPRi of other targets, such as the pluripotency genes *NANOG* and *SOX2* (up to ∼87% and ∼64% knockdown, respectively; Figure 3B). Thus, the suboptimal performance of inducible CRISPRi at *TFRC* may reflect iron-metabolism-linked regulation of this transcript in an acute *versus* chronic perturbation setting (see DISCUSSION), rather than an intrinsic limitation of Tet-inducible sgRNA expression. Of note, comparison of *SOX2* inducible CRISPRi lines in the absence of Tet to NTC controls demonstrated that the system was not significantly leaky (Figure 3B), addressing a key concern for inducible CRISPRi implementations^19^.

**Figure 3.**
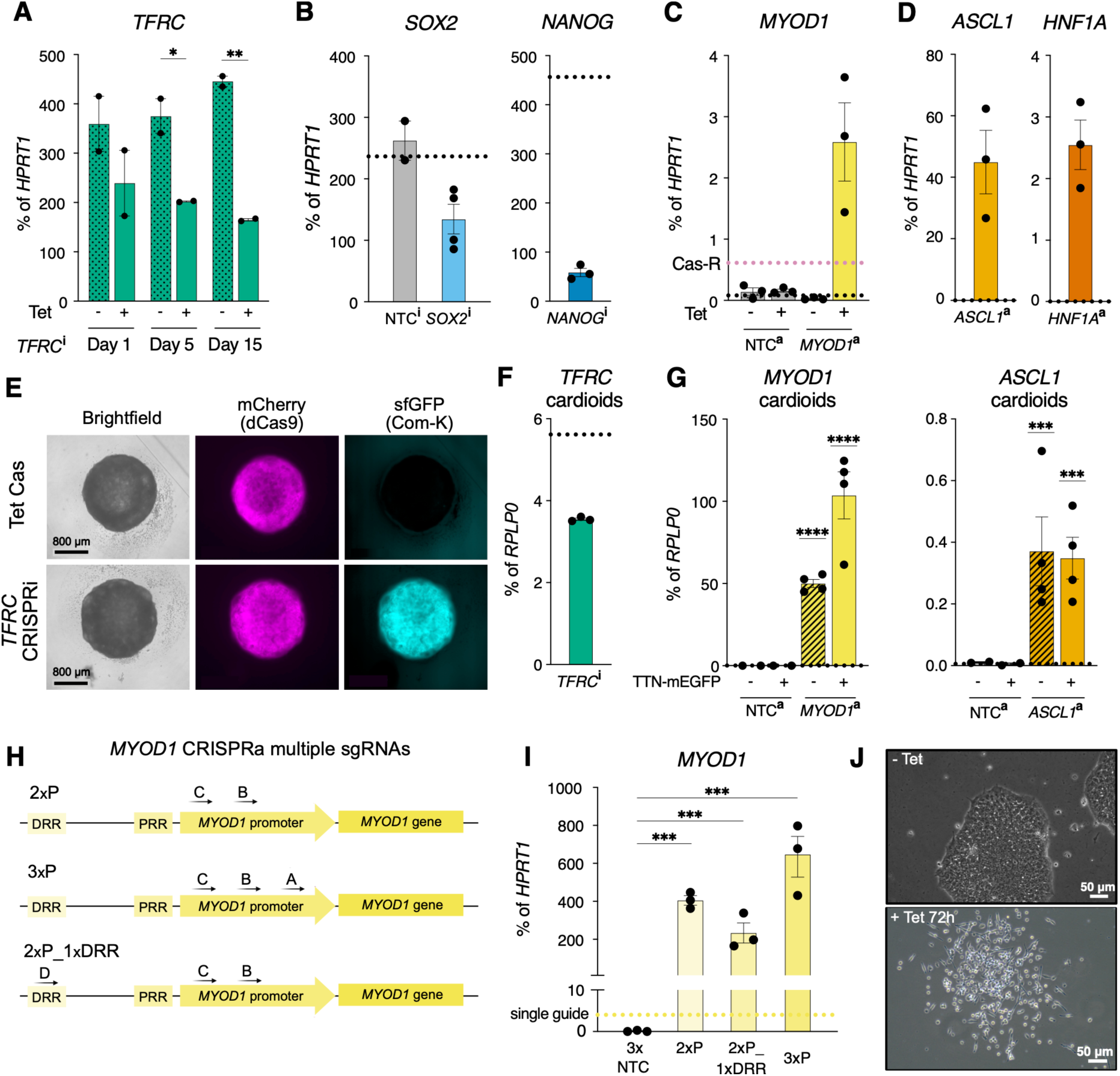
Validation of inducible CRISPRa/i in hiPSCs and cardiac organoids (A) RT-qPCR validation of inducible *TFRC* CRISPRi in a clonal TetCas hiPSC line re-targeted at *AAVS1* with the CRISPRi architecture optimized in Figure 1. The clone was homozygously edited and lacked random integrations. Cells were treated with Tet (+) for the indicated times and compared with matched untreated controls (-). N = 2 cultures; two-way ANOVA with Tet treatment and time as factors followed by Holm-Sidak’s multiple-comparison test for - *versus* +. Here and throughout the figure, * = p < 0.05; ** = p < 0.01; *** = p < 0.001. (B) As in panel A, but for inducible *SOX2* and *NANOG* CRISPRi after 3 days of Tet treatment, with Tet-treated NTC controls shown where applicable. N = 2–4 clones; one-sample two-tailed t-tests against the indicated pooled Tet-untreated reference values (black dotted lines). (C) As in panel A, but for inducible *MYOD1* CRISPRa after 5 days of Tet treatment, compared with matched untreated and NTC controls. The pink dotted line indicates the benchmark level of *MYOD1* expression achieved by architecture-matched constitutive CRISPRa in Cas-R hiPSCs (Figure 1D). N = 3 clones; one-sample two-tailed t-test against the indicated pooled Tet-untreated *MYOD1* CRISPRa reference value (black dotted line). (D) As in panel A, but for inducible *ASCL1* and *HNF1A* CRISPRa after 3 days of Tet treatment. N = 3 clones; one-sample two-tailed t-tests against the indicated pooled Tet-untreated reference values (black dotted lines). (E) Representative widefield fluorescence images of day 7.5 cardiac organoids (cardioids) generated from *TFRC* CRISPRi hiPSCs, showing maintenance of dCas9-2A-mCherry and Com-K-2A-sfGFP. Scale bar, 800 µm. (F) As in panel A, but for inducible *TFRC* CRISPRi in day 7.5 cardiac organoids Tet-treated for 3 days. N = 3 clones; one-sample two-tailed t-test against the indicated Tet-treated NTC reference value (black dotted line). (G) As in panel A, but for inducible *MYOD1* and *ASCL1* CRISPRa, together with NTC controls, in day 7.5 cardiac organoids. Organoids were Tet-treated for 24 h and sorted into TTN-mEGFP^+^ cardiomyocytes and TTN-mEGFP^-^non-cardiomyocytes before analysis. N = 2 differentiations using 1-2 clones per target; one-sample two-tailed t-tests against the indicated pooled Tet-untreated, TTN-mEGFP-matched reference values (black dotted lines). (H) Schematic of the *MYOD1* regulatory regions and sgRNA combinations tested for multiplex CRISPRa. Individual sgRNAs are indicated as A–D. 2xP, two promoter guides; 3xP, three promoter guides; 2xP_1xDRR, two promoter guides plus one distal regulatory region (DRR) guide; PRR, proximal regulatory region, not tested. (I) RT-qPCR validation of multiplex inducible *MYOD1* CRISPRa after 24 h of Tet treatment. The three-guide NTC control (3xNTC) and the single-sgRNA benchmark from panel C (yellow dotted line) are shown for comparison. N = 3 clone pools; one-way ANOVA followed by Holm–Sidak’s multiple-comparison test for all pairwise comparisons. (J) Representative phase-contrast images of hiPSCs carrying the 3xP inducible *MYOD1* CRISPRa system after 72 h of Tet treatment, compared to Tet-untreated controls. Scale bar, 50 µm.

Inducible CRISPRa activated *MYOD1* to ∼3% of housekeeping gene levels, a ∼6-fold improvement compared to the prior configuration-matched non-inducible system. This suggests that, at least for CRISPRa, higher and more homogeneous dCas9 expression alone substantially improves efficiency. Despite this, the system remained tightly controlled by Tet (∼115-fold induction over untreated controls), demonstrating leak-proof control (Figure 3C). RT-qPCR analyses confirmed robust expression of the system components and maintenance of pluripotency before Tet treatment (Figures S3B–S3C). We further validated CRISPRa on genes specific to other germ layers: *ASCL1* (a neuronal master regulator) and *HNF1A* (an important driver of liver differentiation), achieving robust, tightly-gated activation of both targets with only moderate clonal variability (Figure 3D).

To test CRISPRa/i functionality after differentiation, we validated CRISPRa/i activity in hiPSC-derived cardiac organoids recapitulating the developing left ventricle^38^ (Figure S3D). In the inducible *TFRC* CRISPRi line, dCas9-2A-mCherry and Com-K-2A-sfGFP remained homogeneously expressed at day 7.5 of differentiation, when the TTN-mEGFP reporter was not yet detectable by fluorescence microscopy (Figure 3E). Tet treatment at this stage resulted in moderate *TFRC* silencing, consistent with the hiPSCs data (Figure 3F).

For CRISPRa experiments, we replaced sfGFP with TagBFP to enable stringent analysis of hiPSC-derived cardiomyocytes (hiPSC-CMs). We sorted TTN-mEGFP^+^ hiPSC-CMs from TTN-mEGFP^-^ cells (Figure S3E) and observed strong, tightly controlled activation in both fractions for a lineage-adjacent gene, *MYOD1*, and a cross germ layer marker, *ASCL1* (Figure 3G). Interestingly, the relative CRISPRa efficiencies for these targets were inverted compared with hiPSCs, consistent with differentiation-associated chromatin remodeling in hiPSC-CMs that facilitates activation of mesodermal programs while imposing additional epigenetic barriers to activating neuroectodermal factors. As above, RT-qPCR further confirmed stable expression of circuit components across clones (Figure S3F).

To overcome epigenetic barriers that limit activation of *MYOD1* in hiPSCs, we combined multiple sgRNAs targeting distinct regulatory elements of *MYOD1*, including the promoter and the distal regulatory region (DRR; Figure 3H). We generated gene-edited pools in biological triplicates and confirmed by flow cytometry and RT-qPCR that comparable fractions of cells expressed MCP-PH-2A-TagBFP and the other circuit components (Figures S3G–S3H). RT-qPCR after 24 hours of Tet treatment revealed >100-fold increase in *MYOD1* activation relative to the single-guide setting, reaching ∼600% of housekeeping levels with three promoter-targeting guides (Figure 3I). Importantly, baseline *MYOD1* remained near zero in the absence of Tet, indicating tight control even with multiplex sgRNA cassettes. Consistent with strong *MYOD1* induction, hiPSCs treated with Tet for 72 h dispersed from colonies and adopted an elongated morphology despite being maintained in pluripotency media (Figure 3J).

Together, these results demonstrate that inducible CRISPRa/i is effective in both hiPSCs and a key mesodermal differentiated derivative, and that combination of multiple sgRNAs can enhance the magnitude of activation while preserving stringent off-state control.

### CIRI: orthogonal inducible CRISPRa and CRISPRi to rewire cell-fate programs

Having optimized inducible CRISPRa and CRISPRi modules individually, we combined them to develop CIRI (**C**ombinatorial **I**nducible C**R**ISPR in **I**PSCs), a platform enabling simultaneous gene activation and silencing to implement multi-node regulatory logic. We prototyped CIRI for transcription factor-driven “forward programming” of hiPSCs, and hypothesized that activating a lineage master regulator while repressing core pluripotency factors would increase fate transition efficiency by reducing competition between gene regulatory networks.

We first tested this concept in neuronal forward programming, a well-established model of transcription factor-driven fate conversion. To generate neuronal CIRI hiPSCs, we conucleo-fected TetCas hiPSCs with two *AAVS1* targeting vectors: one encoding three Tet-inducible sgRNAs against *POU5F1/OCT4*, *SOX2*, and *NANOG* (OSN) together with the CRISPRi effector, and one encoding an *ASCL1*-targeting CRISPRa cassette (*ASCL1*^a^-*OSN*^i^ CIRI). We generated matched controls carrying only the *ASCL1* CRISPRa cassette, as well as NTC matched controls (Table S4). Compound heterozygous *AAVS1* integrants were selected by dual antibiotic selection, followed by clonal isolation and validation as above. Clones were cultured for 14 days in neuronal-permissive media in the presence of Tet. Under these conditions, *ASCL1* was robustly induced in both *ASCL1* CRISPRa-only and *ASCL1*^a^-*OSN*^i^ CIRI, while CIRI further reduced the already low expression of *POU5F1/OCT4* and *NANOG* at this stage (but not *SOX2*, see the DISCUSSION; Figure S4A). Consistent with lineage priming, neuronal markers *OLIG3* and *MAP2* were induced, with higher levels in CIRI than CRISPRa alone (Figure S4A).

Encouraged by this proof-of-principle, we applied CIRI more thoroughly to skeletal muscle forward programming by combining the CRISPRi cassette against pluripotency factors with the *MYOD1* CRISPRa cassette with three promoter-targeting sgRNAs (*MYOD1*^a^-*OSN*^i^; Figure 4A). To avoid overlap with TTN-mEGFP, we omitted a fluorescent reporter for the CRISPRi effector while retaining a TagBFP reporter for CRISPRa effector. We obtained three compound heterozygous myogenic CIRI clones and matched heterozygous *MYOD1* CRISPRa-only controls, stably expressing dCas9-2A-mCherry and the CRISPRa effector (Figures S4B–S4D; Table S4). Upon Tet treatment in pluripotency media, RT-qPCR showed that pluripotency genes began to decrease by 24 hours in both *MYOD1*^a^-*OSN*^i^ CIRI and *MYOD1* CRISPRa-only conditions, consistent with robust *MYOD1* activation promoting exit from pluripotency. By 72 hours, however, CIRI achieved stronger and near-complete silencing of OSN, whereas we observed residual expression or even partial rebound in the *MYOD1* CRISPRa-only condition (Figure 4B). *MYOD1* induction at 24 hours was robust in both settings, and strengthened at 72 hours, consistent with autoregulation^39^ (Figure 4B). Muscle markers *MEF2C* and *MYOG* were activated only at 72 hours, consistent with the known temporal dynamics of early myogenic differentiation^40^ (Figure 4C). At this time point, transcriptional remodeling was accompanied by morphological changes (Figure 4D). As in prior experiments, no CRISPRa/i activity was detected in the absence of Tet, and NTC controls showed no changes in pluripotency factors or *MYOD1*. RT-qPCR confirmed stable expression of both genetic circuits (Figure S4E).

**Figure 4.**
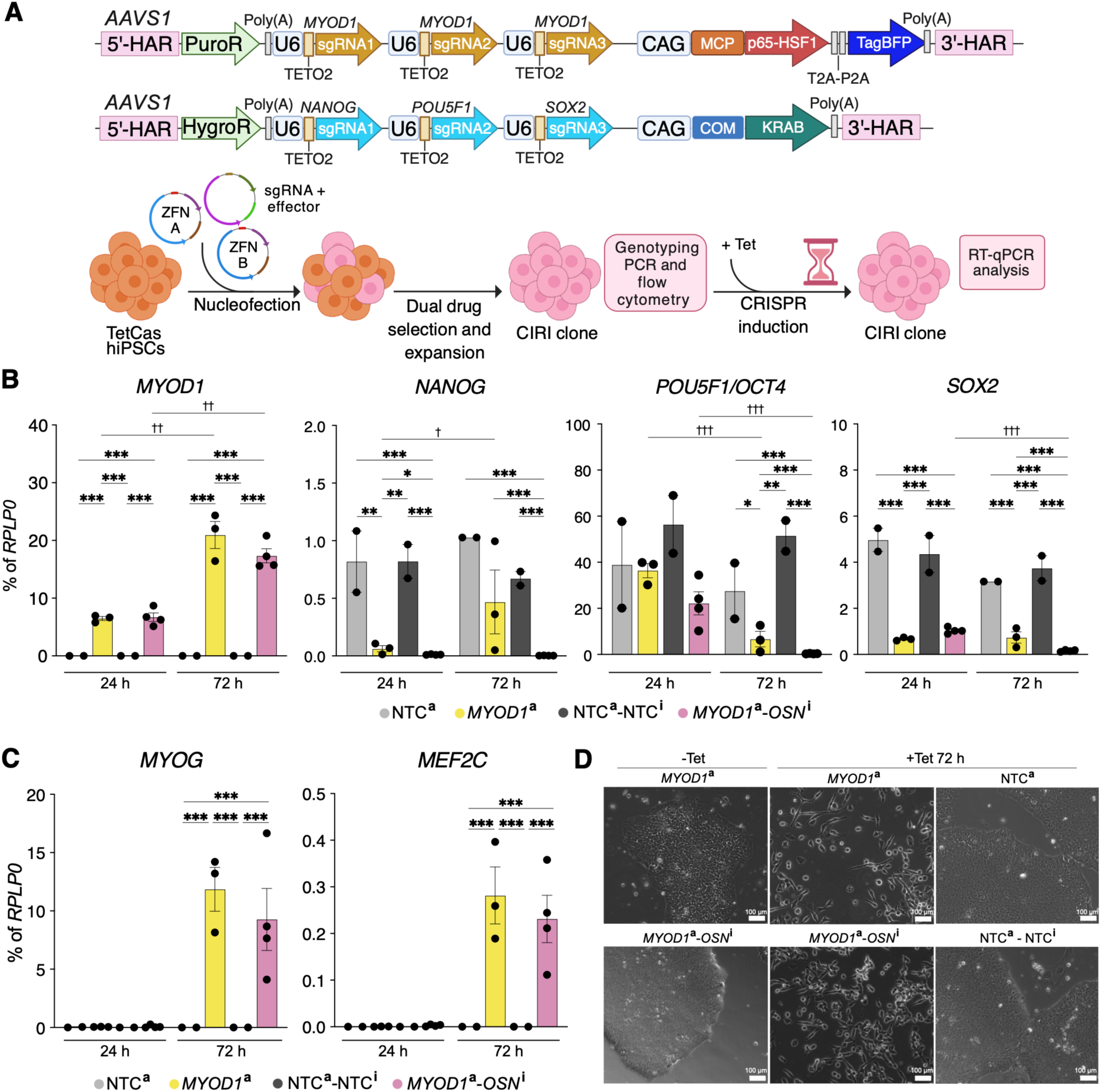
Implementation of combinatorial CRISPRa and CRISPRi in hiPSCs (A) Workflow for simultaneous *MYOD1* activation and pluripotency-factor repression. TetCas hiPSCs were bi-allelically edited at *AAVS1* with inducible CRISPRa and CRISPRi cassettes, each encoding one effector and three sgRNAs: three promoter guides for *MYOD1* activation (3xP; Figure 3H) and individual guides against *POU5F1/OCT4*, *NANOG*, and *SOX2* (OSN) for CRISPRi, hereafter *MYOD1*^a^-*OSN*^i^ CIRI. HygroR, hygromycin B phosphotransferase; other abbreviations as in Figure 1C. Construct not shown to scale. (B) RT-qPCR analysis of *MYOD1* activation and OSN silencing in *MYOD1*^a^-*OSN*^i^ CIRI clones and *MYOD1* CRISPRa-only (*MYOD1*^a^) clones, compared with matched NTC clones. Cells were Tet-treated for 24 h or 72 h. N = 2–4 clones; two-way ANOVA with cell line configuration and time as factors, followed by Holm–Sidak’s multiple-comparison test between configurations within the same time point (asterisks), and between time points within the same configuration (daggers). *, † p < 0.05; **, †† p < 0.01; ***, ††† p < 0.001; p > 0.05 not reported. (C) As in panel B, but assessing downstream activation of myogenic markers *MYOG* and *MEF2C*. Statistical annotations are shown only for the 72 h comparisons, because 24 h values were at or near the detection limit. (D) Representative phase-contrast images of cells from panels B-C after 72 h of Tet treatment. Scale bar, 100 µm.

Overall, these results show that CIRI enables orthogonal, bidirectional regulation of gene sets, and support its use for hiPSC forward programming by coupling activation of lineage master regulators with coordinated repression of the core pluripotency network.

### CIRI drives canalized myogenic forward programming from endogenous loci

Having established that CIRI enables efficient gene expression control, we next tested whether it supports *bona fide* forward programming by culturing *MYOD1*^a^-*OSN*^i^ CIRI hiPSCs in myogenic-permissive media (Figure 5A). *MYOD1*^a^-*OSN*^i^ CIRI hiPSCs began morphological remodeling within 24 hours of Tet treatment and adopted an elongated morphology consistent with myoblast identity by day 4; by day 7, they appeared as partially multinucleated myotubes (Figure S5A). A subset of *MYOD1* CRISPRa cells also elongated, but a sizeable fraction retained compact colony-like regions that appeared as dark foci in phase contrast, with abundance varying by clone and resembling the morphology observed in NTC controls hiPSCs (Figure S5B). At day 7, both CIRI and CRISPRa conditions expressed the TTN-mEGFP reporter, MYOD1, and the sarcomeric muscle marker *α*-actinin-2 (ACTN2), whereas these were absent from the com-pact cluster regions observed in CRISPRa-only cultures and from NTC controls (Figure 5B and Figure S5C).

**Figure 5.**
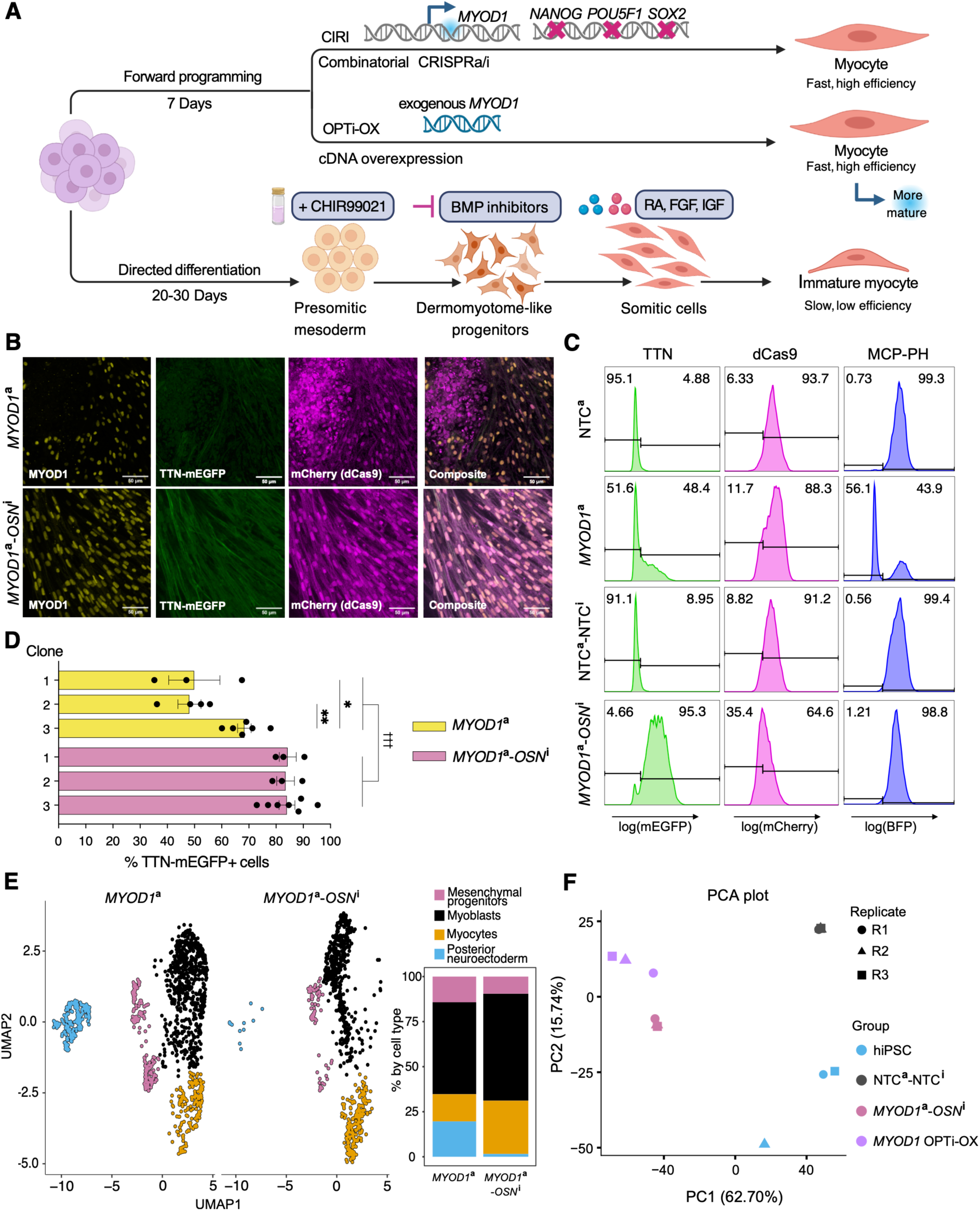
Lineage-canalized myogenic forward programming by simultaneous *MYOD1* activation and repression of pluripotency factors (A) Schematic of 7-day myogenic forward programming with *MYOD1*^a^-*OSN*^i^ CIRI, compared with benchmark *MYOD1* OPTi-OX forward programming and conventional small molecule-directed differentiation^41^. (B) Confocal microscopy after 7 days of myogenic differentiation in representative *MYOD1*^a^-*OSN*^i^ CIRI and *MYOD1*^a^ CRISPRa-only clones. Scale bars, 30 µm. (C) Representative flow cytometry analysis of the conditions shown in panel B, compared with matched NTC clones. BFP quantifies maintenance of the MCP-PH CRISPRa effector. Histograms are scaled to mode. (D) Flow cytometry quantification of TTN-mEGFP^+^ cells from panel C for three clones per condition. N = 3–7 differentiation; two-way ANOVA with cell-line configuration and integration-matched clone class as factors, followed by Holm–Sidak’s multiple-comparison test for the indicated comparisons. Asterisks indicate clone-to-clone differences within the same configuration; daggers indicate the overall comparison between *MYOD1*^a^-*OSN*^i^ CIRI and *MYOD1*^a^ CRISPRa-only cells. * p < 0.05; ** p < 0.01; ††† p < 0.001; p > 0.05 not reported. (E) Single-cell RNA-seq analysis of the experiment described in panels B-D. UMAP, Uniform Manifold Approximation and Projection; clusters were annotated based on marker-gene expression (Figure S5D). Bar plots show the relative proportion of each cell state. (F) Bulk RNA-seq comparison of the conditions described in panels B-D with time-matched *MYOD1* OPTi-OX programming and untreated hiPSCs. N = 3 differentiations. PCA, Principal Component Analysis.

Flow cytometry showed that *MYOD1*^a^-*OSN*^i^ CIRI drove more homogeneous and reproducible differentiation than *MYOD1* CRISPRa-only (>80% *versus* ∼50% of TTN-mEGFP^+^ cells, reproducible across three clones and 3-7 differentiation runs; Figures 5C–5D). This correlated with more homogeneous expression of dCas9-2A-mCherry and MCP-PH-2A-TagBFP in CIRI, while in CRISPRa >50% of cells were negative for the effector (corresponding to TTN-mEGFP^-^cells; Figure 5C). This suggests that differentiation conditions imposed a selective advantage for a small subset of more proliferative cells that poorly expressed the CRISPRa transgene.

scRNA-seq supported and extended these findings. *MYOD1* CRISPRa alone produced a mis-differentiated population with posterior neuroectoderm-like features (∼20%; *SOX2*^+^, *CDX2*^+^, and *PTPRZ1*)^+^, alongside partially differentiated mesenchymal/fibroadipogenic progenitors (∼14%; *PLIN2*^+^ and *UGCG*^+^) and proliferating myoblasts (∼51%; *PAX7*^+^ and *CCND2*^+^), with only ∼15% of myocytes (*MYH7B*^+^, *MYLK*^+^, and *MB*^+^). In contrast, *MYOD1*^a^-*OSN*^i^ CIRI approximately doubled the myocyte fraction and reduced mis-differentiated cells by >10-fold (Figure 5E and Figure S5D).

We next benchmarked *MYOD1*^a^-*OSN*^i^ CIRI against *MYOD1* OPTi-OX, which uses doxycycline (Dox)-inducible cDNA overexpression from a dual safe-harbor isogenic system^42^, and has emerged as a widely used platform for inducible skeletal muscle forward programming^43,44^. We cultured CIRI and OPTi-OX under matched culture conditions with Tet/Dox, and compared them by bulk RNA-seq, also including a CIRI NTC control and undifferentiated hiPSCs. Principal component analysis (PCA) separated negative controls from differentiated samples and placed CIRI close to OPTi-OX along the principal axis of variance associated with myogenic *versus* pluripotency programs (Figure 5F and Figure S5E). Biological replicates were highly concordant, with CIRI triplicates essentially overlapping. Nevertheless, GSEA (Figures S5F–S5G) and differential expression (Figure S5H) indicated that OPTi-OX reached a more mature transcriptional state, consistent with a stronger overexpression of transgenic *MYOD1*.

Together, these results establish CIRI as a robust approach for rapid forward programming from endogenous loci, achieving high efficiency and reproducibility comparable to optimized cDNA overexpression while reducing mis-differentiation compared to CRISPRa alone through coordinated activation of lineage drivers and repression of pluripotency networks.

### Combinatorial pooled CIRI screens nominate synergistic myogenic regulators

Encouraged by the efficiency of CIRI in myogenic forward programming, and motivated by the differentiation gap *versus* OPTi-OX, we next used *MYOD1*^a^-*OSN*^i^ CIRI as basis for scRNA-seq-based screening to identify additional differentiation drivers and/or roadblocks whose combinatorial modulation could accelerate myogenic conversion, enhance myocyte maturation, and/or reduce reliance on exogenous components in the myogenic medium (FGF2, CHIR99021, and retinoic acid - RA; Figure 6A). In particular, removal of RA substantially reduced differentiation yield in the baseline condition, motivating the search for complementary drivers (Figure 6B). Thus, we designed a combinatorial screening in which CRISPRa activates *MYOD1* in combination with an additional myogenic transcription factor^45–47^ (*MEF2C*, *MYF5*, *MYF6*, *MYOG*, or *PAX7*) or epigenetic modifier^48,49^ (*SMARCD3* or *KDM6B*), while CRISPRi represses OSN plus a candidate reprogramming roadblock^50–52^ (*ATF7IP*, *ZNF207*, *CTCF*, or *YBX1*) or *MYOD1* antagonist^53^ (*ID3*, the predominant ID family member expressed in hiPSCs; Figure S6A). Including matched NTC controls, the design totaled 48 combinatorial perturbations.

**Figure 6.**
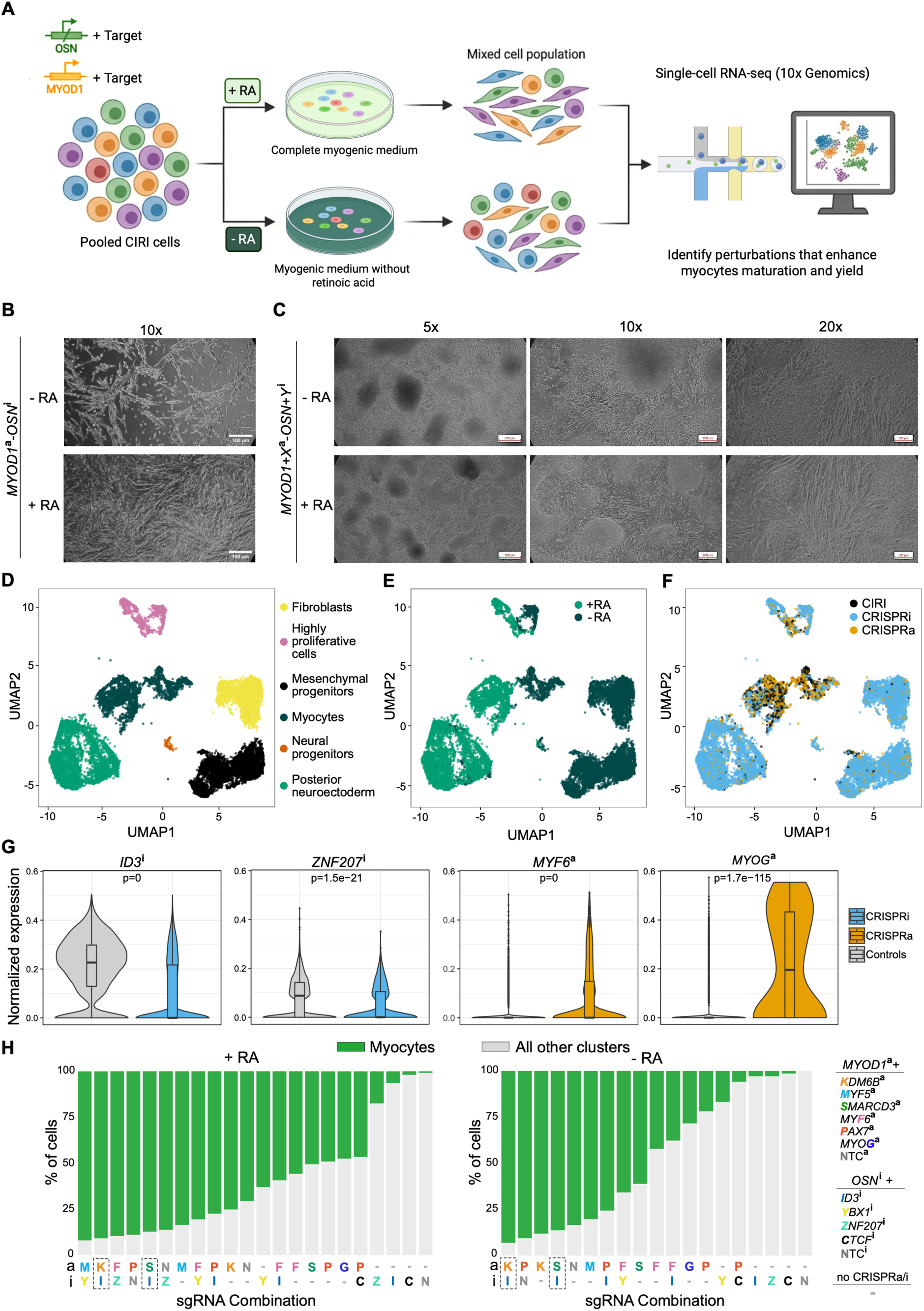
Identification of myogenic enablers and roadblocks by pooled combinatorial CRISPRa/i single-cell RNA-seq screening (A) Schematic of the single-cell RNA-seq screening workflow to identify additional factors that enhance *MYOD1*^a^-*OSN*^i^ CIRI-mediated myogenic programming when activated or repressed. Each perturbation adds one CRISPRa target and one CRISPRi target to the baseline configuration to promote maturation and/or reduce dependence on exogenous retinoic acid (RA). (B) Phase-contrast images of baseline 7 day differentiation with and without RA. Scale bar, 100 µm. (C) As in B, but for dual-guide pooled CIRI cells. Scale bars, 500 µm (5x), 200 µm (10x), and 100 µm (20x). (D) UMAP of the dual-guide pooled CIRI screen after 7 days of myogenic differentiation with and without RA; clusters were annotated based on marker-gene expression (Figure S6C). (E) As in D, colored by RA-containing and RA-free differentiation conditions. (F) As in D, colored by assigned perturbation modality: CRISPRa-only, CRISPRi-only, or both (CIRI). (G) Violin plots of normalized expression for representative CRISPRa/i targets in control and perturbed cells. (H) Quantification of the fraction of assigned cells adopting a myocyte fate for each combinatorial perturbation. Only the added factors on top of the *MYOD1*^a^-*OSN*^i^ baseline are reported. Dashed boxes indicate selected hits.

To interrogate these candidates systematically, we generated pooled CRISPRa and CRISPRi libraries in either single-guide or dual-guide configurations. For CRISPRa, we designed four sgRNAs for each target promoter, and cloned them either individually (single-guide, top two sgRNAs based on efficiency predictions) or in paired combinations (dual-guide) together with two sgRNAs targeting the *MYOD1* promoter. We followed a similar approach for CRISPRi libraries, except that sgRNAs were designed to bind closely to the transcription start sites and were cloned together with sgRNAs targeting OSN. Because repeated Tet-inducible Pol III promoters triggered plasmid recombination during pooled cloning, library construction required iterative optimization; the technical troubleshooting is reported in the Supplemental Text.

We engineered TetCas hiPSCs with pooled CRISPRa and CRISPRi libraries and selected for compound heterozygous *AAVS1* targeting with one cassette per type. We then differentiated edited pools for 7 days under myogenic conditions in the presence or absence of RA (RA+ or RA-). Several cells adopted an elongated, myocyte-like morphology, whereas others formed compact cluster-like regions, consistent with incomplete or diverted fate progression. Gross morphology was similar between RA+ and RA-conditions, and overall yield in RA-appeared improved relative to the baseline RA-experiment without screening perturbations (Figure 6C and Figure S6B).

After 7 days, we profiled perturbation-induced transcriptional states by 10X Genomics micro-fluidics-based 5*^′^* scRNA-seq, capturing both types of expressed sgRNAs using a CRISPR-specific primer alongside transcriptomes. We then leveraged an adapted version of our recently published *catcheR* bioinformatic workflow^54^ to assign perturbations to individual cells and classify them as CRISPRa-only, CRISPRi-only, or CIRI. Across all conditions, 77.4–84.2% of cells contained at least one protospacer, with a mean of 1,122–2,601 CRISPR-associated unique molecular identifiers (UMI) per cell. CRISPRi-only cells represented 51.3-83.2% of assigned cells, consistent with preferential recovery or expansion relative to CIRI cells, which accounted for 4-39%. Full QC statistics are reported in Table S5.

In the dual-guide experiments, unsupervised clustering identified discrete populations corresponding to expected differentiation trajectories. In both RA+ and RA-conditions, a prominent cluster displayed a robust myogenic signature, indicating that myogenic differentiation occurred even independently of RA (Figure 6D and Figure S6C). In fact, RA+ myogenic cells expressed higher levels of the progenitor marker *PAX7* and lower levels of *MYOG* and other definitive myogenic genes, suggesting that while RA supported survival during myogenic differentiation, it limited progression toward a more differentiated state (Figure S6C). This was a reproducible finding, also observed for the single-guide screening (Figure S6D). Besides this, RA+ cultures additionally segregated into clusters enriched for neuroectodermal markers, whereas RA-cultures also produced mesenchymal progenitor– or fibroblast-like states (Figure 6E). Stratification by perturbation class showed that CIRI and CRISPRa-only cells were enriched in the myogenic cluster, while, as expected, CRISPRi-only cells were overrepresented in the non-myogenic trajectories (Figure 6F). Target gene expression changes upon perturbation validated the efficiency of CRISPRa and CRISPRi (Figure 6G).

Integrating perturbation identity with transcriptional outcomes enabled the identification of candidate combinatorial interactions that promote myogenic specification. Combined, dual-guide activation of *KDM6B*, also known as *JMJD3*, together with repression of *ID3* emerged as the strongest reproducible CIRI hit, enriching for cells adopting a myogenic transcriptional state in both RA+ and RA-conditions (Figure 6H). Additional high-ranking candidates included activation of *SMARCD3*, also known as *BAF60C*, again in combination with repression of *ID3*, ranked highly in both conditions (Figure 6H). Hits from the single-guide experiments were less striking, but *KDM6B* and *SMARCD3* activation emerged as lead candidates, in this case combined with *ZNF207* repression (Figures S6E–S6H).

Together, these screens establish CIRI as a pooled single-cell screening framework that can identify candidate activators, repressors, and synergistic combinations to improve forward programming outcomes and rationally refine differentiation conditions.

### Combinatorial hits enable retinoic acid-free myogenic forward programming

We next investigated whether screening-nominated regulators could improve CIRI-driven myogenic forward programming under RA-free conditions, testing whether endogenous regulatory rewiring could replace such a pro-survival exogenous patterning cue associated with a more progenitor-like state, and achieve a maturation level comparable to, or exceeding, *MYOD1* OPTi-OX. To this end, we generated clonal, *AAVS1*-edited TetCas lines in which the baseline *MYOD1*^a^-*OSN*^i^ CIRI configuration was expanded with dual sgRNAs for additional repression of *ID3* and activation of either *SMARCD3*/*BAF60C* (*MS*^a^-*OSNI*^i^) or *KDM6B*/*JMJD3* (*MK*^a^-*OSNI*^i^) (Table S6). Molecular karyotyping confirmed that these lines, baseline *MYOD1*^a^-*OSN*^i^, and benchmark *MYOD1* OPTi-OX hiPSCs lacked newly acquired large-scale copy-number alterations relative to parental WTC11 *TTN*-mEGFP hiPSCs (Figure S7A).

We first validated sgRNA functionality by RT-qPCR in hiPSC clones maintained under pluripotency conditions and exposed to Tet for 72 hours. As expected, Tet treatment induced robust upregulation of *SMARCD3* or *KDM6B* in the corresponding lines, together with repression of *ID3* (Figure 7B).

**Figure 7.**
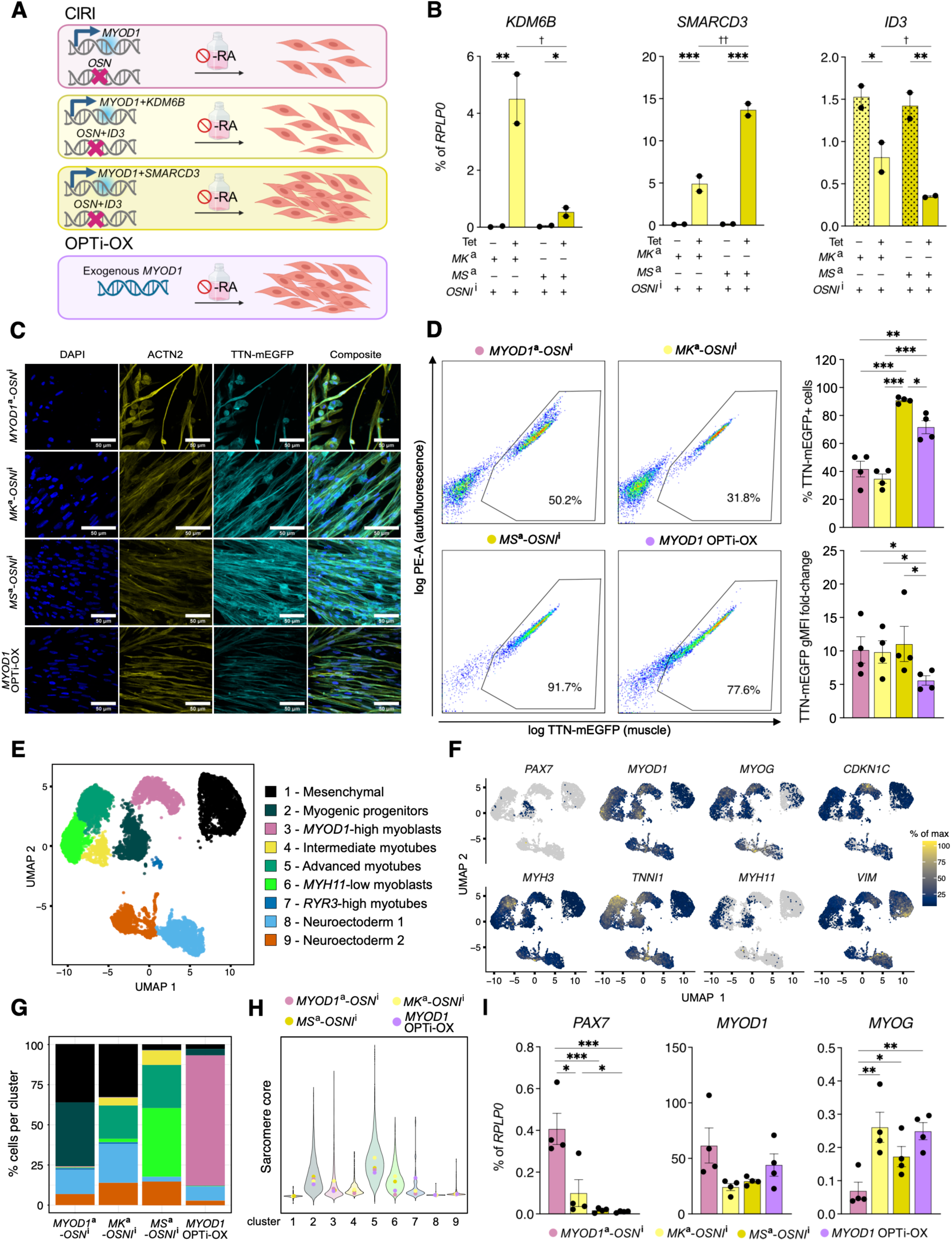
Enhanced and retinoic acid-independent myogenic programming by adding *SMARCD3* or *KDM6B* activation and *ID3* repression (A) Experimental approach. Baseline *MYOD1*^a^-*OSN*^i^ CIRI was expanded with *SMARCD3* or *KDM6B* activation and *ID3* repression, generating *MS*^a^-*OSNI*^i^ and *MK*^a^-*OSNI*^i^ lines, respectively. Cells were differentiated for 7 days in retinoic acid-free (-RA) myogenic medium and compared with baseline CIRI and *MYOD1* OPTi-OX. (B) RT-qPCR validation of sgRNA functionality after 72 h of Tet treatment. N = 2 cultures; two-way ANOVA with cell line and Tet-treatment as factors, followed by Holm–Sidak’s multiple-comparison test between treatment in the same cell line (asterisks), and between cell line within the same treatment (daggers). Here and throughout the figure: *, † p < 0.05; **, †† p < 0.01; ***, ††† p < 0.001; p > 0.05 not reported. (C) Confocal microscopy after 7 days of -RA myogenic differentiation. Scale bars, 50 µm. (D) Flow cytometry quantification of myogenic differentiation after 7 days in -RA myogenic medium, measured as the percentage of TTN-mEGFP^+^ cells and TTN-mEGFP geometric mean fluorescence intensity (gMFI) fold-change *versus* hiPSCs. N = 4 differentiations; here and in panel I, statistical analysis by repeated-measures one-way ANOVA followed by Holm–Sidak’s multiple-comparison test of all pairwise comparisons. (E) UMAP of single-cell RNA-seq data from the experiment described in panels C-D. Cells are colored by transcriptional state, annotated based on marker-gene expression. (F) Feature plots showing representative marker genes used to annotate the transcriptional states in panel E. (G) Relative proportion of each transcriptional state across conditions. (H) Violin plots of myogenic module scores across transcriptional states and conditions. (I) RT-qPCR analysis of selected myogenic markers from the experiment described in panel D.

Both expanded CIRI configurations improved RA-independent myogenic forward programming relative to baseline *MYOD1*^a^-*OSN*^i^ CIRI. In RA-free myogenic medium, the baseline configuration showed poor survival and limited differentiation, whereas both *MS*^a^-*OSNI*^i^ and *MK*^a^-*OSNI*^i^ supported substantially better cell retention and earlier appearance of elongated, myotube-like structures. These structures were already evident by day 5 post-induction, approximately 48 hours before comparable morphology became apparent in *MYOD1* OPTi-OX cultures under the same RA-free conditions (Figure S7B).

Immunofluorescence at day 7 confirmed the presence of elongated, multinucleated TTN-mEGFP^+^/ACTN2^+^ cells with well-defined sarcomeric organization in both expanded CIRI conditions (Figure 7C). Flow-cytometric quantification further showed that *MS*^a^-*OSNI*^i^ produced the highest fraction of TTN-mEGFP^+^ cells among the tested RA-free conditions (>90%), while TTN-mEGFP geometric mean fluorescence intensity was broadly comparable, indicating that the major effect was on the fraction of cells entering the sarcomeric program rather than on per-cell TTN reporter intensity (Figure 7D).

To resolve the cell-state composition underlying these phenotypes, we performed scRNA-seq at day 7 across baseline CIRI, the two expanded CIRI configurations, and *MYOD1* OPTi-OX, all differentiated without RA. Unsupervised clustering identified nine transcriptionally distinct populations (Figures 7E–7F and Figures S7C–S7D). To avoid over-annotation from individual marker genes, we ordered the myogenic populations along a conservative progenitor-to-myotube continuum: *PAX7*^+^ myogenic progenitors, also enriched for *CA3*; *MYOD1*^hi^ differentiating myoblast-like cells, also enriched for *CDKN1C* and *CRABP1*; intermediate differentiating myocytes/myotubes expressing sarcomeric genes such as *TNNI1*; and a more advanced sarcomeric myotube state enriched for *TNNI1* and *MYH3*. In addition, we identified a *SMARCD3*-enriched myogenic population with low-level expression of smooth-muscle-associated markers, including *MYH11*, and a small calcium-handling and excitation-contraction-associated myotube subset marked by *RYR3*. Non-myogenic states comprised a cycling mesenchymal-like population enriched for *VIM* and *MKI67*, together with two residual non-myogenic clusters showing neuroectoderm transcriptional features (Figure 7F and Figure S7D).

Condition-stratified analysis revealed distinct mechanisms of improvement across the two expanded CIRI configurations (Figure 7G). Baseline *MYOD1*^a^-*OSN*^i^ CIRI performed worst, showing an enrichment in a *PAX7*^+^ progenitor-like state rather than in more differentiated myocyte or myotube populations. Both expanded CIRI configurations shifted cells away from this progenitor-like state. However, they did so in different ways: *MS*^a^-*OSNI*^i^ produced the broadest and most homogeneous myogenic output across the lineage continuum, whereas *MK*^a^-*OSNI*^i^ showed stronger enrichment of its myogenic cells in the more mature sarcomeric myotube cluster. *MYOD1* OPTi-OX showed the most homogeneous outcome overall, with most cells concentrated in a single post-mitotic myogenic cluster enriched for the cell-cycle inhibitor *CDKN1C*. However, this homogeneity did not correspond to the highest maturation state in the scRNA-seq analysis: the most mature sarcomeric cluster was more strongly represented within the expanded CIRI conditions. Consistently, sarcomere core module-score analysis indicated improved myogenic maturation in the expanded CIRI configurations relative to baseline CIRI, with the highest-scoring cells found within the mature myotube state rather than in the dominant OPTi-OX cluster (Figure 7H).

RT-qPCR across biological replicates supported the broad progenitor-to-myogenic shift observed by scRNA-seq: *PAX7* was highest in baseline CIRI, whereas *MYOG* was lowest in this condition (Figure 7I). Additional RT-qPCR further confirmed higher *CDKN1C* expression in OPTi-OX and higher *CA3* expression in baseline CIRI, consistent with the cluster-level enrichment of these markers (Figure S7E). By contrast, bulk RT-qPCR did not detect significant differences in the sarcomeric maturation markers *MYH3* and *TNNI1*, nor did it resolve the low-level, subset-restricted *MYH11* signal observed in the *SMARCD3*-activated condition, underscoring the value of single-cell analysis for distinguishing differences in cell-state composition from uniform changes in population-average gene expression (Figure S7E).

Together, these results validate the predictive value of the pooled CIRI screen and show that simultaneous repression of *ID3* with activation of either *SMARCD3* or *KDM6B* improves RA-independent myogenic forward programming.

## DISCUSSION

In this study, we establish CIRI as an inducible, isogenic, and modular CRISPR platform for simultaneous gene activation and repression in hiPSC models. By combining a single constitutive dCas9 chassis with Tet-inducible sgRNAs carrying orthogonal RNA scaffolds, CIRI enables multimodal control of endogenous loci while minimizing transgene burden and preserving tight temporal regulation across pluripotency and differentiation. In practice, this architecture proved sufficient not only for robust CRISPRa and CRISPRi in hiPSCs and their derivatives, but also for canalized forward programming and iterative single-cell refinement of lineage engineering. More broadly, our results define CIRI as both a synthetic-biology platform and an experimental framework for dissecting how coordinated transcriptional perturbations can rewire cell identity.

CIRI occupies a still under-served region of the CRISPRa/i design space. Several groups have described bidirectional or combinatorial CRISPRa/CRISPRi systems based on orthogonal dCas9 proteins for parallel activation and repression of distinct loci^55–60^, most commonly based on direct effector fusions built on *S. pyogenes* and *S. aureus* dCas9 backbones. These studies have provided important proof-of-principle for dual perturbations, epigenetic editing, and paired-gene interaction analyses. However, multiple large protein fusions and multi-component architectures can become cumbersome when scaled or transferred to fragile cellular systems, such as hiPSCs. Conversely, aptamer-based combinatorial CRISPR regulation was elegantly demonstrated in HEK293-derived settings, showing that RNA scaffolds can support orthogonal recruitment of activation and repression modules from a shared dCas9 backbone^25,26^. However, these systems lacked the inducible, safe-harbor, and pluripotent-stem-cell-compatible design features that are essential when perturbing fate determinants whose constitutive activity would rapidly destabilize the starting state. CIRI was developed specifically to bridge these two lines of work: the scalability and composability of scaffold-based regulation, and the temporal control and genetic stability required for deployment in hiPSCs.

A second conceptual advance of this study is the demonstration that inducible CRISPRa and CRISPRi can be used synergistically to guide rapid and highly efficient lineage conversion by simultaneously activating master regulators and extinguishing the pluripotency network. A related concept was previously explored by Black and colleagues, who showed in PSCs that coordinated activation and repression can improve neuronal specification^56^. However, that framework relied on orthogonal CRISPR systems and did not address the specific challenge of building an inducible, isogenic platform tailored to pluripotent cells. Our data with CIRI now show that in hiPSCs, activating lineage-defining factors while concurrently repressing core pluripotency genes can overcome the residual heterogeneity of CRISPRa-only programming, resulting in a more homogeneous and more reproducible transition than activation of the lineage determinant alone. Notably, in the myogenic setting this strategy supported conversion efficiencies approaching completeness and, after iterative refinement, yielded outcomes comparable to optimized cDNA-based programming while exceeding it in selected readouts. We therefore view dual-mode endogenous programming, rather than single-factor activation alone, as a potentially generalizable design principle for stem-cell engineering.

Our myogenic screens further yielded a biological insight that is, in our view, as important as the platform itself. Additional myogenic transcription factors contributed comparatively little over *MYOD1*, whereas repression of *ID3* and activation of the chromatin regulators *SMARCD3/-BAF60C* and *KDM6B/JMJD3* markedly improved both the efficiency and transcriptional maturity of RA-independent programming. This pattern suggests that, in hiPSCs, *MYOD1* already provides the dominant upstream instructive signal, consistent with its long-recognized ability to drive myogenic conversion even when expressed alone^53^. In contrast, the main residual bottleneck may reside in chromatin competence rather than in a lack of additional myogenic transcription-factor input. In this model, *ID3* repression could relieve a classical brake on bHLH-driven differentiation, whereas *SMARCD3/BAF60C* and *KDM6B/JMJD3* could facilitate productive engagement of the myogenic program by enabling locus accessibility and removal of repressive chromatin barriers^49,61,62^. The convergence of these hits on chromatin regulation leads us to hypothesize that in hiPSCs, endogenous *MYOD1* activation is largely sufficient to specify the direction of fate change, but efficient execution of that program depends on permissive chromatin architecture.

The importance of chromatin regulators over added transcription factors may also help explain why *MYOD1*-driven programming can be powerful yet still imperfect. *MYOD1* and *ASCL1* are evolutionarily related bHLH pioneer factors that bind surprisingly similar regions in fibroblasts, even though they normally induce radically different lineages; importantly, *MYOD1* can display pro-neuronal activity through promiscuous occupancy of neuronal loci^63^. This is highly relevant to our data, because *MYOD1* CRISPRa alone produced off-target and incompletely differentiated states, whereas CIRI and especially the *SMARCD3/BAF60C*-enhanced version more efficiently channeled cells into a mature myogenic trajectory. One interpretation is that such epigenetic modifiers do not simply amplify *MYOD1* output, but instead help focus *MYOD1* onto productive myogenic loci and away from alternative accessible programs. However, this focusing effect may not be entirely skeletal-muscle-specific: because *SMARCD3/BAF60C* functions across muscle regulatory contexts, including cardiac and smooth muscle, its activation may also permit low-level engagement of adjacent contractile programs, consistent with the *MYH11*-low subset in the *SMARCD3*-enhanced condition.

Recent work suggests that some master regulators shape cell identity not only through pioneer DNA binding activity, but also through three-dimensional (3D) genome organization. *MYOD1* has been shown to promote long-range intra- and inter-chromosomal contact hubs associated with muscle identity^64^, while *POU5F1/OCT4* can reorganize TAD structure during somatic cell reprogramming^65^. We recently broadened the RNA factory hypothesis to propose that co-regulated loci assemble into specialized inter-chromosomal domains that coordinate transcriptional and post-transcriptional control^66^. Systematic computational analyses suggest that such inter-chromosomal networks may be widespread and biologically interpretable, and could eventually be prioritized bioinformatically for functional screening^67,68^. Together, these observations support the idea that factors such as *MYOD1* and *POU5F1/OCT4* can act as “genome architects”, coupling sequence recognition, chromatin opening, and 3D genome organization to enforce cell identity. In this context, a particularly exciting long-term application of CIRI will be to move from serendipitous discovery of such factors to their systematic functional identification and deployment to program cell fate.

While we thoroughly demonstrated CIRI in skeletal muscle forward programming, our pre-liminary neuronal differentiation experiments suggest that inducible, bidirectional transcriptional rewiring may generalize to other lineages. The observation that *ASCL1* activation produced only partial neuronal priming is consistent with the fact that efficient neuronal reprogramming typically requires *ASCL1* in combination with additional transcription factors (e.g., *DLX1/2* or *BRN2/POU3F2* and *MYT1L*) and/or higher dose and timing control of neurogenic programs^69,70^.

In hindsight, our initial CRISPRi cassette also included *SOX2*, which may be counterproductive in a neuroectodermal context because *SOX2* is maintained and functionally required during early neural induction and neuroepithelial specification from human PSCs^71,72^; this provides a plausible explanation for the paradoxical *SOX2* upregulation in neuronal CIRI lines. Together, these results argue that CIRI cassettes should be refined in a lineage-informed manner (e.g., excluding *SOX2* from the repression module for neuronal programs and using combinatorial neurogenic activators), and motivate extending this strategy to other forward-programming paradigms of biomedical relevance, including hepatic (e.g., *HNF4A/HNF6/FOXA3*-driven)^73^ and cardiac (e.g., *GATA4/MEF2C/TBX5*-driven)^74^ gene-regulatory circuits.

Relative to cDNA overexpression or gene knockout, CRISPRa/i offers a distinct set of advantages that are especially valuable in hiPSC systems. Compared with cDNA-based forward programming, CRISPRa acts at endogenous loci, preserving native promoter context, *cis*-regulatory logic, gene dosage constraints, and at least part of the natural temporal architecture of targetgene induction. This makes it particularly attractive for multiplexed perturbations, where introducing many transgenes with the correct stoichiometry rapidly becomes impractical. Compared with nuclease-based knockouts, CRISPRi avoids double-strand breaks and the attendant p53-dependent stress responses that are especially problematic in hPSCs^11,12^. In addition, both modalities are reversible and can be staged in time, which is a major advantage when interrogating developmental windows or when transient perturbation is preferable to constitutive gene disruption^19,23^.

These strengths, however, coexist with intrinsic limitations. Beyond the established locus-and sgRNA design-dependent variability, which continues to improve as prediction tools and empirical datasets mature, there are non-negotiable compromises rooted in endogenous gene regulation. For CRISPRa, activating transcription at a locus does not bypass post-transcriptional control: alternative promoter usage and alternative splicing can yield isoform mixtures that may differ from the desired product, a particularly relevant issue when forcing expression in a lineage where the splicing program is not optimized for the intended cell identity (as in reprogramming). For CRISPRi, incomplete transcriptional silencing can be buffered by post-transcriptional mechanisms that stabilize the remaining transcripts, reducing the apparent knockdown at the RNA level. Our *TFRC* experiments provide a cautionary example of this principle. While constitutive CRISPRi produced strong repression, tet-inducible CRISPRi yielded a smaller apparent effect despite using the same effector logic. *TFRC* mRNA contains multiple iron-responsive elements (IREs) in its 3’UTR and is subject to IRP1/2-mediated stabilization under iron-limiting conditions^75^, such that modest changes in iron availability or cellular stress can shift its steady-state abundance and compress the observable dynamic range of transcriptional repression^76^. Several future developments could further expand the scope of CIRI. On the repression side, it will be worth exploring alternative or more context-adapted silencing architectures, including designs aimed at installing more durable epigenetic repression rather than simply increasing acute knockdown strength^77,78^. At the same time, our own optimization cautions against assuming that nominally stronger repressors will always perform better in hiPSCs: in our hands, KRAB-MeCP2 was less robust than KRAB alone, indicating that recruitability, chromatin context, and overall circuit architecture matter as much as raw repression potency. CIRI is also naturally extensible to three-way implementations. Because PP7-PCP also functioned well in our optimization experiments, MS2, com, and PP7 scaffolds could in principle support tri-modal recruitment schemes that would be especially attractive for probing the causal logic of the histone code and its context dependence, an area now becoming experimentally tractable through increasingly sophisticated modular epigenome-editing platforms^79^.

More broadly, the same scaffold logic could be redeployed beyond transcriptional control. RNA-aptamer recruitment has already powered CRISPR-based live-cell imaging frameworks such as CRISPRainbow^80^, and more recent methods have substantially expanded the spatial and temporal resolution with which genome dynamics can be followed in living cells^81^. Moreover, CRISPR-guided tethering approaches have shown that genomic loci can be repositioned toward defined nuclear compartments, including the nuclear lamina, Cajal bodies, and PML bodies, by recruiting structural proteins such as Emerin, Coilin, or PML^82^. Additionally, programmable manipulation of chromatin looping and higher-order nuclear structure has advanced from reversible single-loop engineering, as in CLOuD9^83^, to more complex CRISPR-guided control of chromatin folding through bioorthogonal strategies^84^. In this context, CIRI provides a conceptual bridge between gene regulation, live imaging, and 3D genome engineering.

Finally, from an implementation standpoint, preinstalled landing-pad architectures should make CIRI substantially easier to scale, especially for large multi-sgRNA payloads. Single-copy landing-pad systems such as STRAIGHT-IN already enable efficient and precise integration of large DNA cargos in human pluripotent stem cells^85^, and more recent dual landing-pad implementations now support simultaneous, allele-specific, single-copy integration of two payloads or gene circuits in hiPSCs^86^. Building CIRI on such chassis would streamline iterative circuit testing, facilitate standardized assembly of larger guide cassettes through Golden Gate workflows, and could ultimately support self-processing sgRNA arrays for the construction of more complex gene-regulatory networks.

Taken together, our findings support a view of cell-fate engineering in which efficient lineage conversion is not achieved simply by turning on a master regulator, but by jointly controlling the transcriptional, chromatin, and architectural context in which that regulator operates. CIRI provides a tractable way to do so in human pluripotent stem cells, while remaining sufficiently modular to support both mechanistic dissection and iterative optimization. We anticipate that this framework will be useful not only for forward programming and screening, but more broadly for discovering the regulatory combinations that define, stabilize, and mature human cell identities. In that sense, CIRI is a foundation for a more programmable, systems-level biology of cell fate.

## Author Contributions

Conceptualization: F.S., A.B.; Methodology: F.S., A.B.; Software and Formal Analysis: M.L.R., S.Bi., E.B.; Data Curation: E.B.; Investigation: F.S., A.R., N.D., M.T., E.R.C., S.Be., H.K., E.B.; Resources: D.C., J.G.Z., A.B.; Visualization: F.S., A.R., E.B.; Writing - Original Draft: F.S., A.B.; Writing - Review & Editing: A.R., E.B.; Supervision: J.G.Z., E.B., A.B.; Funding Acquisition: J.G.Z., A.B.; Project Administration A.B.

## Acknowledgements

We are grateful to Francesca Anselmi for next-generation sequencing support at the NGS UniTo platform, Marta Gai for microscopy support at the OLMA@MBC platform, and Giulia Savorè for experimental assistance. F.S. was supported by a Ph.D. fellowship from PON R&I Azione IV.4 – FSE REACT-EU. Pilot studies were supported by the National Institutes of Health (NIH U54 DK107979-05S1, J.G.Z. and A.B.; NIH R35 GM124773, J.G.Z.). D.C. is funded by Fondazione CDP (Progetto AVANTI Fondazione Telethon) and Italian Ministry of Health (Piano Operativo Salute Traiettoria 3, T3-AN-09, GENOMED). This project has received funding from the European Research Council (ERC) under the European Union’s Horizon Europe research and innovation programme (Grant Agreement No. 101076026, A.B.). Views and opinions expressed are, however, those of the authors only and do not necessarily reflect those of the European Union or the European Research Council Executive Agency; neither the European Union nor the granting authority can be held responsible for them. We acknowledge financial support under the National Recovery and Resilience Plan (NRRP), Mission 4, Component 2, Investment 1.1, Call for tender No. 104 published on 2 February 2022 by the Italian Ministry of University and Research (MUR), funded by the European Union – NextGenerationEU (CUP D53D23005240006; Grant Assignment Decree No. 970, adopted on 30 June 2023 by MUR, D.C and A.B.). This work was also supported by the Giovanni Armenise-Harvard Foundation (Career Development Award 2021, A.B.).

## STAR METHODS

### KEY RESOURCES TABLE

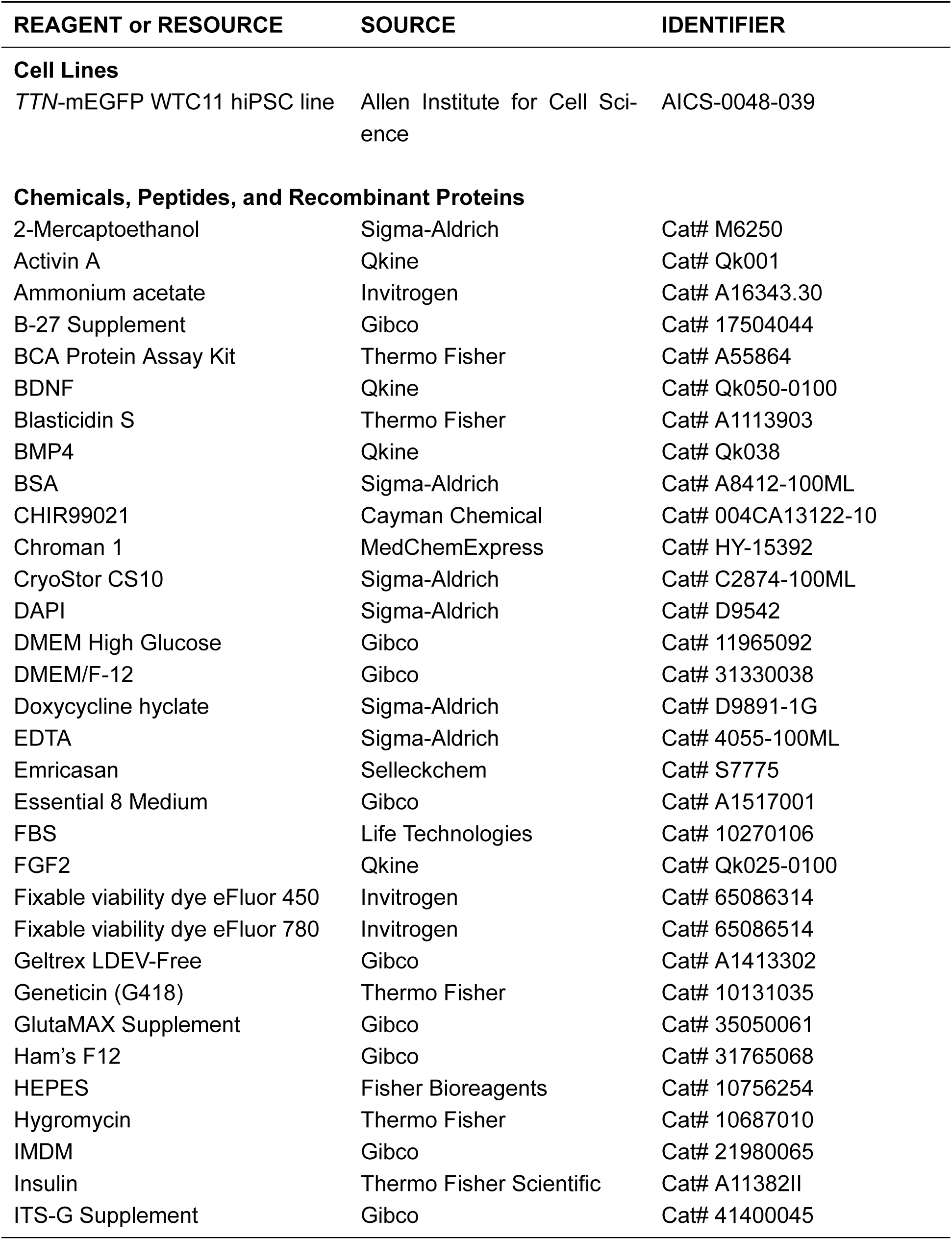
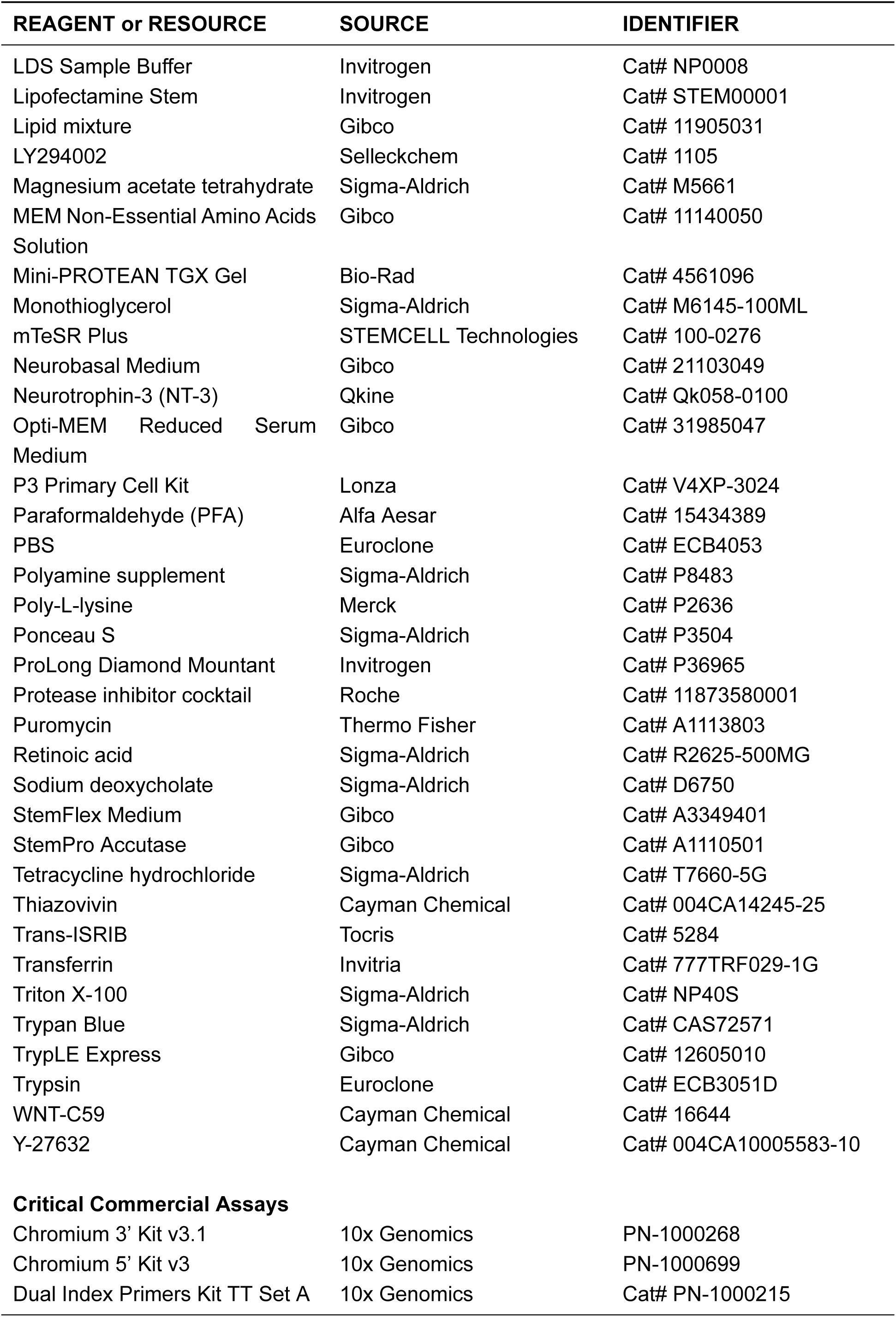
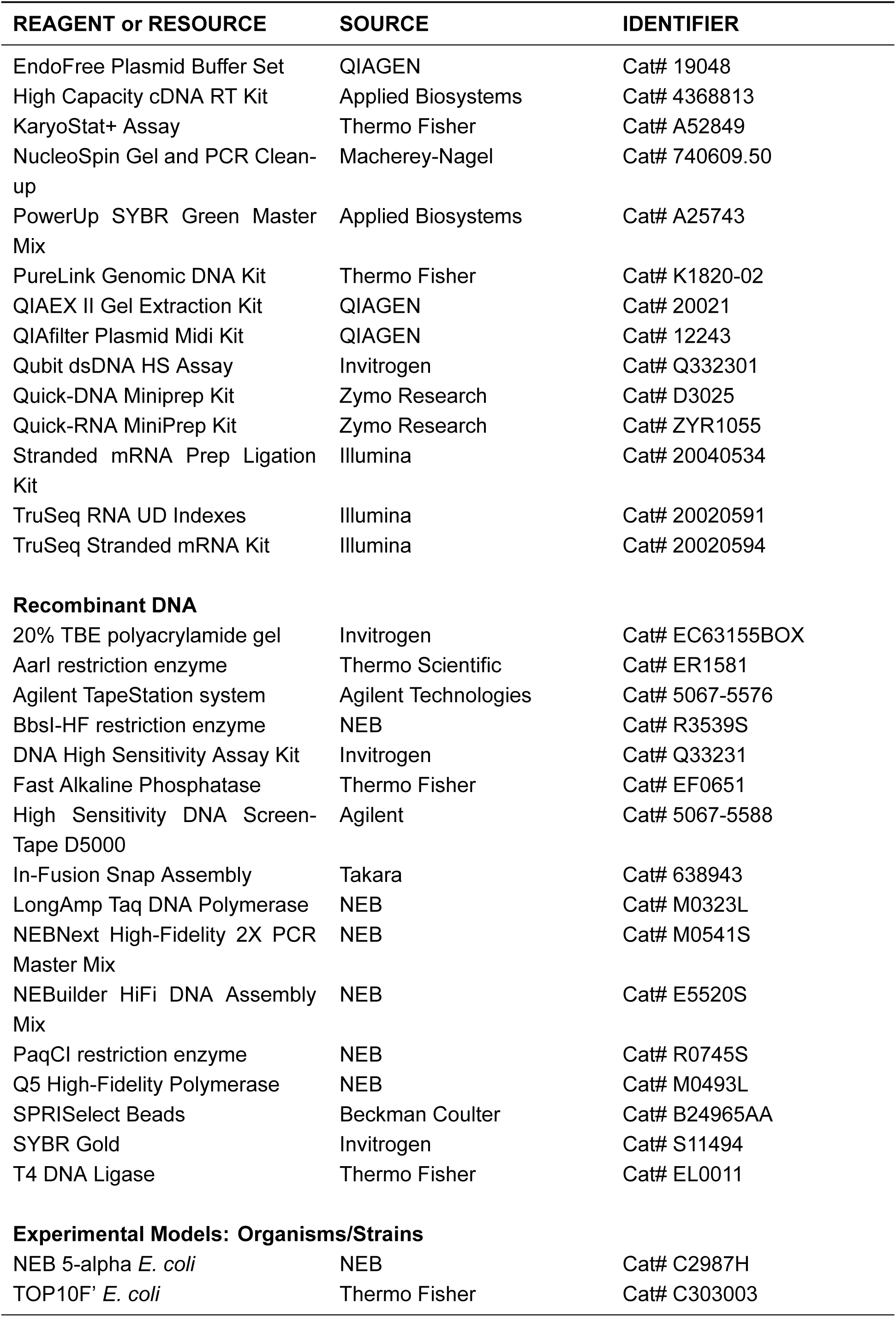
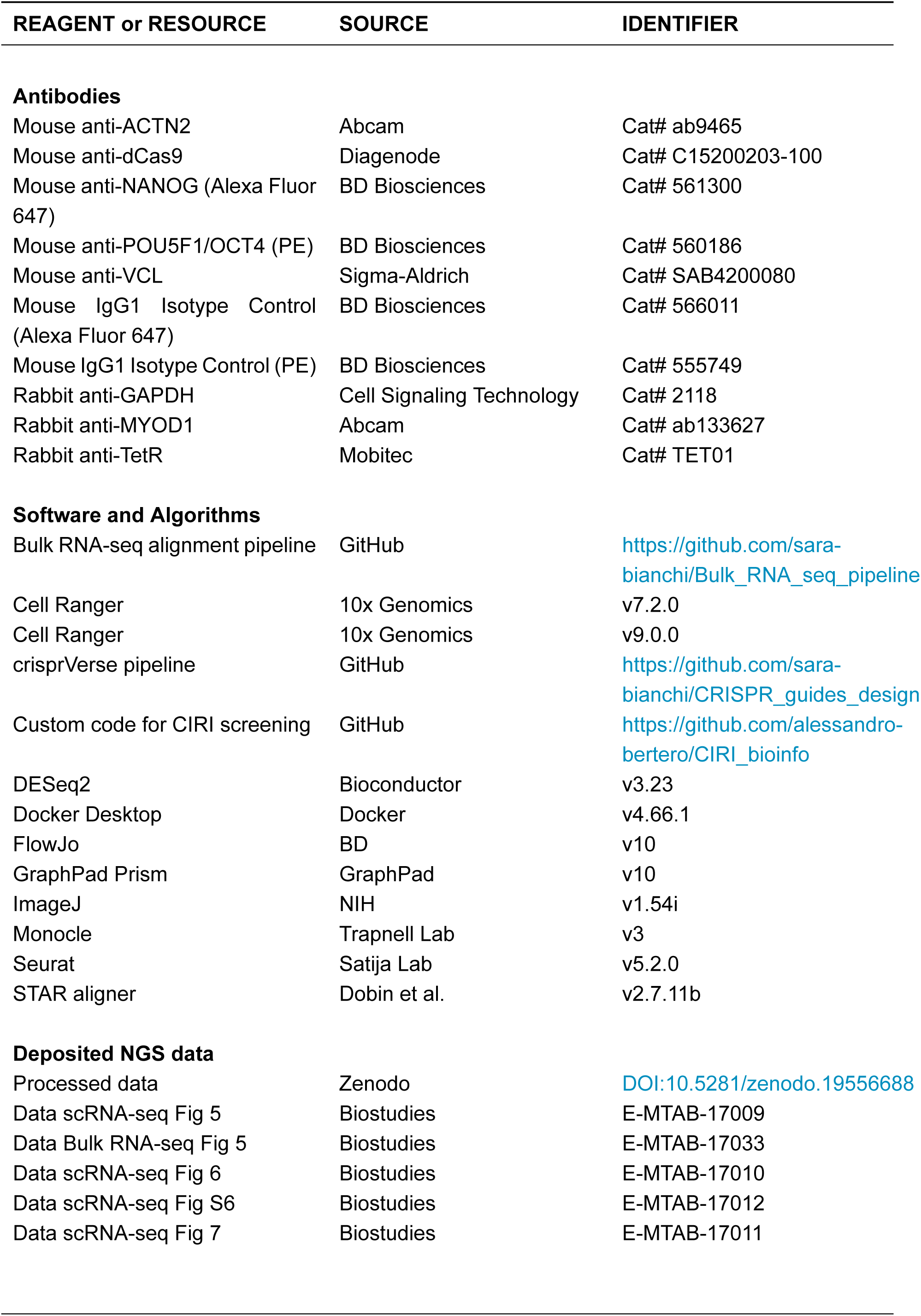
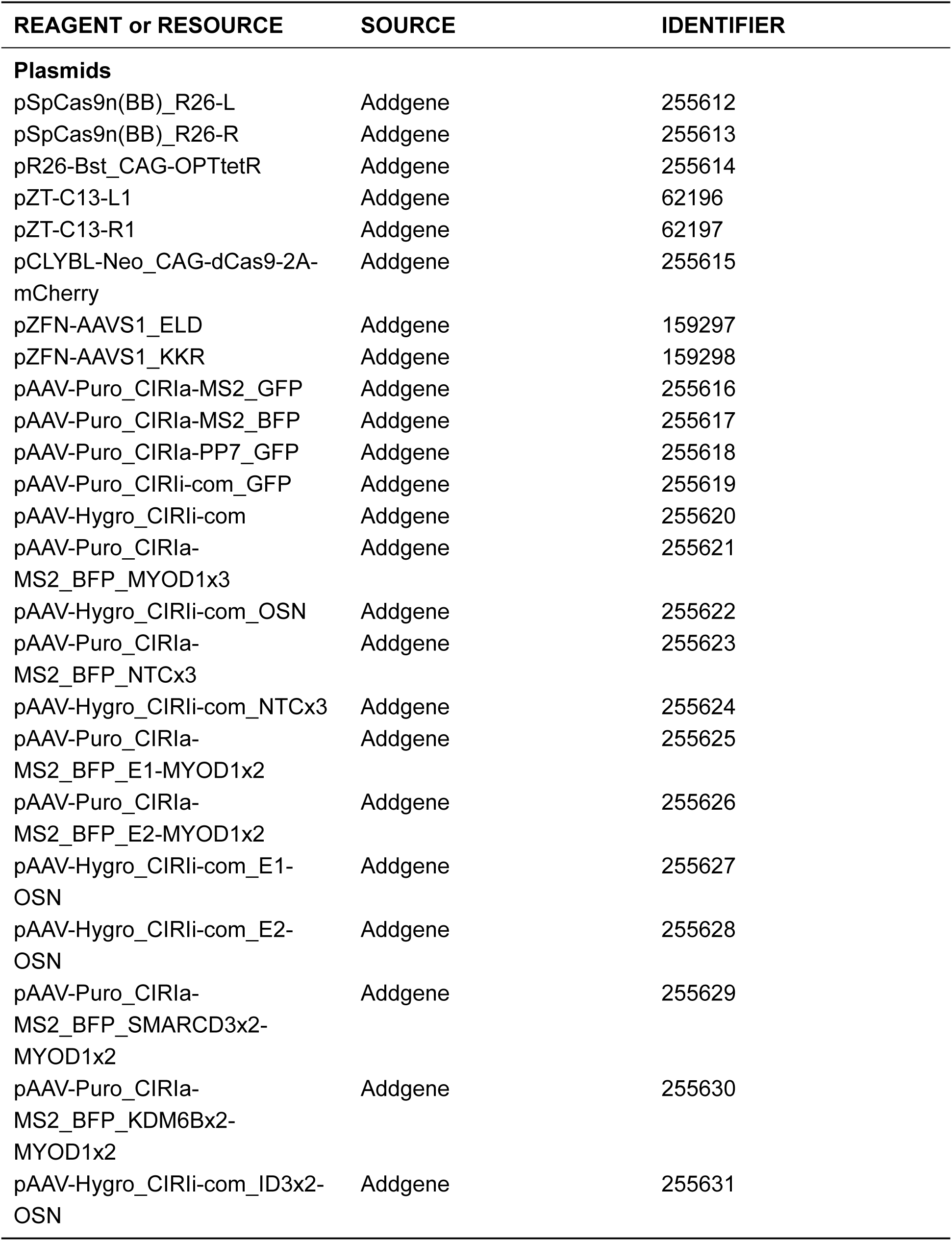

| REAGENT or RESOURCE | SOURCE | IDENTIFIER |
| --- | --- | --- |
| **Cell Lines** |  |  |
| TTN-mEGFP WTC11 hiPSC line | Allen Institute for Cell Science | AICS-0048-039 |
| **Chemicals, Peptides, and Recombinant Proteins** |  |  |
| 2-Mercaptoethanol | Sigma-Aldrich | Cat# M6250 |
| Activin A | Qkine | Cat# Qk001 |
| Ammonium acetate | Invitrogen | Cat# A16343.30 |
| B-27 Supplement | Gibco | Cat# 17504044 |
| BCA Protein Assay Kit | Thermo Fisher | Cat# A55864 |
| BDNF | Qkine | Cat# Qk050-0100 |
| Blasticidin S | Thermo Fisher | Cat# A1113903 |
| BMP4 | Qkine | Cat# Qk038 |
| BSA | Sigma-Aldrich | Cat# A8412-100ML |
| CHIR99021 | Cayman Chemical | Cat# 004CA13122-10 |
| Chroman 1 | MedChemExpress | Cat# HY-15392 |
| CryoStor CS10 | Sigma-Aldrich | Cat# C2874-100ML |
| DAPI | Sigma-Aldrich | Cat# D9542 |
| DMEM High Glucose | Gibco | Cat# 11965092 |
| DMEM/F-12 | Gibco | Cat# 31330038 |
| Doxycycline hyclate | Sigma-Aldrich | Cat# D9891-1G |
| EDTA | Sigma-Aldrich | Cat# 4055-100ML |
| Emricasan | Selleckchem | Cat# S7775 |
| Essential 8 Medium | Gibco | Cat# A1517001 |
| FBS | Life Technologies | Cat# 10270106 |
| FGF2 | Qkine | Cat# Qk025-0100 |
| Fixable viability dye eFluor 450 | Invitrogen | Cat# 65086314 |
| Fixable viability dye eFluor 780 | Invitrogen | Cat# 65086514 |
| Geltrex LDEV-Free | Gibco | Cat# A1413302 |
| Geneticin (G418) | Thermo Fisher | Cat# 10131035 |
| GlutaMAX Supplement | Gibco | Cat# 35050061 |
| Ham's F12 | Gibco | Cat# 31765068 |
| HEPES | Fisher Bioreagents | Cat# 10756254 |
| Hygromycin | Thermo Fisher | Cat# 10687010 |
| IMDM | Gibco | Cat# 21980065 |
| Insulin | Thermo Fisher Scientific | Cat# A11382II |
| ITS-G Supplement | Gibco | Cat# 41400045 |
| LDS Sample Buffer | Invitrogen | Cat# NP0008 |
| Lipofectamine Stem | Invitrogen | Cat# STEM00001 |
| Lipid mixture | Gibco | Cat# 11905031 |
| LY294002 | Selleckchem | Cat# 1105 |
| Magnesium acetate tetrahydrate | Sigma-Aldrich | Cat# M5661 |
| MEM Non-Essential Amino Acids Solution | Gibco | Cat# 11140050 |
| Mini-PROTEAN TGX Gel | Bio-Rad | Cat# 4561096 |
| Monothioglycerol | Sigma-Aldrich | Cat# M6145-100ML |
| mTeSR Plus | STEMCELL Technologies | Cat# 100-0276 |
| Neurobasal Medium | Gibco | Cat# 21103049 |
| Neurotrophin-3 (NT-3) | Qkine | Cat# Qk058-0100 |
| Opti-MEM Reduced Serum Medium | Gibco | Cat# 31985047 |
| P3 Primary Cell Kit | Lonza | Cat# V4XP-3024 |
| Paraformaldehyde (PFA) | Alfa Aesar | Cat# 15434389 |
| PBS | Euroclone | Cat# ECB4053 |
| Polyamine supplement | Sigma-Aldrich | Cat# P8483 |
| Poly-L-lysine | Merck | Cat# P2636 |
| Ponceau S | Sigma-Aldrich | Cat# P3504 |
| ProLong Diamond Mountant | Invitrogen | Cat# P36965 |
| Protease inhibitor cocktail | Roche | Cat# 11873580001 |
| Puromycin | Thermo Fisher | Cat# A1113803 |
| Retinoic acid | Sigma-Aldrich | Cat# R2625-500MG |
| Sodium deoxycholate | Sigma-Aldrich | Cat# D6750 |
| StemFlex Medium | Gibco | Cat# A3349401 |
| StemPro Accutase | Gibco | Cat# A1110501 |
| Tetracycline hydrochloride | Sigma-Aldrich | Cat# T7660-5G |
| Thiazovivin | Cayman Chemical | Cat# 004CA14245-25 |
| Trans-ISRIB | Tocris | Cat# 5284 |
| Transferrin | Invitria | Cat# 777TRF029-1G |
| Triton X-100 | Sigma-Aldrich | Cat# NP40S |
| Trypan Blue | Sigma-Aldrich | Cat# CAS72571 |
| TrypLE Express | Gibco | Cat# 12605010 |
| Trypsin | Euroclone | Cat# ECB3051D |
| WNT-C59 | Cayman Chemical | Cat# 16644 |
| Y-27632 | Cayman Chemical | Cat# 004CA10005583-10 |
| **Critical Commercial Assays** |  |  |
| Chromium 3' Kit v3.1 | 10x Genomics | PN-1000268 |
| Chromium 5' Kit v3 | 10x Genomics | PN-1000699 |
| Dual Index Primers Kit TT Set A | 10x Genomics | Cat# PN-1000215 |
| EndoFree Plasmid Buffer Set | QIAGEN | Cat# 19048 |
| High Capacity cDNA RT Kit | Applied Biosystems | Cat# 4368813 |
| KaryoStat+ Assay | Thermo Fisher | Cat# A52849 |
| NucleoSpin Gel and PCR Clean-up | Macherey-Nagel | Cat# 740609.50 |
| PowerUp SYBR Green Master Mix | Applied Biosystems | Cat# A25743 |
| PureLink Genomic DNA Kit | Thermo Fisher | Cat# K1820-02 |
| QIAEX II Gel Extraction Kit | QIAGEN | Cat# 20021 |
| QIAfilter Plasmid Midi Kit | QIAGEN | Cat# 12243 |
| Qubit dsDNA HS Assay | Invitrogen | Cat# Q332301 |
| Quick-DNA Miniprep Kit | Zymo Research | Cat# D3025 |
| Quick-RNA MiniPrep Kit | Zymo Research | Cat# ZYR1055 |
| Stranded mRNA Prep Ligation Kit | Illumina | Cat# 20040534 |
| TruSeq RNA UD Indexes | Illumina | Cat# 20020591 |
| TruSeq Stranded mRNA Kit | Illumina | Cat# 20020594 |
| **Recombinant DNA** |  |  |
| 20% TBE polyacrylamide gel | Invitrogen | Cat# EC63155BOX |
| AarI restriction enzyme | Thermo Scientific | Cat# ER1581 |
| Agilent TapeStation system | Agilent Technologies | Cat# 5067-5576 |
| BbsI-HF restriction enzyme | NEB | Cat# R3539S |
| DNA High Sensitivity Assay Kit | Invitrogen | Cat# Q33231 |
| Fast Alkaline Phosphatase | Thermo Fisher | Cat# EF0651 |
| High Sensitivity DNA Screen-Tape D5000 | Agilent | Cat# 5067-5588 |
| In-Fusion Snap Assembly | Takara | Cat# 638943 |
| LongAmp Taq DNA Polymerase | NEB | Cat# M0323L |
| NEBNext High-Fidelity 2X PCR Master Mix | NEB | Cat# M0541S |
| NEBuilder HiFi DNA Assembly Mix | NEB | Cat# E5520S |
| PaqCI restriction enzyme | NEB | Cat# R0745S |
| Q5 High-Fidelity Polymerase | NEB | Cat# M0493L |
| SPRISelect Beads | Beckman Coulter | Cat# B24965AA |
| SYBR Gold | Invitrogen | Cat# S11494 |
| T4 DNA Ligase | Thermo Fisher | Cat# EL0011 |
| **Experimental Models: Organisms/Strains** |  |  |
| NEB 5-alpha *E. coli* | NEB | Cat# C2987H |
| TOP10F' *E. coli* | Thermo Fisher | Cat# C303003 |
| **Antibodies** |  |  |
| Mouse anti-ACTN2 | Abcam | Cat# ab9465 |
| Mouse anti-dCas9 | Diagenode | Cat# C15200203-100 |
| Mouse anti-NANOG (Alexa Fluor 647) | BD Biosciences | Cat# 561300 |
| Mouse anti-POU5F1/OCT4 (PE) | BD Biosciences | Cat# 560186 |
| Mouse anti-VCL | Sigma-Aldrich | Cat# SAB4200080 |
| Mouse IgG1 Isotype Control (Alexa Fluor 647) | BD Biosciences | Cat# 566011 |
| Mouse IgG1 Isotype Control (PE) | BD Biosciences | Cat# 555749 |
| Rabbit anti-GAPDH | Cell Signaling Technology | Cat# 2118 |
| Rabbit anti-MYOD1 | Abcam | Cat# ab133627 |
| Rabbit anti-TetR | Mobitec | Cat# TET01 |
| **Software and Algorithms** |  |  |
| Bulk RNA-seq alignment pipeline | GitHub | https://github.com/sara-bianchi/Bulk_RNA_seq_pipeline |
| Cell Ranger | 10x Genomics | v7.2.0 |
| Cell Ranger | 10x Genomics | v9.0.0 |
| crisprVerse pipeline | GitHub | https://github.com/sara-bianchi/CRISPR_guides_design |
| Custom code for CIRI screening | GitHub | https://github.com/alessandro-bertero/CIRI_bioinfo |
| DESeq2 | Bioconductor | v3.23 |
| Docker Desktop | Docker | v4.66.1 |
| FlowJo | BD | v10 |
| GraphPad Prism | GraphPad | v10 |
| ImageJ | NIH | v1.54i |
| Monocle | Trapnell Lab | v3 |
| Seurat | Satija Lab | v5.2.0 |
| STAR aligner | Dobin et al. | v2.7.11b |
| **Deposited NGS data** |  |  |
| Processed data | Zenodo | [DOI:10.5281/zenodo.19556688](https://doi.org/10.5281/zenodo.19556688) |
| Data scRNA-seq Fig 5 | Biostudies | E-MTAB-17009 |
| Data Bulk RNA-seq Fig 5 | Biostudies | E-MTAB-17033 |
| Data scRNA-seq Fig 6 | Biostudies | E-MTAB-17010 |
| Data scRNA-seq Fig S6 | Biostudies | E-MTAB-17012 |
| Data scRNA-seq Fig 7 | Biostudies | E-MTAB-17011 |
| **Plasmids** |  |  |
| pSpCas9n(BB)_R26-L | Addgene | 255612 |
| pSpCas9n(BB)_R26-R | Addgene | 255613 |
| pR26-Bst_CAG-OPTtetR | Addgene | 255614 |
| pZT-C13-L1 | Addgene | 62196 |
| pZT-C13-R1 | Addgene | 62197 |
| pCLYBL-Neo_CAG-dCas9-2A-mCherry | Addgene | 255615 |
| pZFN-AAVS1_ELD | Addgene | 159297 |
| pZFN-AAVS1_KKR | Addgene | 159298 |
| pAAV-Puro_CIRIa-MS2_GFP | Addgene | 255616 |
| pAAV-Puro_CIRIa-MS2_BFP | Addgene | 255617 |
| pAAV-Puro_CIRIa-PP7_GFP | Addgene | 255618 |
| pAAV-Puro_CIRli-com_GFP | Addgene | 255619 |
| pAAV-Hygro_CIRli-com | Addgene | 255620 |
| pAAV-Puro_CIRIa-MS2_BFP_MYOD1x3 | Addgene | 255621 |
| pAAV-Hygro_CIRli-com_OSN | Addgene | 255622 |
| pAAV-Puro_CIRIa-MS2_BFP_NTCx3 | Addgene | 255623 |
| pAAV-Hygro_CIRli-com_NTCx3 | Addgene | 255624 |
| pAAV-Puro_CIRIa-MS2_BFP_E1-MYOD1x2 | Addgene | 255625 |
| pAAV-Puro_CIRIa-MS2_BFP_E2-MYOD1x2 | Addgene | 255626 |
| pAAV-Hygro_CIRli-com_E1-OSN | Addgene | 255627 |
| pAAV-Hygro_CIRli-com_E2-OSN | Addgene | 255628 |
| pAAV-Puro_CIRIa-MS2_BFP_SMARCD3x2-MYOD1x2 | Addgene | 255629 |
| pAAV-Puro_CIRIa-MS2_BFP_KDM6Bx2-MYOD1x2 | Addgene | 255630 |
| pAAV-Hygro_CIRli-com_ID3x2-OSN | Addgene | 255631 |

## RESOURCE AVAILABILITY

### Lead contact

Further information and requests for resources and reagents should be directed and will be fulfilled by the lead contact, Alessandro Bertero.

### Materials availability

Plasmids generated in the study and required to implement the technology have been deposited to Addgene and are listed in the key resources table. The engineered hiPSCs will be made available on request with appropriate MTA.

### Data and code availability

Raw and processed bulk and single-cell RNA-seq data have been deposited at ArrayExpress. Monocle3 CellDataSet class files have been deposited at Zenodo. Accession links and DOIs are listed in the key resources table, and all data will be made publicly available upon publication.

Custom R code used for downstream analyses is available on GitHub at the repository listed in the key resources table. Any additional information required to reanalyze the data reported in this paper is available from the lead contact upon request.

## EXPERIMENTAL MODEL AND STUDY PARTICIPANT DETAILS

### Human cell source

TTN-mEGFP reporter WTC11 hiPSC cell line was cultured in feeder-free conditions at 37 °C, 5% CO2, and normoxia. Briefly, cells were plated on LDEV-free Geltrex-coated culture dishes [Geltrex was diluted in DMEM-F12 at 17 µg/mL], and cultured in commercial medium, either StemFlex, mTeSR Plus, or Essential 8. Cells were passaged in small clumps every 4-5 days using Versene [0.5 mM EDTA diluted in PBS], and replated in medium supplemented with 2 µM Thiazovivin. Media changes were performed 24 h after the split to remove Thiazovivin and then every other day. For cryopreservation, 1 M cells were resuspended in 200 µl CryoStor™CS10 and kept at −150 °C for long-term storage.

## METHOD DETAILS

### Molecular cloning

Unless stated otherwise, cut and paste cloning was performed using restriction enzymes and T4 DNA ligase, while seamless cloning was performed using In-Fusion Snap Assembly Master Mix or NEBuilder HiFi DNA Assembly Master Mix. PCR was performed using Q5 Hot Start High-Fidelity DNA Polymerase using primers from Eurofins Genomics. Synthetic fragments were obtained as gBlock Gene Fragments (IDT). Vectors were dephosphorylated before ligation using Fast Alkaline Phosphatase. QIAEX II Gel Extraction Kit was used for DNA extraction from agarose gels, and Nucleospin Gel and PCR Clean-up for purification of PCR products. Recombinant plasmids were transformed into One Shot TOP10F’ Chemically Competent *E. coli* or NEB™5-alpha Competent *E. coli*. Quick-DNA Miniprep Kit and QIAfilter Plasmid Midi Kit were used for plasmid preparations (using EndoFree Plasmid Buffers for plasmids to be used for genome editing). All these procedures were performed according to the manufacturer’s instructions. Additional molecular biology procedures, such as electrophoresis and *E. Coli* culture, were performed according to standard protocols. All the plasmids were fully sequence-verified by Plasmidsaurus or Eurofins Genomics. Full molecular details for each plasmid, including primer sequences, synthetic fragment sequences, and assembly strategies, are provided in Supplemental Methods.

### sgRNA molecular cloning

Individual protospacers for non-pooled experiments were cloned by digesting the backbone with the AarI type IIS restriction enzyme, generating two non-compatible ends that position the targeting sequence between the Pol III transcription start site and the sgRNA scaffold without leaving restriction scars. Protospacers were introduced by phosphorylation and annealing of complementary oligonucleotides to generate double-stranded inserts with compatible sticky ends. Oligonucleotide sequences are listed in Tables S7–S8. A validated, non-targeting control (NTC) protospacer was used as negative control (Invitrogen TrueGuide), cloned with oligos 5’ - GACCGAAATGTGAGATCAGAGTAAT - 3’ and 5’ - AAACATTACTCTGATCTCACATTTC - 3’.

Vectors carrying multiple sgRNAs were assembled by NEBuilder Hi-Fi DNA assembly of PCR-amplified individual inducible sgRNA cassettes. Assembly primers introduced directional sequence overlaps through three defined block sequences (BL1–BL3; listed in Table S9, which also served as colony PCR primer binding sites to verify correct guide combination and orientation. Block sequences: (BL1) 5’-ACTATGCTGTGTCTTGACAGCAGACCTCGT-3’; (BL2) 5’-TGGACACACAAGTACTGTCGGCAACCACAC-3’; (BL3) 5’-TGAGAACTGCTAGTCTCGTGACAGCGACTT-3’.

The optimization of single- and dual-screening sgRNA cloning strategies are discussed in detail in Supplemental Text, while methods are detailed in Supplemental Methods. The sgRNA sequences are all listed in Table S10.

### Lipofection

For lipofection, hiPSCs were washed with DPBS and dissociated into single cells using Versene for 3 minutes at 37 °C. 200,000 cells were resuspended in medium supplemented with 2 µM Thiazovivin and seeded into a Geltrex-coated well of a 6-well plate. 24 h after seeding, the medium was replaced with Essential 8 supplemented with a transfection mix consisting of 200 µl Opti-MEM Reduced Serum Medium, 2 µl Lipofectamine Stem Transfection Reagent, and a total of 2 µg of plasmid DNA. After 4 h the medium was replaced with fresh culture medium.

### Nucleofection

For nucleofection, hiPSCs were washed with DPBS and dissociated to single cells using Stem-Pro Accutase for 5 minutes at 37 °C. 1 M cells were resuspended in 100 µl P3 Primary Cell 4D-Nucleofector solution and transferred to the conductive polymer nucleovette. For each condition, a total of 10 µg of plasmid DNA was used. The nucleovette was loaded into the Lonza 4D-Nucleofector System X Unit and subjected to the CA137 program. Nucleofected cells were seeded in a well of a 24-well plate pre-coated with Geltrex, using recovery medium supplemented with CEPT cocktail^87^ [50 nM Chroman1, 5 µM Emricasan, Polyamine supplement 1:1000, 0.7 µM Trans-ISRIB]. The following day, the medium was refreshed to remove CEPT. 48 h post-nucleofection, cells were passaged in a 10 cm culture dish to proceed with antibiotic selection.

### Genome editing

*hROSA26* genome editing was performed using CRISPR/Cas9n, as previously described^23^. Briefly, hiPSCs were co-transfected with pSpCas9n(BB)_R26-L, pSpCas9n(BB)_R26-R, and pR26-Bst_CAG-OPTtetR (1:1:2 ratio) to promote HDR-mediated integration of the OPTtetR transgene. 48 h post-transfection, cells were selected with 8 µg/mL Blasticidin S for 4 days.

*CLYBL* genome editing was performed using transcription activator-like effector nucleases (TALENs), as previously described^33^. Briefly, hiPSCs were co-nucleofected with pZT-C13-L1, pZT-C13-R1, and pCLYBL-Neo_CAG-dCas9-2A-mCherry (1:1:2 ratio) to promote HDR-mediated integration of the dCas9 transgene. 48 h post-transfection, cells were selected with Geneticin starting at 50 µg/mL for 48 h, then reducing the concentration to 25 µg/mL on day 3 and to 12.5 µg/mL on the final day of selection.

*AAVS1* genome editing was performed using obligate heterodimer zinc finger nucleases (ZFNs), as previously described^23^. Briefly, hiPSCs were co-nucleofected with pZFN-AAVS1_ELD, pZFN-AAVS1_KKR, and one or two pAAV targeting plasmids (1:1:2 or 1:1:1:1 ratios) to promote HDR-mediated integration of the effectors transgene. 48 h post-transfection, cells were selected for 5 days with 0.5 µg/mL puromycin, hygromycin B, or both, depending on the donor cassette(s). Hygromycin B selection was performed at 25 µg/mL for 1 day, followed by 12.5 µg/mL for the remainder of selection.

### Genotyping

hiPSC pseudoclones from genome editing experiments were manually picked, expanded, and screened by genomic PCR to verify site-specific on-target integration of the transgene and to detect random integrations of the targeting plasmid elsewhere in the genome. The genotyping strategy is described in Figure S1A.

Site-specific transgene integration (INT) was determined by 5’INT and 3’INT junctional PCRs relying on one primer on the transgene 5’ or 3’ end, respectively, and a second primer on the genomic locus outside of the relevant homology arm. Wild-type (WT) PCR was performed using the two primers mapping to the genomic locus outside of the homology arms to determine if only one or both of the two alleles were edited (PCR failed for alleles containing the GC-rich CAG promoter used in all our targeting plasmids).

To monitor possible off-target random integration, two additional backbone (BB) PCRs were performed: 5’BB and 3’BB PCRs, relying on a primer specific to the 5’ or 3’ of the transgene and a second primer on the plasmid backbone outside of the relevant homology arms. Upon successful HDR only the homology arms are incorporated in the GSH, while the rest of the plasmid backbone is lost; therefore, BB PCRs reveal other events of random integration of the linearized plasmid. PCRs were performed with LongAmp Taq DNA Polymerase, according to manufacturer’s instructions. All the primer combinations are listed in Table S11.

### Molecular karyotyping

A frozen pellet of 2 million TetCas hiPSCs was analyzed for potential copy number variations (CNVs) with the KaryoStat+ Genetic Stability Assay Service. Genomic DNA was purified using the PureLink Genomic DNA Purification Kit and 100 ng was used to prepare the GeneArray for KaryoStat, all according to the manufacturer’s instructions. Service reports identified the known chrY p11.2 deletion present in the WTC11 donor background and a small apparent chromosome 7 haploid-state signal that was later confirmed to be present in the parental WTC11 *TTN*-mEGFP hiPSCs used in this study.

For CIRI and OPTi-OX clones CNV analysis, genomic DNA was analyzed by low-resolution SNP-array karyotyping using an Illumina 700K+ marker array, performed by Life & Brain GmbH (Bonn, Germany). SNP-array data were processed in GenomeStudio 2.0.5 with the cnvPartition 3.2.1 plugin using default parameters, except for a minimum probe count of 6 and confidence threshold of 40. Bookmark Analysis XML files were parsed and filtered with custom R scripts. CNVs were retained if they had confidence >85 and were either >350 kbp, or >100 kbp when overlapping recurrent hPSC CNV hotspots.

### Induction of sgRNA expression

Tetracycline was added to the culture medium at 1 µg/mL to induce sgRNA expression. The treatment duration is described for each experiment in the results and/or figure legends.

### Cardiac organoid differentiation

Left ventricle cardiac organoids were generated as previously described^38^, with minor modifications^52^. All media were prepared starting from a basal chemically defined medium, CDM containing 50% IMDM and 50% Ham’s F12 supplemented with 0.4% BSA, 1% lipid mixture, 0.004% monothioglycerol, and 15 µg/mL transferrin.

50,000 hiPSCs/well were seeded in 24-well plates in the presence of 5 µM Y-27632. After 24 h, the medium was replaced with mesoderm induction medium: CDM supplemented with 6 ng/mL FGF2, 5 µM LY294002, 5 ng/mL Activin A, 8 ng/mL BMP4, and 5 µM CHIR99021. After 36 h, cells were dissociated using TrypLE E, resuspended in cardiac mesoderm medium: CDM supplemented with 8 ng/mL BMP4, 1.6 ng/mL FGF2, 10 µg/mL insulin, 2 µM WNT-C59, and 50 nM retinoic acid, 5 µM Y-27632; and 15,000 cells/well were transferred to ultra-low-attachment 96-well. Plates were centrifuged at 140 g for 4 min to promote aggregation. From day 2.5 onward, spheroids were maintained in fresh cardiac mesoderm medium without Y-27632 until day 5.5, when the medium was replaced with cardiac differentiation medium: CDM medium supplemented with 8 ng/mL BMP4, 1.6 ng/mL FGF2, and 10 µg/mL insulin.

Before tetracycline treatment, organoids were imaged using an Incucyte live-cell imaging system to verify dCas9-2A-mCherry expression and TTN-mEGFP reporter status. Organoids were then treated with 1 µg/mL tetracycline on day 6.5 and analyzed at day 7.5.

### Skeletal muscle forward programming

Myogenic CIRI hiPSCs were washed with DPBS and dissociated into single cells using StemPro Accutase for 3 minutes at 37 °C. 100,000 cells were resuspended in StemFlex supplemented with 2 µM Thiazovivin and plated onto a Geltrex-coated well of 24-well dish. The following day, the medium was replaced with StemFlex medium supplemented with 1 µg/mL tetracycline to induce the expression of the sgRNAs. From the following day, cells were cultured in myogenic medium [DMEM High Glucose supplemented with ITS-G 1:100, 40 ng/mL FGF2 145 aa, 1 µM retinoic acid, 3 µM CHIR99021, and 1 µg/mL tetracycline], performing daily medium changes. Where indicated in the results, -RA denotes the same myogenic medium lacking retinoic acid. Cells were analyzed after 7 days of differentiation, unless stated otherwise.

The *MYOD1* OPTi-OX line was used as a benchmark control. This line was generated from a previously described hiPSC line carrying a heterozygous hROSA26-targeted rTTA cassette^24^. An *AAVS1* donor construct encoding doxycycline-inducible *MYOD1* cDNA, based on the OPTi-OX design described by Pawlowski et al.^42^, was synthesized and integrated into the *AAVS1* safe harbor according to the original strategy. A homozygous *AAVS1*-targeted clone was isolated and used as a benchmark control in this study. For skeletal muscle forward programming, this line was differentiated using the same protocol applied to CIRI clones, except that *MYOD1* cDNA expression was induced with 1 µg/mL doxycycline instead of tetracycline.

### Neuronal forward programming

Neurogenic CIRI hiPSCs were washed with DPBS and dissociated into single cells using Versene for 3 min at 37 °C. Cells were resuspended in Essential 8 medium supplemented with 2 µM Thiazovivin and plated onto Geltrex-coated 12-well plates at a density of 75,000 cells per well. After 24 h, the medium was replaced with induction medium [DMEM/F-12 supplemented with 1X GlutaMAX, 1X MEM Non-Essential Amino Acids, 50 µM 2-mercaptoethanol, and 1 µg/mL tetracycline]. After 2 days of induction, the medium was switched to Neurobasal medium supplemented with 1X GlutaMAX, 1X B-27, 10 ng/mL BDNF, 10 ng/mL Neurotrophin-3 (NT-3), and 1 µg/mL tetracycline. Medium changes were performed daily, and cells were cultured for 14 days before analysis.

### Reverse transcription quantitative PCR

Total RNA samples for RT-qPCR were extracted with Quick-RNA MiniPrep Kit. cDNA synthesis was performed with random primers and 500 ng of total RNA using the High Capacity cDNA Reverse Transcription Kit. qPCR reactions were prepared in 10 µL with 10 ng cDNA, 400 nM of each PCR primer and PowerUp SYBR Green 2X Master Mix. qPCR reactions were carried out in technical duplicates on QuantStudio 6 Flex Real-Time PCR System machine from Applied Biosystems. When possible, primers were designed to span exon–exon junctions and/or to amplify across introns, enabling cDNA to be distinguished from genomic DNA. Control samples without reverse transcriptase were analyzed to confirm that genomic DNA contamination was not impactful. Undetectable Ct values were assigned to Ct = 40, corresponding to the maximum number of amplification cycles, before ΔCt calculation. Expression levels were calculated relative to the indicated housekeeping gene using the 2^-ΔCt^ method, with ΔCt = Ct_target_ – Ct_housekeeping_, and are displayed as percentage of housekeeping gene expression. *HPRT1* was used as housekeeping gene for hiPSCs and *RPLP0* for differentiated derivatives. Because relative-expression values derived from 2^-ΔCt^ can span several orders of magnitude and show unequal variance, statistical analyses of RT-qPCR data were performed on the corresponding ΔCt values, which better approximate the assumptions of parametric tests^88^. A list of all the RT-qPCR primers is reported in Table S12.

### Flow cytometry

Expression of fluorescent markers was measured in live cells. Cells were harvested using a solution of TrypLE 1X and Versene (at 3:1 ratio) and incubated at 37 °C for 3 min. The cell suspension was filtered through a 40 µm cell strainer. After centrifugation at 100 g for 5 min, cells were resuspended in sorting buffer [PBS with 1% P/S, 2% FBS, and 10 mM HEPES]. Fixable Viability Dye eFluor 780 was added to the sorting buffer to label dead cells. Live cells were kept on ice, and analyzed using BD FACSVerse Cell Analyzer or Sony SH800S. Data analysis was then performed with FlowJo (v10).

Analyses of pluripotency markers was performed on fixed cells. After harvesting as described above, cells were stained with Fixable Viability Dye eFluor 450 and washed with FACS Buffer [PBS with 5% FBS and 0.05% sodium azide]. Cells were then fixed with 4% PFA for 10 minutes at room temperature (RT), washed with FACS Buffer, and permeabilized with 0.1% Triton X-100 (diluted in PBS) for 20 minutes. Cells were stained for 1 h at RT with fluorophore-conjugated antibodies diluted in staining solution (PBS with 0.75% Saponin, 5% FBS). Antibodies and their working concentrations are reported in Table S13. After 2 washes with staining solution, cells were analyzed with BD FACSVerse Cell Analyzer. All samples were also separately stained with isotype negative control antibodies. Data was analysed with FlowJo (v10).

### Fluorescence-activated cell sorting

For clonal isolation of live hiPSCs, cells were prepared as described for flow cytometry and sorted onto Geltrex-coated 96-well culture dishes containing medium supplemented with CEPT cocktail. The Sony SH800S cell sorter was calibrated with a 100 µm microfluidics chip in targeted mode and operated in the “single cells 3 drops” setting at minimal pressure, not exceeding approximately 100 events/s. After 72 h, the medium was replaced to remove CEPT, and surviving colonies were expanded until ready for manual passaging.

For sorting of cardiac organoid-derived cells, organoids from each condition were pooled after supernatant removal, washed twice with Versene, and incubated in 0.5% trypsin diluted in Versene for 10–20 min at 37 °C, with gentle mixing every 5 min to promote dissociation into single cells. Enzymatic dissociation was stopped by adding ice-cold CDM supplemented with 10% FBS. Cells were centrifuged at 200 g for 5 min at 4 °C, resuspended in FACS buffer [PBS supplemented with 2% FBS and 10 mM HEPES] containing Fixable Viability Dye eFluor 780 to exclude dead cells, filtered, and kept on ice before analysis and sorting. Flow cytometry analysis and cell sorting were performed using a Sony SH800S cell sorter calibrated with a 100 µm microfluidics sorting chip. Live single cells were analyzed for dCas9-2A-mCherry and TTN-mEGFP expression, and TTN-mEGFP^+^ and TTN-mEGFP^-^ populations were collected. Sorted cells were centrifuged and resuspended in Quick-RNA MiniPrep lysis buffer for RNA extraction.

### Western blot

Cells were washed with PBS and detached using Versene (0.5 mM EDTA in PBS) for 2 minutes at 37 °C. Pelleted cells were lysed in RIPA buffer [50 mM HEPES pH 7.5; 500 mM LiCl; 1 mM EDTA pH 8; 1% NP-40; 1% sodium deoxycholate] supplemented with 1X protease inhibitor cocktail. Lysates were sonicated (3 × 30 s) and centrifuged at 10,000 × g for 10 min at 4 °C. Protein concentration was determined using the BCA assay kit, following the manufacturer’s instructions. Proteins were denatured in LDS sample buffer supplemented with 2-mercaptoethanol 4%, resolved on 4–12% Mini-PROTEAN TGX gel, and transferred to nitrocellulose membranes using a wet transfer system (Bio-Rad). Membranes were stained with Ponceau S, blocked in 5% non-fat dry milk in TBS-T, and incubated overnight at 4 °C with primary antibodies. All the antibodies used and their working concentrations are shown in Table S13. After washing, membranes were incubated with HRP-conjugated secondary antibodies in TBS-T (anti-mouse IgG 1:5,000 or anti-rabbit IgG 1:3,000) and visualized using chemiluminescence on a ChemiDoc imaging system (Bio-Rad).

### Immunofluorescence

hiPSCs and differentiated derivatives were cultured on glass slides pre-treated with 0.01% poly-L-lysine for 1 h at RT and then coated with Geltrex. Cells were fixed in 4% paraformaldehyde for 10 minutes at RT and washed three times with PBS. The cells were then permeabilized with 0.1% Triton X-100 (diluted in PBS) for 20 minutes at RT and blocked with 3% BSA and 1% goat serum in PBS for 1 h at RT. Subsequently, cells were incubated for 1 h at RT with the primary antibody diluted in blocking buffer. All the antibodies used and their working concentrations are shown in Table S13. After five washes with PBS, the cells were incubated for 1 h at RT protected from light with the corresponding fluorophore-conjugated secondary antibody (Alexa Fluor 488 or 647) in PBS + 1% BSA. Nuclei were stained with 100 ng/mL DAPI. The glass slides were mounted with ProLong Diamond Mounting medium. Leica TCS SP8 confocal microscope was used for image acquisition. Images were analyzed with ImageJ (v1.54i).

### sgRNA design

CRISPR protospacers used for the screening were designed using a modified version of CRISPR-Verse, run inside a Docker environment available on a GitHub repository. Different target windows around the TSS were selected for CRISPRa (−500 bp to +50 bp) and CRISPRi (−50 bp to +500 bp). Protospacers were ranked based on an alignment score accounting for potential off-target binding sites and a chromatin accessibility score based on ATAC-seq data. Four guides per target were then selected based on their position relative to the TSS, to span the desired regulatory region. The nuclease was set to spCas9 and the reference genome was UCSC hg38. ATAC-seq data were downloaded from GEO accession GSE106689; replicate 2 of ES cells (GSM2845469) was selected because it had the highest sequencing depth. All selected guides are listed in Table S10. As NTC guides, we used validated sequences from the literature: 5’–GAAATGTGAGATCAGAGTAA–3’ (Invitrogen TrueGuide sgRNA Negative Control, non-targeting 1) and 5’–GTATTACTGATATTGGTGGG–3’ (non-targeting control gRNA BRDN0001149198 from Addgene plasmid #80248).

### Bulk RNA sequencing

Total RNA was extracted using the Quick-RNA MiniPrep Kit, following the manufacturer’s instructions, from undifferentiated hiPSCs and from day 7 forward-programmed myogenic CIRI cultures. Differentiation time-matched *MYOD1* OPTi-OX and NTC cultures were used as controls. RNA concentration and integrity were assessed using the RNA Agilent screen tape and TapeStation system. Only samples with high RNA integrity (RIN > 8) were used for library preparation. mRNA libraries were prepared using the Illumina TruSeq® Stranded mRNA Library Prep Kit, in combination with the IDT for Illumina – TruSeq RNA UD Indexes and Illumina Stranded mRNA Prep Ligation Kit, according to the manufacturer’s guidelines. Libraries were quantified using the Qubit dsDNA High Sensitivity Assay Kit and quality was verified using Agilent TapeStation DNA ScreenTape.

Sequencing was performed on an MGI G400 platform using a T7 flow cell PE100 V3 G400, generating paired-end reads of 100 bp. Raw reads were trimmed in paired-end mode to remove adapter sequences (5’-CTGTCTCTTATACACATCT - 3’). An additional trimming step removed 5 bases from the 5’ and 1 base at 3’ end to eliminate technical artifacts. Reads were aligned to a custom GRCh38.112.AB004 reference genome generated from ENSEMBL GRCh38.112, to which the exogenous sequences of OPTtetR, *NANOG*, *MYOD1*, dCas9, MCP-PH-BFP, and COM-K were appended to both the reference GTF and FASTA files using the cat command. Alignment was performed with STAR (v2.7.11b) in a Docker environment using the pipeline available on a GitHub repository.

Raw STAR gene count files (ReadsPerGene.out.tab) were imported into R and merged into a gene-by-sample count matrix using unstranded read counts. Sample metadata were matched to the count matrix by sample identifier. Genes were annotated using Ensembl via biomaRt, and downstream analyses were restricted to protein-coding genes and custom features. A DESeq2 dataset was generated from raw counts and filtered to retain genes with at least 3 counts in a minimum number of samples corresponding to the smallest replicate group across conditions. Variance-stabilized counts obtained using the DESeq2 vst function were used for quality assessment, including library complexity, gene detection, MA plots, Pearson correlation, and principal component analysis (PCA). Differential expression analysis was performed in DESeq2 using a design formula of ∼ group for each comparison after subsetting the relevant samples and setting the appropriate reference group. Genes with an adjusted *p*-value < 0.05 and |log_2_ fold change| > 1 were considered differentially expressed. Functional enrichment analyses were performed using Gene Ontology (GO), KEGG, and ranked GSEA.

### Single cell RNA sequencing

Single-cell suspensions were prepared from skeletal muscle samples collected on day 7 of differentiation by enzymatic dissociation using Trypsin 0.25% diluted in Versene at 37 °C for 5 minutes. To neutralize trypsin, stop solution was added (myogenic media supplemented with 20% FBS). Cells were subsequently washed with PBS and resuspended in PBS for downstream applications. Cell viability and concentration were assessed manually using a hemocytometer. To validate the heterogeneity of cells generated by myogenic forward programming, single-cell libraries were prepared from the parental CIRI and CRISPRa lines using the Chromium Next GEM Single Cell 3*^′^* LT Kit v3.1 (10x Genomics), following the manufacturer’s instructions (User Guide CG000399 Rev B), with approximately 3,300 cells loaded per reaction.

To validate genetically modified CIRI clones, single-cell libraries were prepared using the GEM-X Universal 5*^′^* Gene Expression v3 kit with OCM multiplexing (10x Genomics), according to the manufacturer’s instructions (User Guide CG000770 Rev B), with approximately 8,320 cells loaded per sample/reaction.

Next-generation sequencing of the 3*^′^* LT scRNA-seq libraries was performed on an MGI G400 platform using a T7 PE100 V3 flow cell (PE100 V3 G400), generating 100 bp paired-end reads. Sequencing and read demultiplexing were performed as described above for bulk RNA-seq. The 5*^′^* OCM libraries were sequenced on an Illumina NextSeq 1000 platform using a P2 XLEAP 100-cycle kit with the following read configuration: Read 1, 28 cycles; Read 2, 90 cycles; and i7/i5 indexes, 10 cycles each. Libraries were loaded at 650 pM with 1% PhiX.

Reads from the 3*^′^* LT dataset were aligned using the cellranger multi pipeline in Cell Ranger v7.2 for each of the two samples (CIRI and CRISPRa), using a custom reference genome analogous to that used for bulk RNA-seq and generated with cellranger mkref in Cell Ranger v7.2. The reference is publicly available on Zenodo. Sample-level count matrices were subsequently combined using cellranger aggr to generate a single-cell .h5 matrix for downstream analysis. These steps were performed within a Docker environment based on the image hedgelab/rstudio-hedgelab:iPS2seq_new_cellranger7. Reads from the 5*^′^* OCM dataset were processed using Cell Ranger v9 with the appropriate pipeline for 5*^′^*gene expression and OCM multiplexing, using GRCh38-2024A reference downloaded from the 10x Genomics website. These analyses were performed within a Docker environment based on the image hedgelab/rstudio-hedgelab:iPS2seq_new_cellranger9.

Further downstream analysis was performed using Seurat and Monocle3. A Seurat object was generated from the filtered feature-barcode matrix after restricting the count matrix to protein-coding genes. Cell-level metadata were reconstructed from Cell Ranger output and merged with the aggregation information. Quality-control metrics were calculated for each cell, including the number of detected genes (nFeature_RNA), total UMI counts (nCount_RNA), per-centage of mitochondrial transcripts, and percentage of ribosomal transcripts. Cells from the 3*^′^* LT dataset with fewer than 300 detected genes, fewer than 200 UMIs, mitochondrial transcript percentage ≥ 15%, or ribosomal transcript percentage ≥ 18% were excluded. Cells from the 5*^′^* OCM dataset with fewer than 300 detected genes, fewer than 100 UMIs, mitochondrial transcript percentage ≥ 10%, or ribosomal transcript percentage ≥ 1% were filtered. For both datasets, data were scaled and subjected to PCA, and the first 30 principal components were used for graph-based clustering and UMAP visualization in Seurat.

Filtered counts and metadata were converted into a cell_data_set object in Monocle3. Monocle3 preprocessing was performed with 30 dimensions, followed by UMAP dimensionality reduction and graph-based clustering across multiple resolutions. The clustering solution selected for downstream analysis for the 3*^′^* LT dataset corresponded to a resolution of 8 × 10^-4^, and a principal graph was learned using learn_graph. For the 5*^′^* OCM dataset the clustering solution selected for downstream analysis was obtained with k = 15 and a resolution of 2×10^-4^, followed by principal graph learning with learn_graph.

For both datasets, cluster marker genes were identified using Monocle3 top_markers. Additionally, differential expression analyses between clusters were performed in Seurat using the Wilcoxon rank-sum test implemented in FindMarkers, with logfc.threshold = 0 and no minimum expression cutoff. Based on marker expression, clusters were annotated.

The sarcomeric core score was calculated to quantify the enrichment of a curated set of sarcomere-associated structural genes, including key components of thick and thin filaments, Z-discs, M-lines, and regulatory/troponin complexes (*TTN*, *NEB*, *ACTN2*, *TCAP*, *MYBPC1*, *MYBPC2*, *MYL1*, *MYL2*, *TNNI1*, *TNNI2*, *TNNT1*, *TNNT3*, *TPM1*, *TPM2*, *MYOM1*, *MYOM2*, *CSRP3*). The score was computed using Seurat’s AddModuleScore function, which summarizes the relative expression of these genes per cell after comparison with expression-matched control genes.

### Single cell RNA sequencing CRISPR screening

scRNA-seq for the CRISPR screening experiments was performed using the Chromium GEM-X Single Cell 5’ Reagent Kit v3 (10x Genomics), which enables simultaneous capture of mRNA transcripts and CRISPR guide RNAs through a 5’ scRNA-seq workflow. hiPSCs targeted with pooled sgRNA libraries were differentiated into skeletal muscle following the protocol described above, with or without retinoic acid supplementation. Single-cell suspensions were prepared as described in the previous section. Cell viability was evaluated by Trypan Blue staining, and only samples with viability >85% were used for library preparation.

Approximately 29,000 cells were loaded onto the Chromium X controller for GEM generation and barcoding. To enable sgRNA detection the Single Cell 5’ FeatureBarcode CRISPR Enrichment Kit was used according to the manufacturer’s instructions (10x Genomics User Guide CG000734 Rev A). Libraries were assessed using High Sensitivity DNA ScreenTape D5000 and quantified with Qubit dsDNA High Sensitivity Assay Kit. Sequencing was performed on an Illumina NextSeq 2000 platform using a P4 XLEAP-100 cycle kit with the following configuration: 28 cycles for Read 1, 91 cycles for Read 2, and 10 cycles for the index read.

Raw base call files were demultiplexed using cellranger mkfastq in Cell Ranger v7.2. The resulting FASTQ files were subsequently processed using Cell Ranger v9 for alignment, gene-expression quantification, and CRISPR guide-feature counting, following the 10x Ge-nomics workflow for 5*^′^* gene expression with CRISPR guide capture. Reads were aligned to the GRCh38-2024A reference genome downloaded from the 10x Genomics website. For each sample, cellranger multi was run using a feature reference CSV file containing the CRISPR guide sequences, enabling simultaneous processing of gene-expression and guide-capture li-braries. After sample-level processing, count matrices were combined using cellranger aggr to generate an aggregated single-cell .h5 matrix for downstream analysis. All Cell Ranger steps were performed within Docker environments based on the images described in the previous section.

Guide assignment and quality control were performed using an adapted version of our catcheR workflow^54^ within the Docker environment provided in the associated GitHub repository. Cells were first classified based on the detection of fixed CRISPRa and CRISPRi guide sequences, which define the perturbation modality detected in each cell. Fixed guides consisted of two *MYOD1* sgRNAs for CRISPRa and sgRNAs targeting *POU5F1/OCT4*, *SOX2*, and *NANOG* for CRISPRi. For each fixed guide modality, UMI-count distributions were modelled using Kernel Density Estimation (KDE). The minimum UMI threshold for guide detection was defined as the first local minimum separating the low-UMI background peak from the higher-UMI signal peak. To ensure sufficient resolution across samples with different sequencing depths, KDE curves were estimated using a dynamically defined number of points, corresponding to the next power of two covering at least twice the maximum observed UMI count, with lower and upper bounds of 1,024 and 16,384 points, respectively. A bandwidth adjustment of 0.1 was used, and local minima were identified from sign changes in the first derivative of the KDE curve. In the single-guide experiment, fixed-guide detection thresholds were 23.5 UMIs for CRISPRa and 178.8 UMIs for CRISPRi. In the dual-guide experiment, the fixed CRISPRi threshold was 292.2 UMIs. Since the fixed CRISPRa distribution in the dual-guide experiment did not show a clear bimodal separation, the CRISPRa threshold was set to 23.5 UMIs, matching the single-guide experiment.

Variable guide assignment was then performed separately for CRISPRa and CRISPRi perturbations, only in cells passing the corresponding fixed-guide threshold. In the single-guide experiment, a variable guide was assigned when the top-ranked guide had at least 10 UMIs and was supported by at least five times more UMIs than the second-ranked guide. In the dual-guide experiment, the two top-ranked variable guides were assigned when their combined UMI count was at least 4 and at least ten-fold higher than the UMI count of the third-ranked guide. In case of ties between the second- and third-ranked guides, priority was given to the guide targeting the same gene as the top-ranked guide. Because dual-guide constructs were designed to encode two variable guides targeting the same gene, cells assigned to discordant guide pairs were removed. Cells not satisfying these assignment criteria were considered ambiguous and excluded from downstream perturbation-level analyses.

Single-cell transcriptome quality control was then performed on the assigned cells. For each cell, mitochondrial and ribosomal transcript percentages were calculated and inspected. Cells with fewer than 250 detected genes were removed. Protein-coding genes were retained, and genes with fewer than 3 total UMIs across the dataset were filtered out. The filtered matrices were processed with Monocle3 for dimensionality reduction and clustering. Clustering was performed using a resolution of 1 × 10^-5^ for the dual-guide experiment and 0.5 × 10^-4^ for the single-guide experiment. For each perturbation combination, the fraction of assigned cells mapping to this cluster was calculated, and for visualization and follow-up experiments, cells with such fraction fewer than 10 were excluded.

### Statistical analysis

Statistical analyses were performed using GraphPad Prism 10. The type and number of replicates, the statistical test used, and the test results are described in the figure legends. Unless stated otherwise, all graphical data are presented as mean ± standard error of the mean. When representative results are presented, the experiments were reproduced in two or more independent cultures. Bulk RNA-seq differential expression was assessed in DESeq2 with Benjamini–Hochberg correction. Single-cell differential expression was performed in Seurat using Wilcoxon rank-sum tests. Fisher’s exact tests were used where indicated for cluster-composition comparisons.

