## Supplemental Information for "Combinatorial and Inducible CRISPRa/i Enables Canalized hiPSC Forward Programming and Iterative Refinement *via* Single-Cell Genomics"

#### Supplemental Items

Supplemental Text

Figure ED1. Optimization of pooled sgRNA cloning strategy to overcome U6 promoter-mediated recombination

Figure ED2. Pooled single- and dual-guide sgRNA cloning strategies

Supplemental Methods

Figure S1. Extended optimization of sgRNA scaffolds for orthogonal CRISPRa/i in hiPSCs

Figure S2. Extended generation and validation of TetCas hiPSCs

Figure S3. Extended validation of inducible CRISPRa/i in hiPSCs and cardiac organoids

Figure S4. Extended validation of combinatorial CRISPRa/i in hiPSCs

Figure S5. Extended characterization of CIRI-mediated myogenic forward programming

Figure S6. Extended characterization of pooled combinatorial CRISPRa/i screening

Figure S7. Extended validation of enhanced retinoic acid-independent myogenic programming

Table S1. Genotyping and flow-cytometry results for sgRNA recruitment architecture testing

Table S2. Genotyping and flow-cytometry results for dCas9-2A-mCherry *CLYBL* targeting

Table S3. Genotyping and flow-cytometry results for inducible CRISPRa and CRISPRi targeting

Table S4. Genotyping results for CIRI and CRISPRa clones

Table S5. Guide-capture QC for single- and dual-sgRNA screens

Table S6. Genotyping and flow-cytometry results for screen-hit validation

Table S7. Cloning primers for CRISPRa sgRNAs

Table S8. Cloning primers for CRISPRi sgRNAs

Table S9. Cloning primers for multi-guide sgRNA cassettes

Table S10. Designed CRISPRa/i sgRNAs for screening targets

Table S11. Genotyping primers

Table S12. RT-qPCR primers

Table S13. Antibodies

### Supplemental Text

#### Optimization of pooled molecular cloning for CIRI genetic screens

The sgRNA molecular cloning workflow required extensive iterative optimization. Indeed, both Gibson assembly and ligation-based cloning strategies using our original plasmids with multiple U6-TO-driven sgRNAs proved highly prone to self-assembly without the sgRNA insert. Sanger sequencing indicated that these recombination events occurred between homologous regions of two consecutive U6-inducible promoters, generating plasmids lacking the first sgRNA sequence (Figure ED1A).

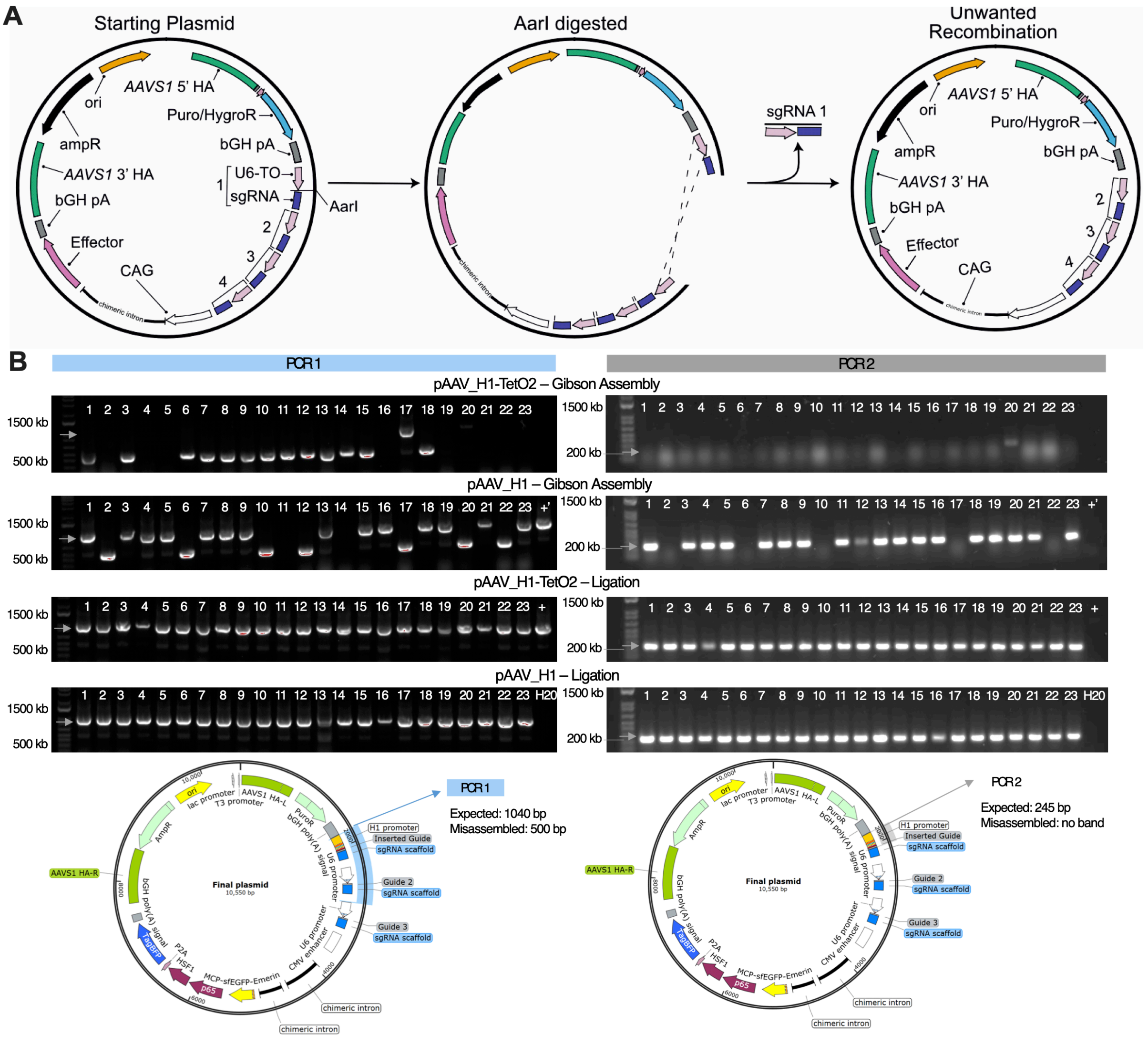


#### Extended Data Figure ED1. Optimization of pooled sgRNA cloning strategy to overcome U6 promoter-mediated recombination

(A) Schematic of the unwanted recombination event frequently observed between identical U6-TetO2 promoters.
(B) Development of a pooled sgRNA cloning strategy that overcomes the limitation shown in panel A. Plasmids carrying alternative promoters were tested using Gibson assembly or ligation. Individual bacterial clones were screened by colony PCR using two primer pairs: PCR 1 to exclude unwanted recombination events, and PCR 2 to confirm specific insert presence. Substitution of U6-TetO2 with H1-TetO2 or H1 promoters, combined with a ligation-based protocol, enabled 100% pooled cloning efficiency.

To address the issue, we replaced the first U6 promoter with a similar Tet-inducible H1 promoter (H1-TO) harboring the same TetO2 sequence, a 20 bp motif recognized and bound by the Tet repressor. Wary that this repeated sequence may still be sufficient to drive plasmid self-assembly, we also generated plasmids with a truncated H1 promoter lacking the TetO2, which would instead be reintroduced during cloning together with the sgRNAs. We then compared Gibson assembly and ligation to determine the most efficient workflow for pooled sgRNA cloning. Gibson assembly was ineffective when using the H1-TO plasmid as backbone, and yielded similarly poor results with the H1 plasmid. In contrast, the ligation-based cloning strategy achieved 100% efficiency with both backbones, and was therefore selected for subsequent experiments (Figure ED1B). Having identified an effective cloning strategy, we developed two distinct protocols for single-guide and dual-guide screens.

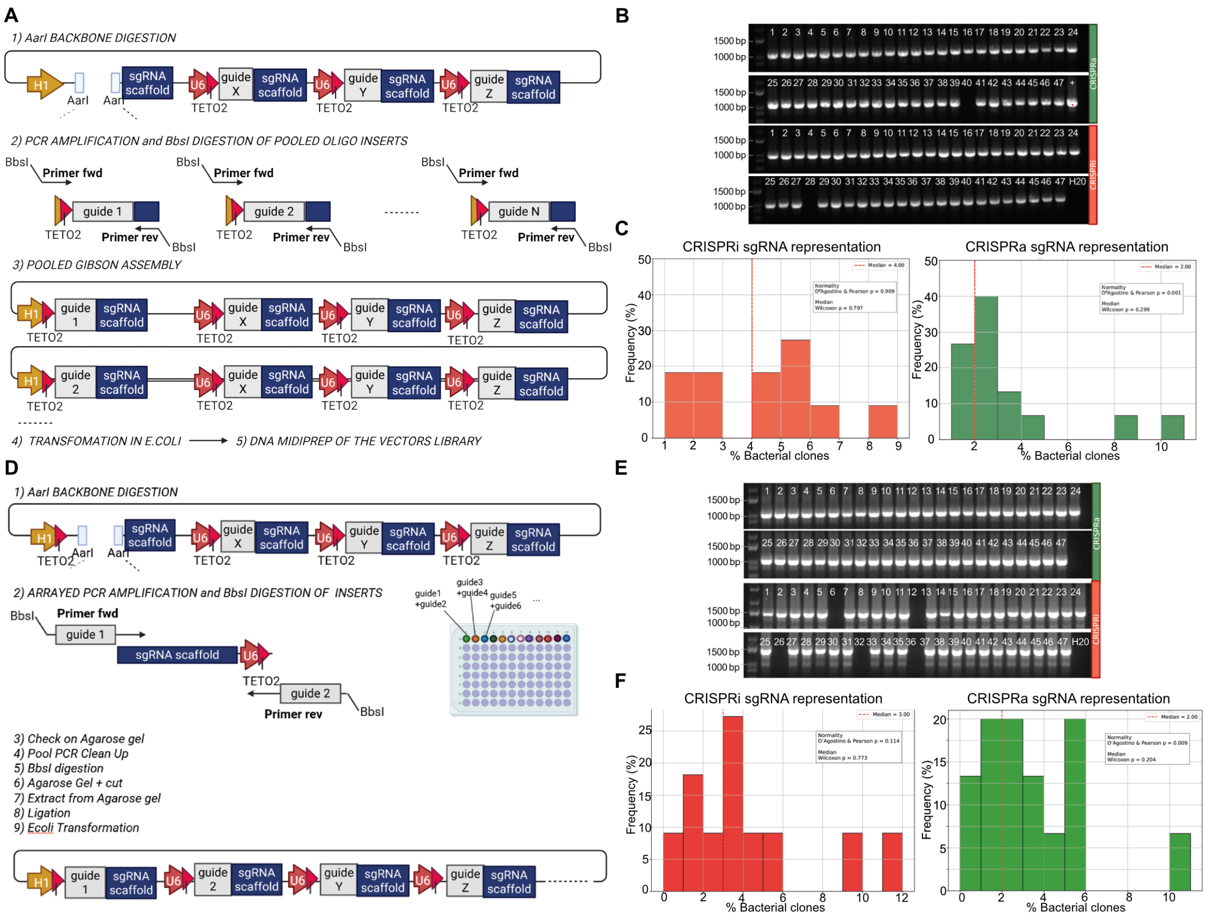


#### Extended Data Figure ED2. Pooled single- and dual-guide sgRNA cloning strategies

(A) Schematic of the pooled single-guide cloning workflow.

(B) Colony PCR on individual bacterial clones after pooled single-sgRNA cloning in CRISPRi and CRISPRa backbones. Expected band: 1044 bp. “+” = original plasmid control.

(C) Quantification of sgRNA representation measured by Sanger sequencing of bacterial colony PCR amplicons from panel B. Dashed lines indicate the expected median representation. Normality was tested by D’Agostino–Pearson test, and deviation from the expected median was tested by Wilcoxon signed-rank test. N = 47 clones.

(D) Schematic of the pooled dual-guide cloning workflow.

(E) Colony PCR on individual bacterial clones after pooled dual-sgRNA cloning in CRISPRa (top two rows) and CRISPRi backbones (bottom two rows). Expected bands: 1050 bp for CRISPRa and 1470 bp for CRISPRi.

(F) Quantification of sgRNA representation measured by Sanger sequencing of bacterial colony PCR amplicons from panel E. Plotting and statistical analysis as described for panel C. N = 47 clones.

We first generated pooled single-sgRNA libraries by inserting individual guides into plasmids already containing two or three sgRNAs. In this workflow, guides together with the TetO2 sequence were PCR-amplified in pools, digested with a type IIS restriction enzyme to generate compatible ends, and subsequently cloned into the final vector to obtain the two separate libraries, one for CRISPRa and one for CRISPRi (Figure ED2A). Colony PCR of 47 bacterial clones revealed no undesired recombination in either library, and Sanger sequencing confirmed correct insertion of the expected sgRNAs (Figure ED2B). Guides were evenly represented, with no significant deviation from the expected median representation, ruling out strong biases during synthesis, amplification, or cloning (Figure ED2C).

For the generation of dual-sgRNA libraries, we adopted an arrayed strategy in which each target guide pair was ordered as primers and used separately to amplify the sgRNA scaffold together with the downstream promoter (Figure ED2D). The setup was chosen to prevent template-switching events observed when similar sequences were amplified and cloned in pooled formats[10]. Amplicons were pooled only after PCR amplification, digested to generate compatible ends, and ligated into the final CRISPRa and CRISPRi vectors. Colony PCR of 47 bacterial clones revealed no undesired recombination in either library, and Sanger sequencing confirmed correct insertion of the expected sgRNAs (Figure ED2E). Guides were evenly represented, with no significant deviation from the expected median representation, ruling out strong biases during synthesis, amplification, or cloning (Figure ED2E).

### Supplemental Methods

#### Plasmids

**pSpCas9n(BB)_R26-L** and **pSpCas9n(BB)_R26-R** *hROSA26* locus gRNA/Cas9n expression plasmids were previously described[23].

*AAVS1* locus ZFN expression plasmids and *CLYBL* locus TALEN expression plasmids were obtained from Addgene (**pZFN-*AAVS1*_ELD** #159297, **pZFN-*AAVS1*_KKR** #159298, **pZT-C13-L1** #62196, and **pZT-C13-R1** #62197).

**pR26-Bst_CAG-OPTtetR** was generated by excising the EGFPd2 cDNA of pR26-Bst_CAG-EGFPd2 using BamHI and MluI and replacing it with the OPTtetR cDNA from pR26-Neo_CAG-OPTtetR, cut with the same restriction enzymes.

**pCLYBL-Neo_CAG-dCAS9-2A-mCherry** was generated by cutting pC13N-iCAG.copGFP (Addgene, #66578) with BsrGI and MluI and inserting through an In-Fusion reaction: (1) cDNA of homo sapiens codon-optimized S. pyogenes catalytically inactive Cas9 (dCas9; containing D10A and H840A mutations) followed by HA tag, two SV40 nuclear localisation sequences, and a flexible linker, all amplified as a single fragment from pSLQ2312 (a kind gift of Jesse Zalatan) using primers 5'-CAAAGAATTGTGTACAACCATGGACAAGAAGTACAGCATCGG C-3' and 5'- CGCTGCTGCCGTTGCTCCCACTACCAATGCCGTCC-3'; (2) P2A-T2A-mCherry cDNA, obtained by synthesis (5'-GCAACGGCAGCAGCGGATCCGGAAGCGGAGCTACTAA CTTCAGCCTGCTGAAGCAGGCTGGAGACGTGGAGGAGAACCCTGGACCTGGAAGCGG AGAGGGCAGAGGAAGTCTGCTAACATGCGGTGACGTCGAGGAGAATCCTGGACCTATG GTGAGCAAGGGCGAGGAGGATAACATGGCCATCATCAAGGAGTTCATGCGCTTCAAGG TGCACATGGAGGGCTCCGTGAACGGCCACGAGTTCGAGATCGAGGGCGAGGGCGAGG GCCGCCCCTACGAGGGCACCCAGACCGCCAAGCTGAAGGTGACCAAGGGTGGCCCCC TGCCCTTCGCCTGGGACATCCTGTCCCCTCAGTTCATGTACGGCTCCAAGGCCTACGTG AAGCACCCCGCCGACATCCCCGACTACTTGAAGCTGTCCTTCCCCGAGGGCTTCAAGT GGGAGCGCGTGATGAACTTCGAGGACGGCGGCGTGGTGACCGTGACCCAGGACTCCT CCCTGCAGGACGGCGAGTTCATCTACAAGGTGAAGCTGCGCGGCACCAACTTCCCCTC CGACGGCCCCGTAATGCAGAAGAAGACCATGGGCTGGGAGGCCTCCTCCGAGCGGAT GTACCCCGAGGACGGCGCCCTGAAGGGCGAGATCAAGCAGAGGCTGAAGCTGAAGGA CGGCGGCCACTACGACGCTGAGGTCAAGACCACCTACAAGGCCAAGAAGCCCGTGCA GCTGCCCGGCGCCTACAACGTCAACATCAAGTTGGACATCACCTCCCACAACGAGGAC TACACCATCGTGGAACAGTACGAACGCGCCGAGGGCCGCCACTCCACCGGCGGCATG GACGAGCTGTACAAGTAATCTAGATAAACGCGTAGCTCGCTGATCAG-3').

sgRNA scaffold sequences were modified to extend two internal stem loop sequences to accommodate for MS2 or PP7 aptamers, or to append a com hairpin at the end of the sgRNA scaffold, in line with previously described designs[25,26]. The 10X Genomics capture sequence 2 (cs2) was attached at the 3' in MS2 and PP7 designs, while the 10X capture sequence 1 (cs1) was incorporated on an internal stem loop in com designs.

**pAAV-Puro_ClRIa-MS2_GFP** was constructed in two steps. First, the H1-TO_sgRNA cassette of pAAV-Puro_siKO was cut with BstBI and HincII, and replaced through an NEBuilder Hi-Fi reaction with: (1) U6 promoter containing a tet operon (TO) after the TATA box, obtained by synthesis (5’-GTGGGCTCTATGGGTCAATTCGGGTACCGAGGGCCTATTTCCCATGATTCCTTCATATTTGCATATACGATACAAGGCTGTTAGAGAGATAATTAGAATTAATTTGACTGTAAACACAAAGATATTAGTACAAAATACGTGACGTAGAAAGTAATAATTTCTTGGGTAGTTTGCAGTTTTAAAATTATGTTTTAAAATGGACTATCATATGCTTACCGTAACTTGAAAGTATTTCGATTTCTTGGCTTTATATATCTCCCTATCAGTGATAGAGACC-3’); (2) S. pyogenes sgRNA scaffold with 2 internal MS2 sites, PCR-amplified from pJF097.6 (a kind gift of Jesse Zalatan) using primers 5’-CCCTATCAGTGATAGAGACCGGGTGCAGGTGGACTGACTCACCTGCACCTGTTTAAGAGCTAGGCCAACATG-3’ And 5’-GATTACTATTAATAACTAGGACGGTATCGATAAAAAAACCTTAGCCGCTAATAGGTGAGCGCACCGACTCGGTGCC-3’, thus inserting a pair of upstream AarI cut sites and a downstream cs2. Second, the OPTtetR cDNA was cut from resulting vector with MreI and MluI, and replaced through an In-Fusion reaction with: (1) a PCR product from pAAV-Puro_siKO to reintroduce CAG intron with primers 5’-GCGCGGCGGCCCCCGGAGCGCCGGCGGCTGTCGAGGCGCG-3’ and 5’-GGTGGCGGCTTAATTAAATGATCCGAATTAATTCTTTGCCAAAATGATG-3’; (2) SV40 NLS and MCP cDNA, PCR amplified from pHRK090 with primers 5’-AATTCGGATCATTTAATTAAGCCGCCACCATGCCCAAAAAGAAAAGAAA-3’ and 5’-CCACCAGCGCTTGGCGCGCCACCGCTGCCGCTACCGTAGA-3’; (3) SV40 NLS, p65-HSF1 and P2A-T2A cDNA, obtained by synthesis (5’-GGCGCGCCAAGCGCtGGTGGAGGAGGATCTGGTGGAGGAGGATCTGGTGGGGGCGGCTCTGGCGGAGGAGGCAGTGGTCCCAAGAAGAAACGCAAGGTGGCAGCAGCGGGTTCCCCTTCCGGGCAAATATCAAATCAAGCCCTCGCTTTGGCCCCTTCTAGTGCGCCGGTCCTGGCCCAAACTATGGTCCCCAGTTCTGCTATGGTTCCACTTGCCCAACCGCCAGCCCCAGCACCAGTGCTTACGCCGGGTCCGCCTCAAAGCTTGAGTGCTCCGGTCCCAAAATCCACTCAAGCGGGCGAGGGAACTTTGTCAGAAGCATTGCTGCATCTCCAATTTGATGCGGATGAAGACCTCGGGGCACTGCTTGGTAACAGCACGGACCCCGGAGTCTTTACGGATCTTGCTTCCGTGGACAATAGTGAGTTTCAACAACTTCTCAATCAAGGCGTTTCCATGTCCCACAGCACTGCGGAGCCAATGCTCATGGAATACCCTGAAGCGATAACACGGCTCGTCACTGGGTCCCAGAGACCTCCTGATCCTGCTCCGACGCCCCTTGGGACCAGTGGTCTCCCGAATGGTCTCTCAGGTGACGAGGACTTCTCATCTATTGCTGATATGGATTTCTCTGCACTCCTTAGTCAAATCTCAAGCTCCGGGCAGGGTGGTGGGGGCTCTGGTTTTTCCGTTGACACTAGCGCTCTCTTGGATCTCTTCTCCCCCAGCGTGACAGTCCCGGATATGAGTCTCCCGGATCTCGATAGCTCTCTTGCCTCTATACAAGAACTCCTCAGCCCGCAGGAACCACCCAGGCCTCCAGAAGCTGAAAATTCTTCCCCCGATTCCGGTAAACAATTGGTCCACTACACGGCGCAACCGTTGTTCCTGCTTGATCCCGGATCTGTGGATACGGGTTCAAACGACCTCCCCGTTCTGTTTGAGCTGGGGGAGGGATCATATTTAGCGAGGGTGACGGGTTCGCGGAAGACCCGACAATTAGTTTGTTGACAGGTAGTGAACCCCCGAAGGCCAAGGATCCGACAGTTTCATGCACTGGTTCAGGGGATATCGCCACAAACTTCTCTCTTCTTAAGCAGGCGGGTGATGTTGAGGAGAACCCGGGACCAGGCTCTGGGGAAGGGAGAGGAAGTCTGCTCACATGTGGCGACGTAGAGGAAAATCCAGGGCCT-3’); (4) superfolder GFP, PCR amplified from pSLQ4352 (a kind gift of Jesse Zalatan) using primers 5’-TAGAGGAAAATCCAGGGCCTAGCAAAGGAGAAGAACTTTT-3’ and 5’-GCTGATCAGCGAGCTACGCGTTTATCTAGAATTTAAATCTATTTGTAGAGCTCATCCATGCC-3’.

The versions of pAAV-Puro_CIRIa-MS2_GFP where cs2 was substituted with cs1 or removed, were created by excising U6-TO-sgRNA cassette of pAAV-Puro_CIRIa-MS2_GFP with KpnI and ClaI and replacing it through a NEBuilder Hi-Fi with similar cassettes obtained by PCR of pAAV-Puro_CIRIa-MS2_GFP with the common primer 5’-CTATGGGTCAATTCGGGTAC-3’ and either primer 5’-GATTACTATTAATAACTAGGACGGTATCGATAAAAAAATTGCTAGGACCGGCCTTAAAGCGCACCGACTCGGTGCC-3’, to swap cs2 with cs1, or primer 5’-GATTACTATTAATAACTAGGACGGTATCGATAAAAAAAGCACCGACTCGGTGCC-3’, to remove cs2.

**pAAV-Puro_CIRIa-PP7_GFP** was obtained in two steps. First, the NLS-MCP cDNA of pAAV-Puro_CIRIa-MS2_GFP was excised with PacI and AscI and substituted through a NEBuilder Hi-Fi reaction with SV40 NLS and PP7 cDNA PCR amplified from pJZC43 (a kind gift of Jesse Zalatan) using primers (5’-GGCAAAGAATTAATTCGGATCATTTAATTAAGCCGCCACCATGCCCAAAAAG-3’ and 5’-CAGATCCTCCTCCACCAGCGCTTGGCGCGCCACCGCTTCCGGAGCCACGGCCCAGCGG-3’). Second, the U6-TO-sgRNA cassette of the resulting intermediate plasmid was excised with KpnI and ClaI and substituted through a NEBuilder Hi-Fi reaction with: (1) the aforementioned U6-TO synthetic fragment; (2) S. pyogenes sgRNA scaffold with two internal PP7 aptamers and two upstream AarI sites, obtained by synthesis (5’-GGGTGCAGGTGGACTGACTCACCTGCACCTGTTTAAGAGCTAGGCCACATAAGGAGTTTATATGGAAACCCTTATGCTGCAGGGCCTAGCAAGTTTAAATAAGGCTAGTCCGTTATCAACTTGGCCACATAAGGAGTTTATATGGAAACCCTTATGCTGCAGGGCCAAGTGGCACCGAGTCGGTGC-3’) and PCR amplified with the common primer 5’-CCCTATCAGTGATAGAGACCGGGTGCAGGTGGACTGACTCACCTGC-3’ and either primer 5’-GATTACTATTAATAACTAGGACGGTATCGATAAAAAAACCTTAGCCGCTAATAGGTGAGCGCACCGACTCGGTGCC-3’, to add cs2 at the 3’ end followed by a Pol III terminator, or primer 5’-GATTACTATTAATAACTAGGACGGTATCGATAAAAAAAGCACCGACTCGGTGCC-3’, to only add the Pol III terminator.

**pAAV-Puro_CIRIa-MS2_BFP** was built by excising sfGFP from pAAV-Puro_CIRIa-MS2_GFP with EcoRV and SwaI and replacing it through a NEBuilder Hi-Fi with (1) P2A-T2A site PCR amplified from pAAV-Puro_CRISPRa-GFP with primers 5’-CAGTTTCATGCACTGGTTCAGGGGAT-3’ and 5’-CATGTTCTCCTTAATCAGCTCGTCAGGCCCTGGATTTTCCTCTACGTC-3’ and (2) TagBFP, PCR amplified from CRISPRoff-v2.1 (Addgene #167981) with primers 5’-GACGTAGAGGAAAATCCAGGGCCTAGCGAGCTGATTAAGGAGAACATG-3’ and 5’-CGAGCTACGCGTTTATCTAGAATTTAAATCTAATTAAGCTTGTGCCCCAGTTTGC-3’ creating appropriate overlaps.

**pAAV-Puro_CIRIi-com_GFP** was obtained in two steps. First, the NLS-MCP-NLS-p65-HSF1 cDNA of pAAV-Puro_CIRIa-MS2_GFP was excised with PacI and EcoRV and substituted through a NEBuilder Hi-Fi reaction with a synthetic fragment encoding SV40 NLS and *Homo sapiens* codon-optimized Com-*KRAB* (5’-GCCGCCACCATGCCCAAAAAGAAGCGGAAAGTCGGCAGCATGAAGTCCATCCGGTGCAAGAACTGCAACAAGCTGCTGTTCAAGGCCGACAGCTTCGACCACATCGAGATCAGATGCCCCAGATGCAAGCGGCACATCATCATGCTGAACGCCTGCGAGCACCCCACCGAGAAGCACTGTGGAAAGAGAGAGAAGATCACCCACAGCGACGAGACAGTCAGATACGGCAGCACAAGCGGCCACAAGCTTAATGGCGGAGGCGGAGGAATGGACGCCAAGTCTCTTACCGCCTGGTCTAGAACCCTGGTCACCTTCAAGGACGTGTTCGTGGACTTCACCCGGGAAGAGTGGAAGCTGCTGGATACAGCCCAGCAGATCGTGTA

CCGGAACGTGATGCTGGAAAACTACAAGAACCTGGTGTCCCTGGGCTACCAGCTGACC AAGCCTGACGTGATCCTGCGGCTGGAAAAGGGCGAAGAACCT-3'), PCR amplified with primers 5'-GGCAAAGAATTAATTCGGATCATTTAATTAAGCCGCCACCATGCCCAAAAAG-3' and 5'-TAAGAAGAGAGAAGTTTGTGGCGATATCAGGTTCTTCGCCCTTTTCCAGC-3' to insert appropriate overlaps. Secondly, the U6-TO-sgRNA cassette of the resulting intermediate plasmid was excised with KpnI and ClaI and substituted through a NEBuilder Hi-Fi reaction with: (1) the aforementioned U6-TO synthetic fragment; (2) S. pyogenes sgRNA scaffold with internal cs1 from 10X Genomics and com aptamer on the 3', obtained by synthesis (5'-GTTTA AGAGCTATGCTGGAAACAGCATAGCAAGTTTAAATAAGGCTAGTCCGTTATCAACTTGGC CGCTTTAAGGCCGGTCCTAGCAAGGCCAAGTGGCACCGAGTCGGTGCCTGAATGCCTG CGAGCATC-3').

The versions of pAAV-Puro_CIRIi-com_GFP where cs1 was substituted with cs2 or removed, were created by excising the U6-TO-sgRNA cassette of pAAV-Puro_CIRIi-com_GFP with KpnI and ClaI and replacing it through a NEBuilder Hi-Fi reaction with similar cassettes composed of: (1) the aforementioned U6-TO synthetic fragment; (2) S. pyogenes sgRNA scaffold with internal cs2 from 10X Genomics and com aptamer on the 3', obtained by synthesis ( 5'-TTTA AGAGCTATGCTGGAAACAGCATAGCAAGTTTAAATAAGGCTAGTCCGTTATCAACTTGGC CGCTCACCTATTAGCGGCTAAGGGGCCAAGTGGCACCGAGTCGGTGCCTGAATGCCTG CGAGCATC-3'), or the same cassette without CS (5'-TTTAAGAGCTATGCTGGAAACAGCA TAGCAAGTTTAAATAAGGCTAGTCCGTTATCAACTTGGCCGGCCAAGTGGCACCGAGTC GGTGCCTGAATGCCTGCGAGCATC-3').

**pAAV-Hygro_CIRIi-com** was obtained in two steps. First, the *AAVS1* 5' homology arm and the Puromycin resistance cassette of pAAV-Puro_CIRIi-com_GFP were excised with PmeI and substituted by a NEBuilder Hi-Fi reaction with: (1) *AAVS1* 5' homology arm PCR amplified from pAAV-Puro_CIRIi-com_GFP with primers 5'-CCCTCACTAAAGGGACTAGTCCTGCAG-3' and 5'-GGCTTTTTCATCTCGAGCCTAGGGCCGG-3' and (2) Hygroycin resistance cassette PCR amplified from PGK plasmid (Addgene #169744) with primers 5'-GCCCTAGGCTCGAGATGA AAAAGCCTGAACTCACCGCG-3' and 5'-CACAGTCGAGGCTGATCAGCGGGTTTCCTATT CCTTTGCCCTCGGAC-3' to insert appropriate overlaps. Secondly, the sfGFP cassette was excised with EcoRV and SwaI and replaced through a NEBuilder Hi-Fi with a synthetic cDNA 5'-GGGCGAAGAACCTGATATCtAGATTTAAATTCTAGATAAACGCGTAGC-3' to only add the stop codon.

**pAAV-Puro_CIRIa-MS2_BFP_E2-MYOD1x2** and **pAAV-Hygro_CIRIi-com_E2-OSN**, were generated by excising the U6-TO cassette of the first empty guide RNA with KpnI and AarI and replacing it through a NEBuilder Hi-Fi reaction with similar cassette containing the H1-TO promoter PCR amplified from pAAV-Puro_siKO with primers 5'-GTGGGCTCTATGGGTCAATTCG–3' and 5'-CATGTTGGCCTAGCTCTTAAACAGGTGCA–3'

A similar strategy was applied for **pAAV-Puro_CIRIa-MS2_BFP_E1-MYOD1x2** and **pAAV-Hygro_CIRIi-com_E1-OSN**, where the U6-TO was replaced by two fragments containing (1) H1 promoter and (2) a fragment to reintroduce AarI sites. The fragments were PCR amplified respectively from pAAV-Puro_siKO with same primer forward and primer reverse 5'- GGGAAC

TTATAAGATTCCCAAATCCAAAG–3’ and from pAAV-Puro_CRISPRa-BFP with primers 5’-GGGAATCTTATAAGTTCCCGGGTGCAGGTGGAC–3’ and same primer reverse.

Vectors carrying multiple sgRNAs (**pAAV-Puro_CIRIa-MS2_BFP_MYOD1x3**, **pAAV-Hygro_CIRIi-com_OSN**, and their corresponding controls **pAAV-Puro_CIRIa-MS2_BFP_NTCx3** and **pAAV-Hygro_CIRIi-com_ NTCx3**) were generated as described in the sgRNA molecular cloning method section (see STAR METHODS).

To generate the **single-sgRNA insert library**, oligonucleotides were designed to include TetO2 and the 20-nt protospacers, with the first nucleotide constrained to G or A to match the transcription-initiation preference of the H1 promoter. Oligo ends were flanked by BbsI recognition sites and designed as follows: 5’-BbsI recognition site I–TetO2–protospacer–BbsI recognition site III–PCR adapter-3’. The sequence was: 5’-TGGAAGACGTTCCCTATCAGTGATAGAGATCCCG--protospacer--GTTTGAGTCTTCGGAATGCC-3’. Oligo pools were purchased as IDT oPools 50 pmol, resuspended in nuclease-free water to obtain a 50 µM stock, and diluted 1:100 to a working concentration of 0.5 µM. For each PCR reaction, 2 µL of the diluted pool was used. PCR amplification was carried out using Q5 Hot Start High-Fidelity DNA Polymerase and the following primers (Eurofins Genomics): 5’-TGGAAGACGTTCCCTATCAG-3’ and 5’-GGCATTCCGAAGACTCAAAC-3’. PCR conditions were as follows: annealing temperature 55 °C, extension time 10 s, 15 cycles. Amplified products were purified using the NucleoSpin Gel and PCR Clean-up Kit and eluted in 20 µL of Buffer EB. Purified PCR products were analyzed on a TapeStation using High Sensitivity DNA ScreenTape to confirm the expected size and assess non-specific bands. Concentrations were determined using the Qubit dsDNA High Sensitivity Assay Kit. Purified PCR products were digested with BbsI-HF at 37 °C for 1 h to generate sticky ends suitable for directional ligation. Because the digested products were relatively short, they were resolved on a 20% Novex TBE polyacrylamide gel under low-voltage conditions (80 V for 4 h at 4 °C) in pre-chilled TBE buffer to minimize guide denaturation and preserve uniform representation. After electrophoresis, the gel was post-stained with SYBR Gold and the desired DNA fragment was excised. DNA was extracted from the gel by passive diffusion in a buffer containing 500 mM ammonium acetate, 10 mM magnesium acetate tetrahydrate, and 1 mM EDTA pH 8.0 in nuclease-free water. The gel slice was incubated overnight at 37 °C. DNA was precipitated by adding 2 volumes of ethanol and purified using the NucleoSpin Gel and PCR Clean-up Kit. The recipient vectors (pAAV-Puro_CIRIa-MS2_BFP_E2-MYOD1x2 and pAAV-Hygro_CIRIi-com_E2-OSN) were digested with PaqCI, a functional isoschizomer of AarI, resolved by agarose gel electrophoresis, and extracted using the QIAEX II Gel Extraction Kit. Ligation was performed using T4 DNA Ligase with a vector-to-insert molar ratio of 1:3 and incubated at room temperature for 15 min. Recombinant plasmids were transformed into NEB 5-alpha Competent *E. coli* (High Efficiency) following the manufacturer’s instructions. For quality control, colonies were picked and screened by colony PCR to exclude unwanted recombination events using the following primers: 5’-CGAACGCTGACGTCATCAACC-3’ and 5’-CTGTCACGAGACTAGCAGTTC-3’.

To generate the **dual-sgRNA insert library**, each protospacer was included in an oligonucleotide primer used to amplify the sgRNA scaffold and downstream promoter sequence. Primer ends were flanked by BbsI recognition sites to generate overhangs compatible with recipient vectors digested using PaqCI/AarI-compatible cloning sites. Oligonucleotides were designed as follows: 5'-BbsI recognition site I–protospacer–PCR-compatible end-3'. Specifically: forward primer 5'-TGGAAGACGTTCCCG–protospacer–GTTTAAGAGCTAGGCCAAC-3'; and reverse primer 5'-GCCGAAGACTCAAAC–protospacer–CGGTCTCTATCACTGATAGGG-3'. Protospacers in reverse primers were ordered as reverse-complement sequences. Oligonucleotides were purchased from IDT in plates as dry powder, resuspended, and diluted to the working concentration used for PCR amplification. PCR amplification was carried out using Q5 Hot Start High-Fidelity DNA Polymerase following the manufacturer's instructions, with annealing temperature 55 °C, extension time 10 s, and 15 cycles. A fraction of each amplified product was analyzed by agarose gel electrophoresis to confirm guide-specific amplification. The remaining reactions were pooled, purified using the Nucleospin Gel and PCR Clean-up Kit, and eluted in 20 µL of Buffer EB. Purified PCR products were analyzed on a TapeStation using High Sensitivity DNA ScreenTape to confirm the expected size and assess non-specific bands. Concentrations were determined using the Qubit dsDNA High Sensitivity Assay Kit. PCR products were digested with BbsI-HF at 37 °C for 1 h to generate sticky ends suitable for directional ligation. Digested products were resolved on a 1% agarose gel, and the desired DNA fragment was excised and purified using the QIAEX II Gel Extraction Kit. The recipient vectors (pAAV-Puro_CIRIa-MS2_BFP_E1-MYOD1x2 and pAAV-Hygro_CIRIi-com_E1-OSN) were digested with PaqCI, resolved by agarose gel electrophoresis, and purified using the QIAEX II Gel Extraction Kit. Ligation was performed using T4 DNA Ligase with a vector-to-insert molar ratio of 1:3 and incubated at room temperature for 15 min. Recombinant plasmids were transformed into NEB 5-alpha Competent *E. coli* (High Efficiency) following the manufacturer's instructions. For quality control, colonies were picked and screened by colony PCR to exclude unwanted recombination events using a common forward primer, 5'-CGAACGCTGACGTCATCAACC-3', together with a CRISPRa-specific primer in BLOCK1, 5'-GCTGTCAAGACACAGCATAG-3', or a CRISPRi-specific primer in BLOCK3, 5'-CTGTCACGAGACTAGCAGTTC-3'.

Plasmids for screening-hit validation **pAAV-Puro_CIRIa-MS2_BFP_SMARCD3x2-MYOD1x2**, **pAAV-Puro_CIRIa-MS2_BFP_KDM6Bx2-MYOD1x2**, and **pAAV-Hygro_CIRIi-com_ID3x2-OSN** were generated during pool-library construction, following the dual-guide protocol described above. Individual colonies were manually isolated from the pool for sequencing analysis and conserved as glycerol stocks, enabling recovery of single plasmids corresponding to each experimental condition.

### Supplemental Figures

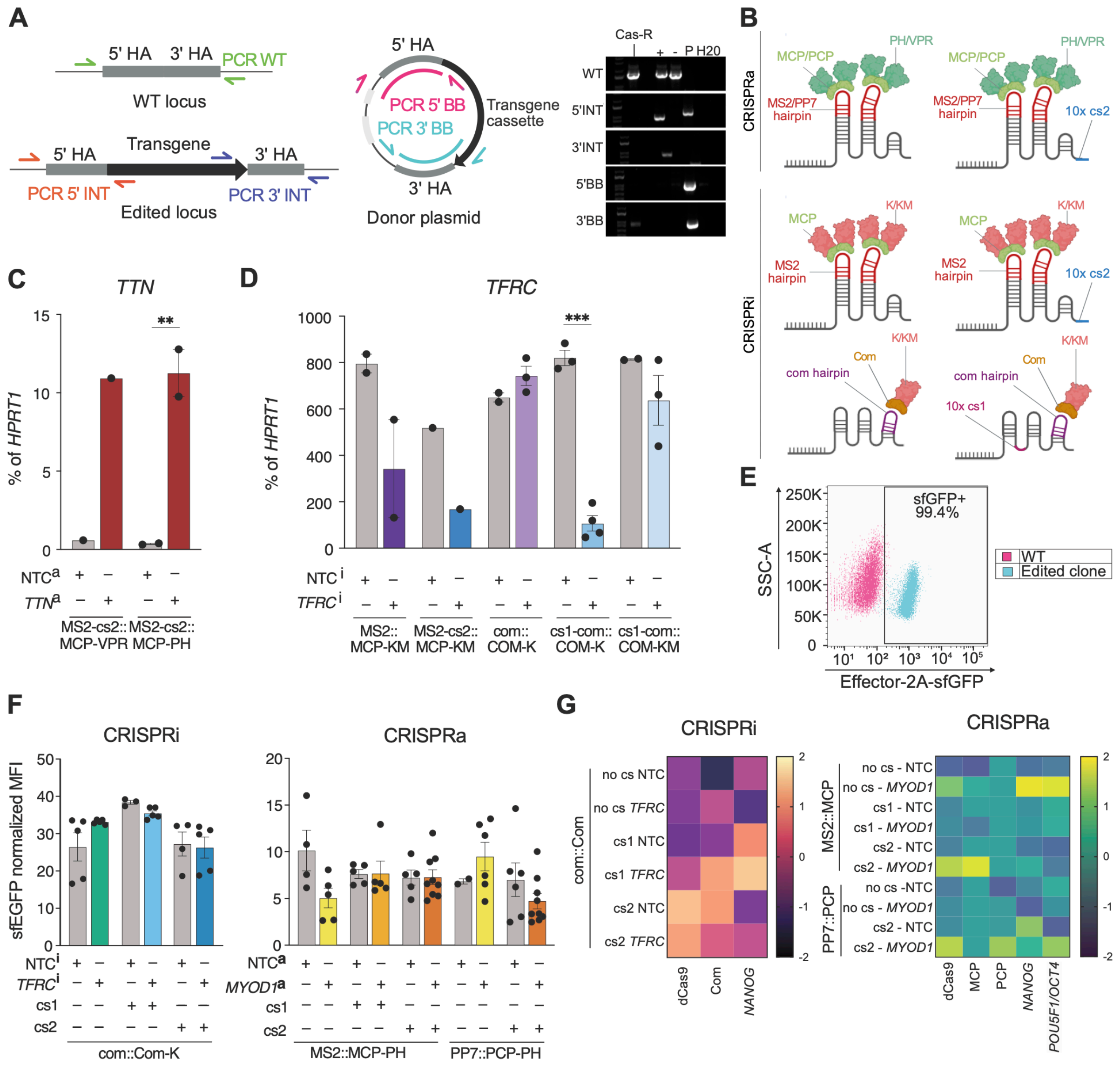


#### Supplemental Figure S1. Extended optimization of sgRNA scaffolds for orthogonal CRISPRa/i in hiPSCs

Legend on the next page

#### Supplemental Figure S1. Extended optimization of sgRNA scaffolds for orthogonal CRISPRa/i in hiPSCs (continued)

(A) Genotyping strategy and validation of dCas9 integration. Left, schematic of genomic PCR assays used to assess genome editing. Site-specific transgene integration was detected by 5'INT and 3'INT junctional PCRs; WT PCR assessed retention of the unedited allele; and 5'BB and 3'BB backbone PCRs were used to detect off-target random integration of the targeting plasmid. Right, genotyping PCRs for dCas9 targeting at the *CLYBL* locus. Cas9-R hiPSCs carried randomly integrated dCas9 rather than on-target *CLYBL* integration. "+" positive-control clone with on-target integration; "-" parental WTC11 hiPSCs; P, targeting plasmid control for backbone PCRs.

(B) Schematic of sgRNA-scaffold and RBP–effector combinations tested for CRISPRa and CRISPRi. MS2, PP7, and com hairpins were tested with their cognate MCP, PCP, and Com RNA-binding proteins, respectively, together with the indicated activation or repression domains and optional 10X Genomics capture sequences.

(C) RT-qPCR analysis of *TTN* expression after CRISPRa using MS2-cs2 sgRNAs recruiting either MCP-PH or MCP-VPR. PH, p65-HSF1; VPR, VP64-p65-Rta; NTC, non-targeting control. N = 1–2 clones; two-way ANOVA with activator and protospacer type as factors, followed by Holm–Sidak's multiple-comparison test for the *TTN versus* NTC protospacer within each activator. Here and throughout the figure * $p < 0.05$; ** $p < 0.01$; *** $p < 0.001$.

(D) RT-qPCR analysis of *TFRC* expression after CRISPRi using the indicated guide::effector architectures. K, KRAB; KM, KRAB-MeCP2; cs1/2, 10X Genomics capture sequences 1/2. N = 1–4 clones; two-way ANOVA with guide::effector architecture and protospacer type as factors, followed by Holm–Sidak's multiple-comparison test for *TFRC versus* NTC comparisons within each architecture.

(E) Representative flow cytometry plot showing homogeneous effector-2A-sfGFP expression in an edited clone compared with WT hiPSCs. Only clones with > 90% $sfGFP^+$ cells were selected for downstream analysis.

(F) Flow cytometry quantification of effector-2A-sfGFP mean fluorescence intensity (MFI) across CRISPRi and CRISPRa clones analyzed in Figure 1D and carrying the indicated capture sequences and recruitment systems. N = 2–9 clones; two-way ANOVA with guide::effector architecture and protospacer type as factors, followed by Holm–Sidak's multiple-comparison test for targeting *versus* NTC comparisons within each architecture, and all pairwise architecture comparisons within the same guide type. $p > 0.05$ for all comparisons.

(G) Heatmaps of RT-qPCR analysis for dCas9, RBP–effector components, and pluripotency markers in the CRISPRi and CRISPRa clones from panel F. Values are shown as Z-scores.

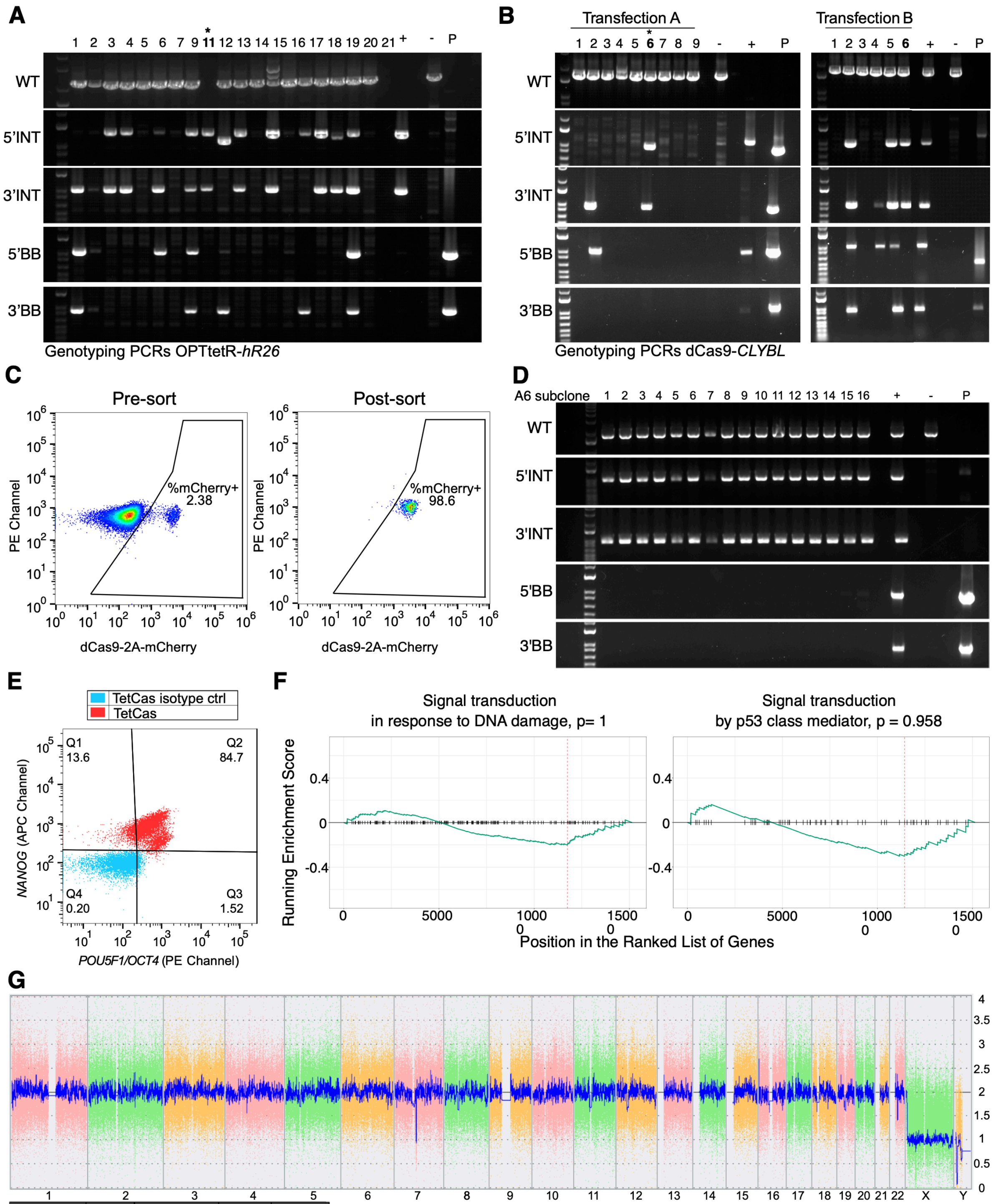


| Chromosome | Type | Cytoband Start | CN State | Size (kbp) |
|---|---|---|---|---|
| Y | Loss | p11.2 | 0.00 | 2,935 |

#### Supplemental Figure S2. Extended generation and validation of TetCas hiPSCs

Legend on the next page

#### Supplemental Figure S2. Extended generation and validation of TetCas hiPSCs (continued)

(A) Genotyping PCRs to verify OPTtetR integration at the *hROSA26* locus. The genotyping strategy is conceptually identical to that in Figure S1A: 5'INT and 3'INT junctional PCRs detect site-specific integration, WT PCR detects retention of the unedited allele, and 5'BB and 3'BB backbone PCRs detect off-target random integration of the targeting plasmid. Clone 11, marked with an asterisk, carried heterozygous *hROSA26* integration without additional random integrations and was selected to generate Tet hiPSCs. "-" parental WTC11 hiPSCs; "+" positive-control clone with on-target integration; P, targeting plasmid control for backbone PCRs.
(B) As in panel A, but to verify dCas9-2A-mCherry integration at the *CLYBL* locus in Tet hiPSCs. Results are shown for two independent transfections. Clone A6, marked with an asterisk, carried heterozygous *CLYBL* integration without additional random integrations and was selected for single-cell subcloning by FACS of mCherry$^{+}$ cells.
(C) Flow cytometry analysis of dCas9-2A-mCherry expression in clone A6 before and after single-cell sorting.
(D) As in panel B, but to verify dCas9-2A-mCherry integration at the *CLYBL* locus after mCherry$^{+}$ subcloning. Sixteen subclones were analyzed; all carried heterozygous *CLYBL* integration without detectable random integrations. Clone A6.10 was selected as the final TetCas hiPSC line for downstream experiments.
(E) Flow cytometry analysis of NANOG and POU5F1/OCT4 expression in TetCas hiPSCs, confirming maintenance of pluripotency-marker expression after sequential genome editing.
(F) Gene Set Enrichment Analysis (GSEA) from RNA-seq data comparing TetCas and parental WT hiPSCs (Figure 2G). N = 3 cultures; no significant enrichment was detected.
(G) Copy-number variation (CNV) analysis of TetCas hiPSCs by comparative genomic hybridization (CGH)-based molecular karyotyping. Whole-genome copy-number profile across autosomes and sex chromosomes is shown. Colored traces indicate raw probe signals for individual chromosomes, while the blue trace shows the normalized smoothed signal used to infer copy-number state. Values of 2, 3, and 1 correspond to normal diploid copy number, gain, and loss, respectively. Aberrations, when present, are indicated by red arrows. The chrY p11.2 loss corresponds to a known deletion present in the male WTC11 donor background. The small apparent haploid-state signal on chromosome 7 was later confirmed to be present in the parental WTC11 *TTN*-mEGFP hiPSCs used for this study (Figure S7A); no additional newly acquired large-scale CNVs were detected.

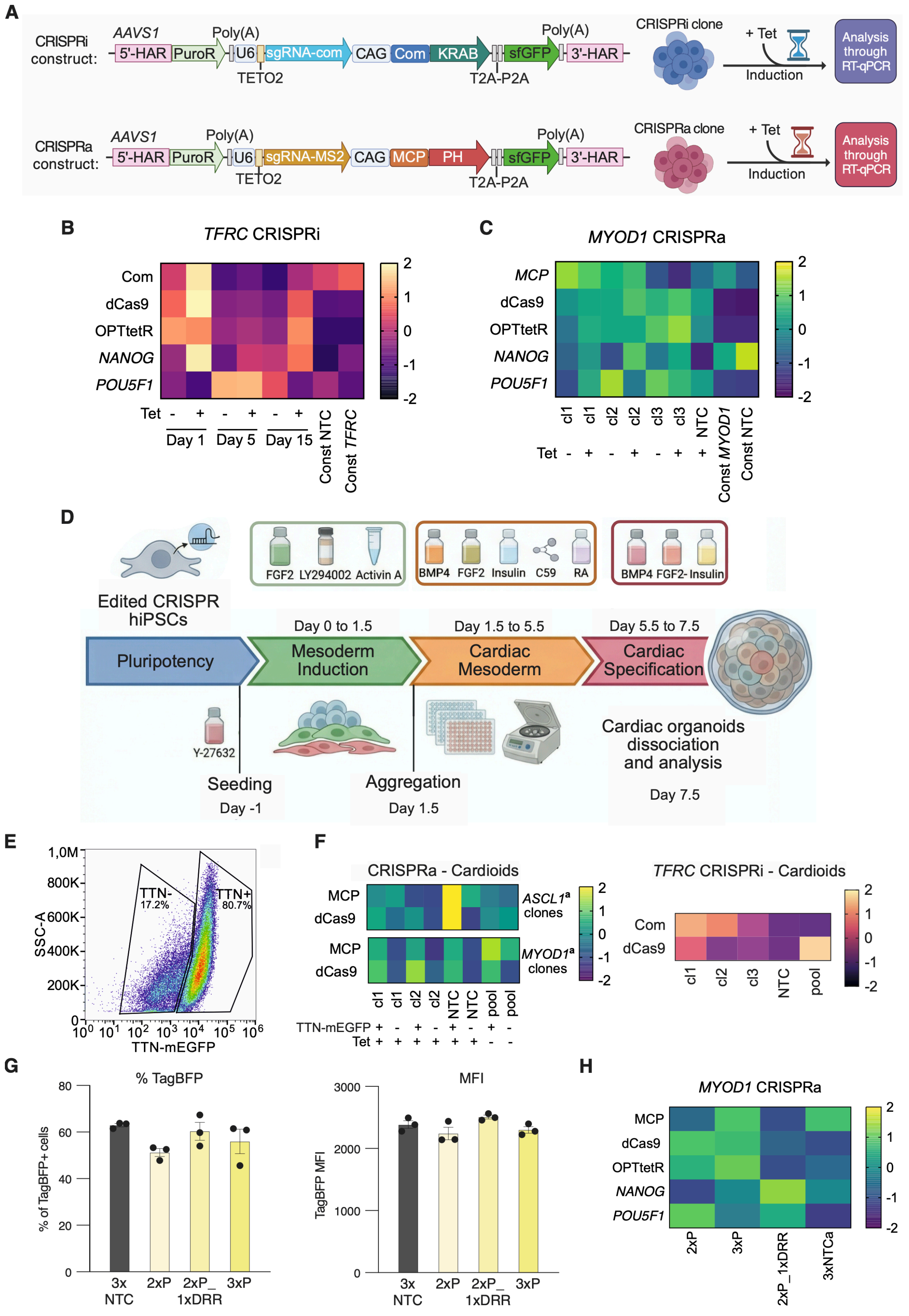


**Supplemental Figure S3. Extended validation of inducible CRISPRa/i in hiPSCs and cardiac organoids**

Legend on the next page

#### Supplemental Figure S3. Extended validation of inducible CRISPRa/i in hiPSCs and cardiac organoids (continued)

(A) Workflow for inducible CRISPRa/i validation in TetCas hiPSCs. Cells were re-targeted at the *AAVS1* locus with the selected CRISPRi or CRISPRa cassettes, each encoding a Tet-inducible sgRNA scaffold and a constitutively expressed RBP–effector module. The workflow is analogous to Figure 1C, except that sgRNA expression is Tet-inducible and cells were treated with tetracycline before RT-qPCR analysis.

(B) Heatmap of RT-qPCR analysis for CRISPRi system components and pluripotency markers in the *TFRC* CRISPRi clones analyzed in Figure 3A, compared to constitutive (const.) *TFRC* CRISPRi and matched NTC controls from Figure 1D. Values are shown as Z-scores.

(C) Heatmap of RT-qPCR analysis for CRISPRa system components and pluripotency markers in the *MYOD1* CRISPRa clones analyzed in Figure 3C, compared to constitutive (const.) *MYOD1* CRISPRa and matched NTC controls from Figure 1D. Values are shown as Z-scores.

(D) Schematic of the cardiac organoid (cardioid) differentiation protocol. Edited CRISPRa/i hiPSCs were differentiated in 3D using small molecule-guided aggregation[38], then dissociated and analyzed at day 7.5.

(E) Representative flow cytometry gating of day 7.5 cardiac organoids, showing TTN-mEGFP$^{+}$ cardiomyocyte and TTN-mEGFP$^{-}$ non-cardiomyocyte populations used for sorted RT-qPCR analysis.

(F) Heatmaps of RT-qPCR analysis for CRISPRa and CRISPRi system components in day 7.5 cardiac organoids from the experiments described in Figures 3F–3G. Values are shown as Z-scores.

(G) Flow cytometry quantification of MCP-PH-2A-TagBFP expression in TetCas hiPSCs re-targeted at *AAVS1* with multiplex inducible CRISPRa cassettes (Figures 3H–3I). Data are from three independently edited pools selected with puromycin and analyzed without clonal isolation. 3xNTC, three non-targeting control sgRNAs; 2xP, two promoter guides; 3xP, three promoter guides; 2xP_1xDRR, two promoter guides plus one distal regulatory region (DRR) guide. N = 3 independently edited pools; one-way ANOVA followed by Holm–Sidak's multiple-comparison test for all pairwise comparisons. $p > 0.05$.

(H) Heatmap of RT-qPCR analysis for CRISPRa system components and pluripotency markers in multiplex *MYOD1* CRISPRa pools from the experiment described in panel G. Values are shown as Z-scores.

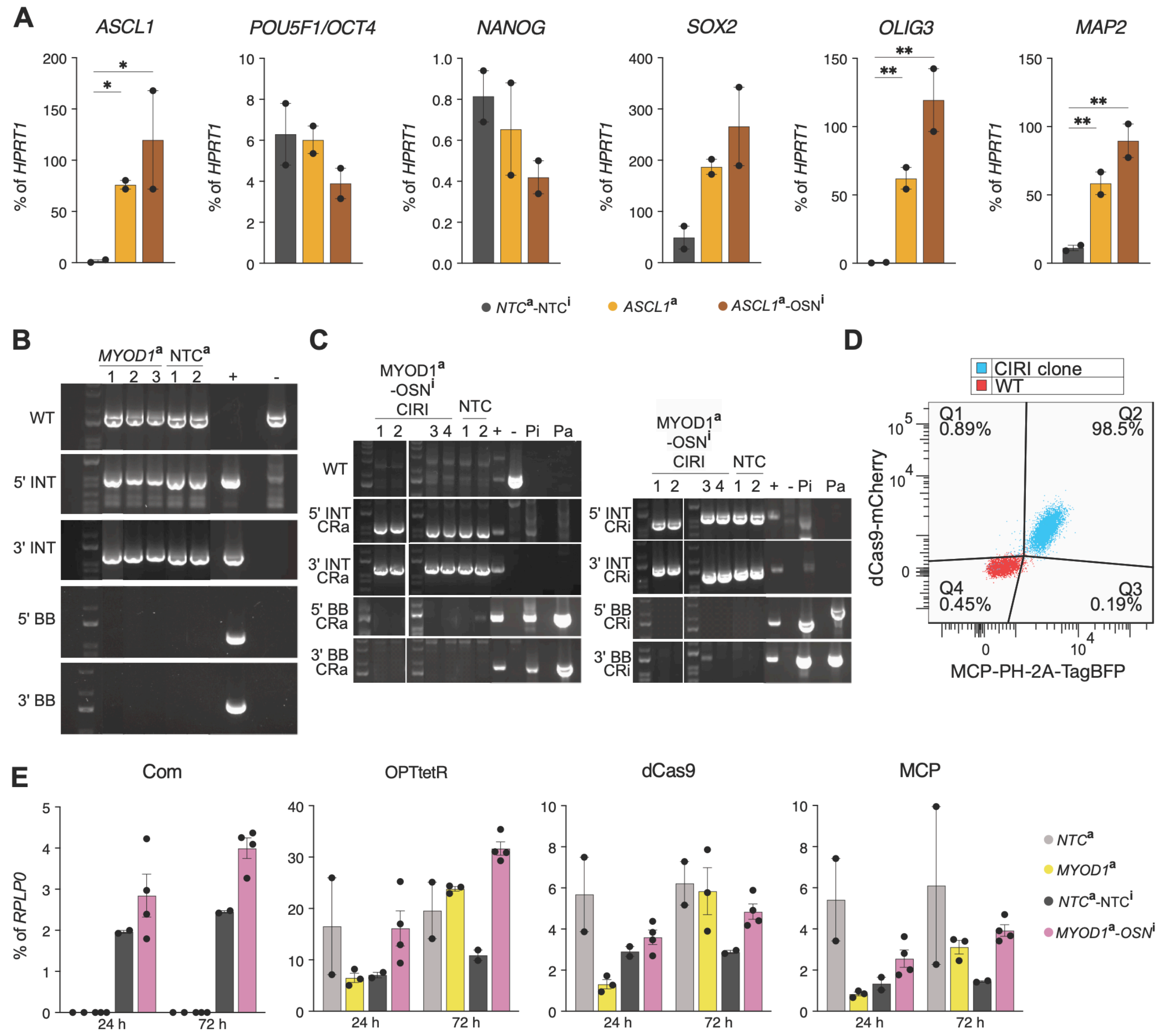


#### Supplemental Figure S4. Extended validation of combinatorial CRISPRa/i in hiPSCs

Legend on the next page

#### Supplemental Figure S4. Extended validation of combinatorial CRISPRa/i in hiPSCs (continued)

(A) RT-qPCR analysis of *ASCL1*, pluripotency markers, and early neurogenic markers *OLIG3* and *MAP2* in *ASCL1*$^{a}$-*OSN*$^{i}$ CIRI, *ASCL1* CRISPRa-only, and NTC control clones after 14 days of Tet-induction in neuronal-permissive media, compared to NTC controls. N = 2 clones; one-way ANOVA followed by Holm–Sidak's multiple-comparison test for all pairwise comparisons. * $p < 0.05$; ** $p < 0.01$; $p > 0.05$ not reported.

(B) Genotyping PCRs to verify *AAVS1* integration of the 3xP *MYOD1* inducible CRISPRa cassette or 3xNTC control cassette in TetCas hiPSCs. The strategy is conceptually identical to that in Figure S1A, and all selected clones carried homozygous *AAVS1* integration without additional random integrations. "-" parental WTC11 hiPSCs; "+" positive-control clone with on-target and off-target integrations.

(C) As in B, but to verify *AAVS1* integration in *MYOD1*$^{a}$-*OSN*$^{i}$ CIRI and matched NTC control clones. Compound heterozygous clones carrying both CRISPRa and CRISPRi cassettes integrated at *AAVS1*, without additional random integrations, were identified using cassette-specific INT and BB PCRs. CRi/a, CRISPRi/a cassette; Pi/a, CRISPRi/a targeting plasmid.

(D) Representative flow cytometry analysis confirming homogeneous MCP-PH-2A-TagBFP and dCas9-2A-mCherry expression in a CIRI clone, compared with WT hiPSCs.

(E) RT-qPCR analysis of CRISPRa/i system-component expression in the clones analyzed in Figures 4B–4D, including *MYOD1*$^{a}$-*OSN*$^{i}$ CIRI, *MYOD1* CRISPRa-only (*MYOD1*$^{a}$), and matched NTC controls after 24 h and 72 h of Tet treatment. This panel validates expression of the CRISPRa/i chassis and effector modules; no statistical analysis was performed.

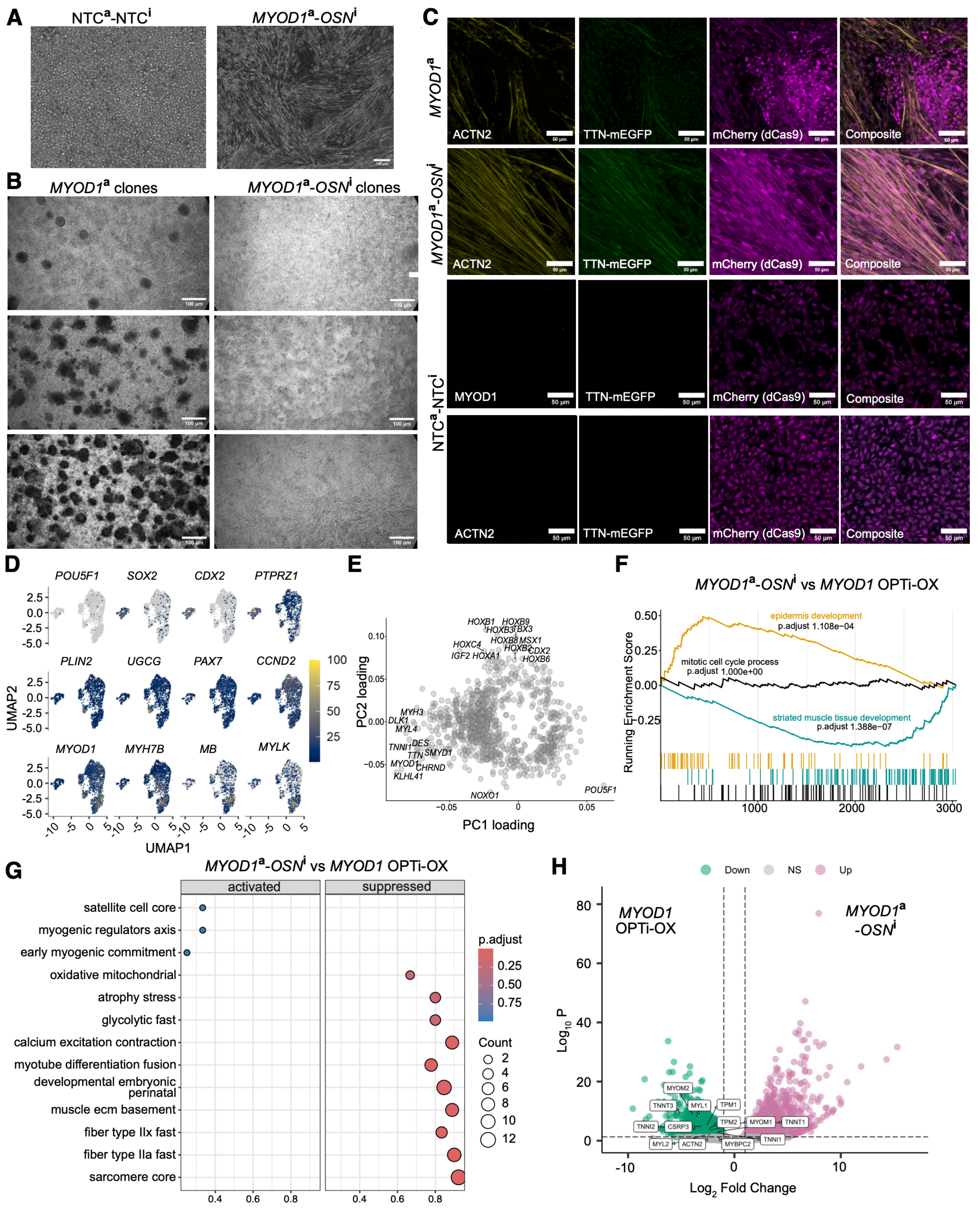


#### Supplemental Figure S5. Extended characterization of CIRI-mediated myogenic forward programming

Legend on the next page

#### Supplemental Figure S5. Extended characterization of CIRI-mediated myogenic forward programming (continued)

(A) Representative phase-contrast images after 7 days of culture in Tet-containing myogenic medium, comparing *MYOD1*$^{a}$-*OSN*$^{i}$ CIRI and matched NTC clones. CIRI cells display elongated mesenchimal morphology, whereas NTC controls retain a compact epithelial morphology. Scale bar, 100 µm.
(B) Representative low-magnification phase-contrast images after 7 days of myogenic differentiation for three *MYOD1*$^{a}$-*OSN*$^{i}$ CIRI and three *MYOD1*$^{a}$ CRISPRa-only clones. CIRI clones show more homogeneous differentiation, whereas CRISPRa-only clones retain misdifferentiated regions visible as darker foci. Scale bars, 100 µm.
(C) Confocal microscopy after 7 days of myogenic differentiation, showing ACTN2 immunostaining, TTN-mEGFP reporter activation, and maintenance of dCas9-2A-mCherry expression in representative *MYOD1*$^{a}$-*OSN*$^{i}$ CIRI and *MYOD1*$^{a}$ CRISPRa-only clones. CIRI NTC controls stained for MYOD1 or ACTN2 are shown as negative controls (also relative to Figure 5B). Composite images show ACTN2 and mCherry reporter, except for the third row from the top, reporting MYOD1 and mCherry signal. Scale bars, 50 µm.
(D) Feature plots showing expression of representative marker genes used to annotate the transcriptional states identified in the single-cell RNA-seq analysis shown in Figure 5E.
(E) Gene loadings on PC1 and PC2 from the bulk RNA-seq principal component analysis shown in Figure 5F. PC1 separates *MYOD1*$^{a}$-*OSN*$^{i}$ CIRI and *MYOD1* OPTi-OX samples after 7 days of differentiation, enriched for myogenic programs, from Tet treatment-matched NTC control samples and hiPSCs, enriched for pluripotency-associated genes.
(F) Gene Set Enrichment Analysis (GSEA) comparing *MYOD1*$^{a}$-*OSN*$^{i}$ CIRI and *MYOD1* OPTi-OX cells, showing enrichment of representative developmental, cell cycle, and muscle-related gene sets.
(G) As in F, but for additional muscle differentiation-related Gene Ontology terms.
(H) Volcano plot of bulk RNA-seq differential expression between *MYOD1*$^{a}$-*OSN*$^{i}$ CIRI and *MYOD1* OPTi-OX cells. Selected genes involved in sarcomere formation and muscle maturation are labeled.

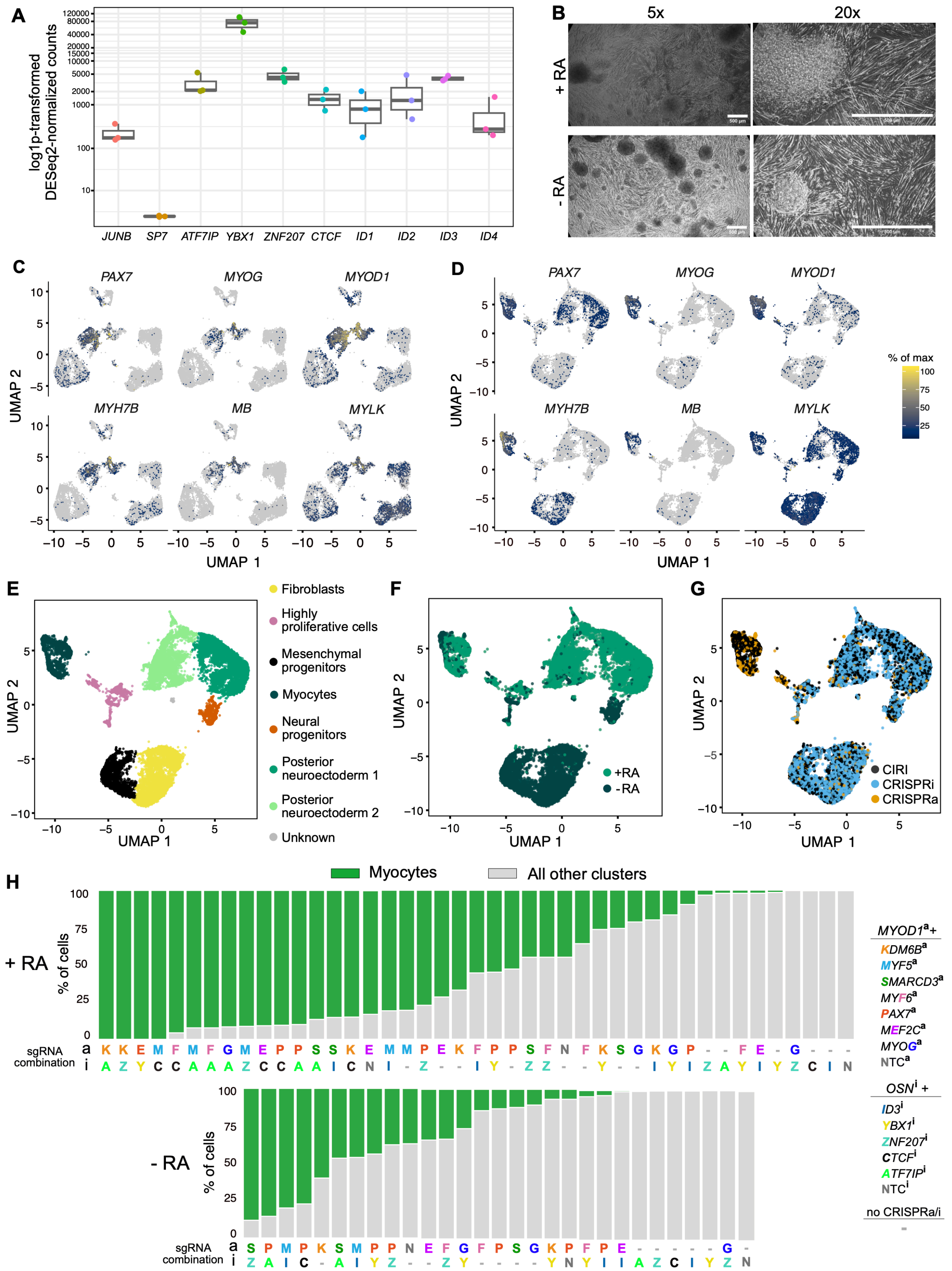


### Supplemental Figure S6. Extended characterization of pooled combinatorial CRISPRa/i screening

Legend on the next page

#### Supplemental Figure S6. Extended characterization of pooled combinatorial CRISPRa/i screening (continued)

(A) Expression in hiPSCs of candidate forward-programming roadblock genes considered for the pooled CRISPRi screen, shown as DESeq2-normalized counts on a log scale. N = 2 cultures. *JUNB*, *SP7*, *ATF7IP*, and *ZNF207* were considered based on Missinato *et al.*[50]; *JUNB* and *SP7* were not included in the final library because of low expression in hiPSCs. Among the ID-family candidates, *ID3* was selected because it was the predominant expressed family member, whereas *ID1*, *ID2*, and *ID4* showed lower expression.
(B) Representative phase-contrast images of single-guide pooled CIRI cells after 7 days of myogenic differentiation in medium with (+RA) or without (-RA) retinoic acid. Images at 5x and 20x magnification. Scale bars, 500 µm.
(C) Feature plots showing expression of representative marker genes used to annotate the transcriptional states for the dual-sgRNA CIRI screen described in Figure 6D. Expression is shown as percentage of maximum expression.
(D) As in panel C, but for the transcriptional states identified in panel E.
(E) UMAP of the single-guide pooled CIRI screen after 7 days of myogenic differentiation. Cells were clustered by single-cell RNA-seq and annotated based on marker-gene expression (panel D).
(F) As in panel E, colored by differentiation condition (+RA or -RA).
(G) As in panel E, colored by assigned perturbation modality: CRISPRa-only, CRISPRi-only, or both (CIRI).
(H) Quantification of the fraction of assigned cells adopting a myocyte fate for each combinatorial perturbation in the single-guide CIRI screen in +RA and -RA conditions. Green indicates myocytes; grey indicates all other transcriptional clusters. Bars are ranked by myocyte fraction within each condition, highlighting the strongest myocyte-enriching sgRNA combinations at the left. “-” indicates no assigned CRISPRa/i perturbation.

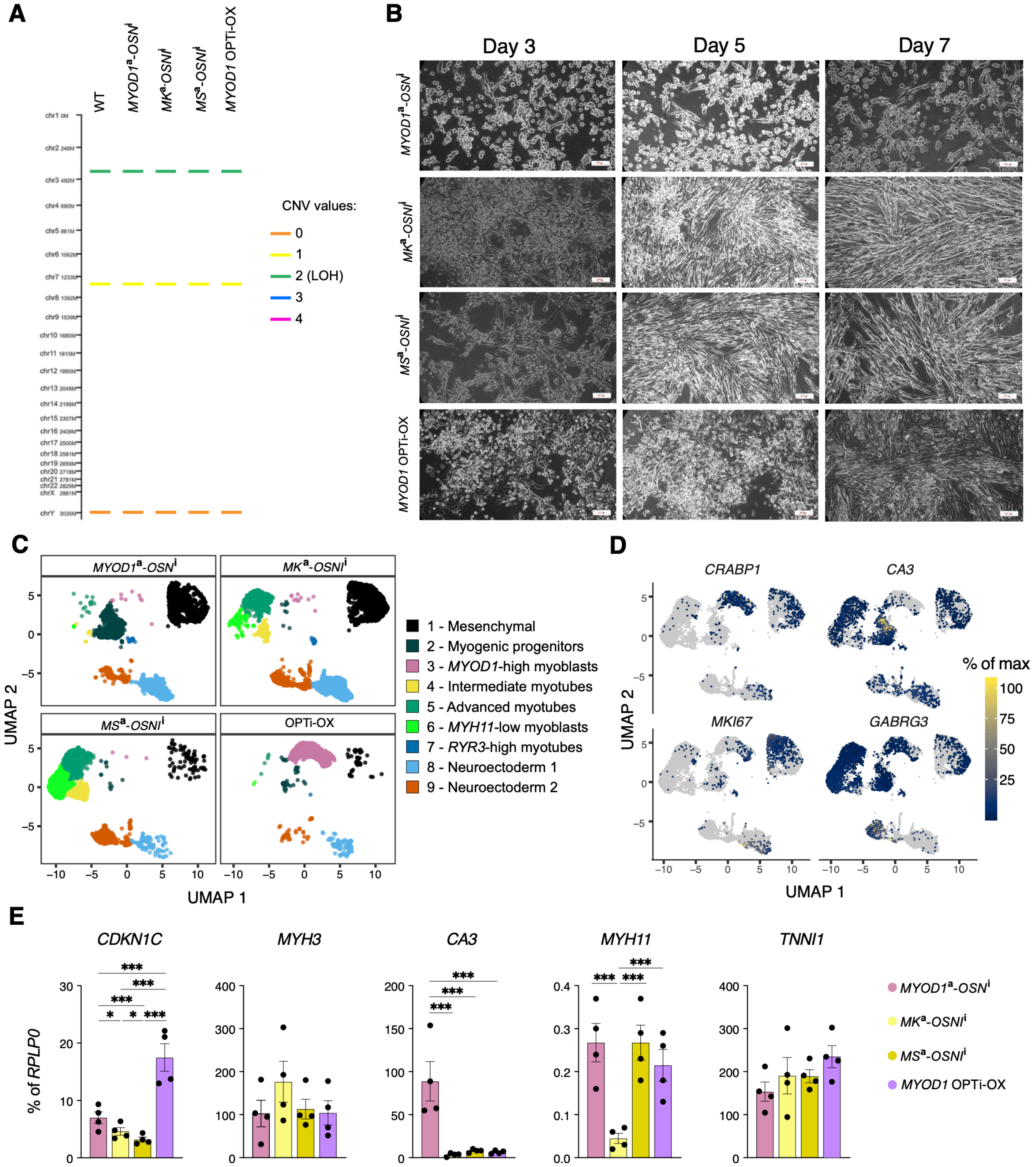


#### Supplemental Figure S7. Extended validation of enhanced retinoic acid-independent myogenic programming

Legend on the next page

#### Supplemental Figure S7. Extended validation of enhanced retinoic acid-independent myogenic programming (continued)

(A) Copy-number variation (CNV) analysis of parental WTC11 hiPSCs, CIRI lines, and the *MYOD1* OPTi-OX benchmark line. Colored segments indicate detected CNVs and copy-number-neutral loss of heterozygosity (LOH), with inferred copy-number states shown in the legend.

(B) Representative phase-contrast images showing the time course of myogenic forward programming at days 3, 5, and 7 in retinoic acid-free (-RA) myogenic medium. Baseline *MYOD1*$^{a}$-*OSN*$^{i}$ CIRI, *SMARCD3*- or *KDM6B*-enhanced CIRI lines with *ID3* repression, and *MYOD1* OPTi-OX cells are shown. Scale bars, 200 μm.

(C) UMAP representation of the single-cell RNA-seq dataset of Figure 7E, split by condition. Cells are colored by transcriptional state, annotated based on marker-gene expression (Figure 7F and panel D).

(D) Feature plots showing representative marker genes contributing to annotation of the transcriptional states of panel C.

(E) RT-qPCR analysis of selected lineage and myogenic markers after 7 days of -RA myogenic differentiation, comparing baseline *MYOD1*$^{a}$-*OSN*$^{i}$ CIRI, *SMARCD3*- or *KDM6B*-enhanced CIRI lines with *ID3* repression, and *MYOD1* OPTi-OX cells. N = 4 differentiations; one-way ANOVA followed by Holm–Sidak's multiple-comparison test for all pairwise comparisons. * $p < 0.05$; ** $p < 0.01$; *** $p < 0.001$; $p > 0.05$ not reported.

### Supplemental Tables

| sgRNA::effector | Target | # clones picked | %NT clones (a) | %HOM clones (b) | %HET clones (b) | %BB clones (c) | >90% sfGFP+ (d) |
|---|---|---|---|---|---|---|---|
| com::Com-K | *TFRC* | 6 | 16.6 | 83.3 | 0 | 100 | 100 |
| | NTC | 6 | 16.6 | 83.3 | 0 | 100 | 100 |
| com-cs1::Com-K | *TFRC* | 6 | 16.6 | 83.3 | 0 | 100 | 100 |
| | NTC | 6 | 16.6 | 50 | 33.3 | 100 | 100 |
| com-cs2::Com-K | *TFRC* | 6 | 0 | 50 | 50 | 100 | 100 |
| | NTC | 6 | 0 | 33.3 | 66.6 | 100 | 100 |
| MS2::MCP-PH | *MYOD1* | 12 | 66.6 | 0 | 33.3 | 50 | 100 |
| | NTC | 6 | 0 | 0 | 100 | 66.6 | 66.6 |
| MS2-cs1::MCP-PH | *MYOD1* | 6 | 0 | 0 | 100 | 66.6 | 66.6 |
| | NTC | 6 | 0 | 16.6 | 83.3 | 100 | 50 |
| MS2-cs2::MCP-PH | *MYOD1* | 12 | 16.6 | 66.6 | 16.6 | 80 | 80 |
| | NTC | 6 | 0 | 16.6 | 83.3 | 100 | 50 |
| PP7::PCP-PH | *MYOD1* | 6 | 0 | 50 | 50 | 66.6 | 83.3 |
| | NTC | 6 | 66.6 | 0 | 33.3 | 100 | 100 |
| PP7-cs2::PCP-PH | *MYOD1* | 13 | 30.7 | 7.7 | 53.8 | 77.7 | 100 |
| | NTC | 8 | 0 | 12.5 | 87.5 | 100 | 100 |

**Table S1. Genotyping and flow-cytometry results for sgRNA recruitment architecture testing.**
(a) Percentage of clones with incorrect targeting, defined as absence of 5'- and 3'-integration PCR bands with retention of the WT locus PCR band, or evidence of targeting with incorrect 5'- or 3'-integration PCR product size.
(b) Percentage of clones with correct on-target integration, classified as heterozygous (HET) or homozygous (HOM).
(c) Percentage of correctly targeted clones with additional off-target integration (backbone PCR positivity).
(d) Percentage of clones with >90% GFP expression among correctly targeted clones.

| # Transfection | # clone | WT | 5'INT | 3'INT | 5'BB | 3'BB | % mCherry+ cells |
|---|---|---|---|---|---|---|---|
| A | 1 | yes | no | no | no | no | 0 |
| | 2 | yes | no | yes | yes | no | 0 |
| | 3 | yes | no | no | no | no | 0 |
| | 4 | yes | no | no | no | no | 0 |
| | 5 | yes | no | no | no | no | 0 |
| | 6 | yes | yes | yes | no | no | 3.48 |
| | 7 | yes | no | no | no | no | 0 |
| | 8 | yes | no | no | no | no | 0 |
| | 9 | yes | no | no | no | no | 0 |
| B | 1 | yes | No | no | no | no | 0 |
| | 2 | yes | Yes | yes | yes | yes | 0 |
| | 3 | yes | no | no | no | no | 0 |
| | 4 | yes | no | no | yes | no | 8.75* |
| | 5 | yes | yes | yes | yes | yes | 20.90 |
| | 6 | yes | yes | yes | no | no | 0 |

**Table S2. Genotyping and flow-cytometry results for dCas9-2A-mCherry *CLYBL* targeting.** Two independent transfection experiments were performed, A and B. Most clones were not correctly targeted. Clones A2, B2, and B6 showed discordance between genotyping and mCherry expression, with absent or low detectable mCherry despite evidence of partial or complete targeting. Clone B4 showed low mCherry mean fluorescence intensity (*). Clone A6 was selected because it carried on-target integration without detectable backbone PCR positivity, indicating no additional random integration. However, only 3.48% of cells were mCherry+, suggesting either incomplete clonality, heterogeneous transgene expression, or transgene silencing.

| Application | Target gene | # clone | Genotyping | BB PCR$^+$ | % Effector$^+$ cells |
|---|---|---|---|---|---|
| CRISPRa | *MYOD1* | 1 | +/+ | no | 92.2 |
| | | 2 | +/+ | no | 96.1 |
| | | 3 | +/- | no | 95.5 |
| | *ASCL1* | 1 | +/- | yes | 99.7 |
| | | 2 | +/- | yes | 99.5 |
| | | 3 | +/- | yes | 99.1 |
| | *HNF1A* | 1 | +/- | yes | 99.6 |
| | | 2 | +/- | yes | 98.9 |
| | | 3 | +/- | yes | 98.9 |
| CRISPRi | *TFRC* | 1 | +/+ | no | 98.3 |
| | | 2 | +/+ | no | 98.6 |
| | *NANOG* | 1 | +/- | yes | 99.3 |
| | | 2 | +/+ | yes | 99.7 |
| | | 3 | +/+ | yes | 91.8 |
| | *SOX2* | 1 | +/- | no | 98.33 |
| | | 2 | +/- | no | 98.61 |
| | | 3 | +/+ | no | 98.88 |
| | | 4 | +/+ | no | 99.55 |

**Table S3. Genotyping and flow-cytometry results for inducible CRISPRa and CRISPRi targeting.** The table reports genotyping results for the analyzed clones, including homozygous (+/+) or heterozygous (+/-) on-target integration, backbone-PCR positivity (BB PCR$^+$), and the percentage of effector-2A-sfGFP$^+$ cells confirming expression of the CRISPR effector cassette.

| Configuration | # clones genotyped | % NT (a) | % CRISPRa (b) | % CRISPRi (c) | % CIRI (d) | % no BB (e) |
|---|---|---|---|---|---|---|
| *ASCL1*$^a$-*OSN*$^i$ CIRI | 17 | 0 | 23.5 | 0 | 76.5 | 0 |
| Px3*MYOD1*$^a$-*OSN*$^i$ CIRI | 34 | 0 | 41.2 | 0 | 58.8 | 32.3 |
| NTCx3$^a$-NTCx3$^i$ CIRI | 9 | 0 | 44.5 | 0 | 55.5 | 33.3 |
| Px3*MYOD1* CRISPRa | 15 | 6.6 | 93.3 | / | / | 80 |
| *ASCL1* CRISPRa | 10 | 0 | 80 | / | / | 80 |
| NTCx3 CRISPRa | 4 | 25 | 75 | / | / | 50 |

**Table S4. Genotyping results for CIRI and CRISPRa clones.**
(a) Percentage of clones with incorrect targeting, defined as absence of 5'- and 3'-integration PCR bands with retention of the WT locus band, or evidence of targeting with incorrect 5'- or 3'-integration PCR product size.
(b) Percentage of clones with correct on-target integration of the CRISPRa cassette only.
(c) Percentage of clones with correct on-target integration of the CRISPRi cassette only.
(d) Percentage of clones with correct on-target integration of both CRISPRa and CRISPRi cassettes.
(e) Percentage of correctly targeted clones without detectable backbone PCR positivity.

| Condition | Total cells | ≥1 guide | No guide | CRISPRa-only | CRISPRi-only | CIRI | Unassigned | Guide UMI/cell |
|---|---|---|---|---|---|---|---|---|
| +RA SG | 18,664 | 81.4% | 18.6% | 9.9% | 51.3% | 38.8% | 18.6% | 1,122 |
| -RA SG | 14,274 | 77.4% | 22.6% | 6.7% | 61.2% | 32.0% | 22.6% | 1,162 |
| +RA DG | 20,909 | 80.0% | 20.0% | 16.6% | 73.5% | 10.4% | 19.7% | 1,499 |
| -RA DG | 14,247 | 84.2% | 15.8% | 12.8% | 83.3% | 4% | 15.1% | 2,601 |

**Table S5. Guide-capture QC for single- and dual-sgRNA screens.**
Total cells correspond to confidently mapped barcoded cells before guide-assignment filtering. "≥1 guide" and "No guide" indicate the fraction of cells with or without at least one detected CRISPR protospacer UMI. CRISPRa-only, CRISPRi-only, and CIRI percentages are calculated among guide-assigned cells. SG, single-guide screen; DG, dual-guide screen; UMI per cell, mean detected CRISPR protospacer UMIs per cell.

| Condition | # clones genotyped | % CIRI | % Effector-BFP$^{+*}$ |
|---|---|---|---|
| *MS*$^a$-*OSN*$^i$ CIRI | 20 | 75 | 92.1% |
| *MK*$^a$-*OSN*$^i$ CIRI | 22 | 9 | 93.0% |

**Table S6. Genotyping and flow-cytometry results for screen-hit validation.**
% CIRI indicates the percentage of clones with correct on-target integration of both CRISPRa and CRISPRi cassettes. *Percentage of Effector-2A-TagBFP$^+$ cells in the clone selected for downstream experiments.

| sgRNA target | Top oligo 5’-3’ | Bottom oligo 5’-3’ | Reference |
|---|---|---|---|
| *MYOD1* -130 | GACCGGGCCCCTGCGGCCACCCCG | AAACCGGGGTGGCCGCAGGGGCCC | [14] |
| *MYOD1* -202 | GACCGCTCCCTCCCTGCCCGGTAG | AAACCTACCGGGCAGGGAGGGAGC | [14] |
| *MYOD1* -262 | GACCGAGGTTTGGAAAGGGCGTGC | AAACGCACGCCCTTTCCAAACCTC | [14] |
| *MYOD1* DRR | GACCGGCTGGATTGGGTTTCCAG | AAACCTGGAAACCCAATCCAGCC | [89] |
| *ASCL1* -181 | GACCGCGGGAGAAAGGAACGGGAGG | AAACCCTCCCGTTCCTTTCTCCCGC | [16] |
| *HNF1A* -240 | GACCGACACGGATAAATATGAACCT | AAACAGGTTCATATTTATCCGTGTC | [90] |

#### Table S7. Cloning primers for CRISPRa sgRNAs.

| sgRNA target | Top oligo 5’-3’ | Bottom oligo 5’-3’ | Reference |
|---|---|---|---|
| *NANOG* +21 | GACCGCCAGCAGAACGTTAAAATCC | AAACGGATTTTAACGTTCTGCTGGC | [17] |
| *POU5F1* +22 | GACCGTCGCAAGCCCTCATTTCACC | AAACGGTGAAATGAGGGCTTGCGAC | [17] |
| *SOX2* -1 | GACCGCCCTGACAGCCCCCGTCACA | AAACTGTGACGGGGGCTGTCAGGGC | [17] |
| *TFRC* +3 | GACCGCTCAGAGCGTCGGGATATC | AAACGATATCCCACGCTCTGAGC | [19] |

#### Table S8. Cloning primers for CRISPRi sgRNAs.

| Primer name | Primer sequence 5’-3’ |
|---|---|
| Guide_1 FWD | CTATGGGTCAATTCGGGTAC |
| Guide_1 REV | ACGAGGTCTGCTGTCAAGACACAGCATAGTCGGAACTCCATATATGGGCTA |
| Guide_2 FWD | ACTATGCTGTGTCTTGACAGCAGACCTCGTGAGGGCCTATTTCCCATG |
| Guide_2 REV | GTGTGGTTGCCGACAGTACTTGTGTGTCCACGGAACTCCATATATGGGCTA |
| Guide_3 FWD | TGGACACACAAGTACTGTCGGCAACCACACGAGGGCCTATTTCCCATG |
| Guide_3 REV | GATTACTATTAATAACTAGGACGGTATCG |

#### Table S9. Cloning primers for multi-guide sgRNA cassettes.

| Application | Target gene | sgRNA protospacer | Distance to TSS (bp) | Alignment score | ATAC score |
|---|---|---|---|---|---|
| CRISPRa | *KDM6B* | AGGCGTGGCCCTGAACGAAT[1] | -338 | 3 | 21.35 |
| | | GGCGCGTGATCCAAATGAGT[2] | -234 | 0 | 14.01 |
| | | ACCCCGTACACTGCACCCTG[1] | -184 | 5 | 8.05 |
| | | CCCGCACCCGGCATCCTAGC[2] | -140 | 3 | 41.89 |
| | *MEF2C* | CGGGGGGGATTGAAGGATAC[1] | -346 | 1 | 30.74 |
| | | AAACATCGCGTAAAAAAGAC[2] | -311 | 1 | 31.82 |
| | | CTGGTTACTTTTAATCCGAC[1] | -200 | 2 | 6.85 |
| | | CCTCGGCGCGCGCGAATGCG[2] | -92 | 0 | 14.01 |
| | *MYF5* | TGATTCCTCACGCCCAGGAT[1] | -383 | 3 | 0.30 |
| | | TCTCTAGATAGGCTAAAACA[2] | -242 | 4 | 0.84 |
| | | ACTGCCCCTTTTGACGCTAA[1] | -165 | 0 | 0.27 |
| | | ATATCCCACCGCAACCCCCG[2] | -92 | 0 | 0.63 |
| | *MYF6* | ACCCGGCTATTTTGGAGCAA[1] | -376 | 2 | 0.00 |
| | | TTAGAATTCAGATGAGCGCA[2] | -317 | 3 | 0.14 |
| | | TCTAAACTGCCCAAAATGGA[1] | -231 | 4 | 0.57 |
| | | GCACTAGCTGCAATCACATT[2] | -88 | 6 | 13.33 |
| | *MYOG* | CGACTGATGTAGTGTGGTTA[1] | -481 | 0 | 0.38 |
| | | CCGAAGAAGCTGGTGGGTAT[2] | -386 | 3 | 1.88 |
| | | ACTAATCAAATTACACCCGA[1] | -234 | 2 | 22.03 |
| | | GCACATCAAGGCGTTTACAG[2] | -126 | 2 | 19.31 |
| | *PAX7* | TCTCGGAGTCCCGGCTGTAT[1] | -379 | 1 | 15.50 |
| | | GAGCCGATCCCGGCGAGTTG[2] | -255 | 0 | 3.05 |
| | | CCGGCTCGACCTCGTTTTGG[1] | -204 | 0 | 15.23 |
| | | TGAGCGCGATCTGATAGGTT[2] | -73 | 1 | 11.97 |
| | *SMARCD3* | ATTCGCCGGCCGCGCCAGGC[1] | -386 | 2 | 603.82 |
| | | CTATTTGTGGCGAGAGCCGG[2] | -336 | 0 | 7.45 |
| | | CGCCGCTTATCTGCGCTGTT[1] | -193 | 1 | 5.36 |
| | | CAGAAATAGTCCGGCGGCCC[2] | -128 | 0 | 9.85 |
| CRISPRi | *ATF7IP* | TAACGGCCCCGGCGCGTGCA[1] | -24 | 0 | 2.50 |
| | | TGTTTTTGAATCTGCGGAGG[2] | 11 | 7 | 182.51 |
| | | AAAGCTGAGGCGGCAACGTC[1] | 73 | 1 | 6.23 |
| | | GCTGCGCGGGACGGCTCTGT[2] | 100 | 4 | 3.78 |
| | *CTCF* | TGGAGCGATTAAACCGTGCG[1] | 8 | 0 | 2.22 |
| | | GCACGGTTTAATCGCTCCAC[2] | -8 | 1 | 2.22 |
| | | GCGGAGCTCGCCGGAGACGC[1] | 79 | 3 | 4.52 |
| | | TGCGGACGCGCGGAGCTCGC[2] | 70 | 1 | 3.84 |
| | *ID3* | GGCTCTATAAGTGACCGCCG[1] | -20 | 0 | 3.95 |
| | | CACTGTAGCGGGACTTCTTT[2] | 25 | 2 | 16.92 |
| | | GGTGCGCGGCTGCTACGAGG[1] | 106 | 0 | 7.04 |
| | | CGCAGTCTGGCCATCGCCCG[2] | 146 | 1 | 7.83 |
| | *YBX1* | ACCGATCGAACTAGCGAGAA[1] | 4 | 0 | 20.13 |
| | | GCCATTCTCGCTAGTTCGAT[2] | 15 | 0 | 23.42 |
| | | CCTAGGGCGGGTCGCTCGTA[1] | -25 | 1 | 429.48 |
| | | GCCTAGGGCGGGTCGCTCGT[2] | -26 | 1 | 4.37 |
| | *ZNF207* | CTCCCGTCAAGCACTGCGGT[1] | -37 | 3 | 217.59 |
| | | GGAGCGGGGAACGAGGCCGT[2] | 0 | 12 | 20.70 |
| | | CGGCTACCACCACGTCCCAC[1] | 28 | 6 | 15.94 |
| | | TTGCCGGTAGAACACAGTTA[2] | 128 | 4 | 14.96 |

**Table S10. Designed CRISPRa/i sgRNAs for screening targets.** [1] and [2] indicate guide-pair assignments used in the dual-guide setting. For the single-guide setting, the first two guides for each target were selected and used separately. ATAC scores are reported with two decimal places.

| Locus | PCR type | Plasmid | Primer binding site | Primer sequence |
|---|---|---|---|---|
| *hROSA26* | Locus PCR | pR26-Bst_ CAG-OPTtetR | Genome (5') | GAGAAGAGGCTGTGCTTCGG |
| | | | Genome (3') | ACAGTACAAGCCAGTAATGGAG |
| | 5'-INT PCR | pR26-Bst_ CAG-OPTtetR | Genome (5') | GAGAAGAGGCTGTGCTTCGG |
| | | | Splice acceptor | AAGACCGCGAAGAGTTTGTCC |
| | 3'-INT PCR | pR26-Bst_ CAG-OPTtetR | bGH poly(A) | GAGAATAGCAGGCATGCTG |
| | | | Genome (3') | ACAGTACAAGCCAGTAATGGAG |
| | 5'-BB PCR | pR26-Bst_ CAG-OPTtetR | Vector backbone (5') | CGTTGTAAAACGACGGCCAG |
| | | | Splice acceptor | AAGACCGCGAAGAGTTTGTCC |
| | 3'-BB PCR | pR26-Bst_ CAG-OPTtetR | bGH poly(A) | GAGAATAGCAGGCATGCTG |
| | | | Vector backbone (3') | TGACCATGATTACGCCAAGC |
| *CLYBL* | Locus PCR | pCLYBL-Neo_ CAG-dCAS9-2A-mCherry | Genome (5') | TAAGTGACCCCTGGCGAGAC |
| | | | Genome (3') | AGATGGAGCAGTGGATGACA |
| | 5'-INT PCR | pCLYBL-Neo_ CAG-dCAS9-2A-mCherry | Genome (5') | TAAGTGACCCCTGGCGAGAC |
| | | | T2A-P2A | AAGACTTCCTCTGCCCTCTC |
| | 3'-INT PCR | pCLYBL-Neo_ CAG-dCAS9-2A-mCherry | HS4 insulator | TCCAGGACGGAGTCAGTGAG |
| | | | Genome (3') | AGATGGAGCAGTGGATGACA |
| | 5'-BB PCR | pCLYBL-Neo_ CAG-dCAS9-2A-mCherry | Vector backbone (5') | CACAGGAAACAGCTATGACC |
| | | | T2A-P2A | AAGACTTCCTCTGCCCTCTC |
| | 3'-BB PCR | pCLYBL-Neo_ CAG-dCAS9-2A-mCherry | HS4 insulator | TCCAGGACGGAGTCAGTGAG |
| | | | Vector backbone (3') | TTTTCCCAGTCACGACGTTG |
| *AAVS1* | Locus PCR | All pAAV plasmids | Genome (5') | CTGTTTCCCCTTCCCAGGCAGGTCC |
| | | | Genome (3') | TGCAGGGGAACGGGGCTCAGTCTGA |
| | 5'-INT PCR | pAAV-Puro plasmids | Genome (5') | CTGTTTCCCCTTCCCAGGCAGGTCC |
| | | | PuroR | TCGTCGCGGGTGGCGAGGCGCACCG |
| | | pAAV-Hygro plasmids | Genome (5') | CTGTTTCCCCTTCCCAGGCAGGTCC |
| | | | HygroR | CCGCAGGACATATCCACGCCCTC |
| | 3'-INT PCR | All pAAV plasmids | bGH poly(A) linker | AACGCGTAGCTCGCTGATC |
| | | | Genome (3') | TGCAGGGGAACGGGGCTCAGTCTGA |
| | | pAAV with BFP plasmid | TagBFP | GCCAGATACTGCGACCTCCCTAG |
| | | | Genome (3') | TGCAGGGGAACGGGGCTCAGTCTGA |
| | | pAAV with KRAB plasmid | KRAB | GCGGCTGGAAAAGGGCGAAG |
| | | | Genome (3') | TGCAGGGGAACGGGGCTCAGTCTGA |
| | 5'-BB PCR | pAAV-Puro plasmids | Vector backbone (5') | TGAGGAAGAGTTCTTGCAGCTC |
| | | | PuroR | ATGCTTCCGGCTCGTATGTT |
| | | pAAV-Hygro plasmids | Vector backbone (5') | TGAGGAAGAGTTCTTGCAGCTC |
| | | | HygroR | CCGCAGGACATATCCACGCCCTC |
| | 3'-BB PCR | All pAAV plasmids | bGH poly(A) linker | AACGCGTAGCTCGCTGATC |
| | | | Vector backbone (3') | ATGCACCACCGGGTAAAGTT |
| | | pAAV with BFP plasmid | TagBFP | GCCAGATACTGCGACCTCCCTAG |
| | | | Vector backbone (3') | ATGCACCACCGGGTAAAGTT |
| | | pAAV with KRAB plasmid | KRAB | GCGGCTGGAAAAGGGCGAAG |
| | | | Vector backbone (3') | ATGCACCACCGGGTAAAGTT |

**Table S11. Genotyping primers.**

| Gene | Forward primer sequence 5’-3’ | Reverse primer sequence 5’-3’ |
|---|---|---|
| *ASCL1* | GCAACCGGGTCAAGTTGGT | GTCGTTGGAGTAGTTGGGGG |
| *CA3* | ATCTTCACTGGGGCTCTTCG | AACCAAATGAAGCTCCGCTG |
| *CDKN1C* | CGATCAAGAAGCTGTCCGGG | GGCTCTTTGGGCTCTAAATTGG |
| COM | GGCTCTGACTGACCGCGTTA* | TTGTTGCAGTTCTTGCACCG |
| dCas9 | GGCTCTGACTGACCGCGTTA* | TTGCTGGGCACCTTGTACTC |
| *HNF1A* | TGGCCATGGACACGTACAG | GCTGCTTGAGGGTACTTCTG |
| *HPRT1* | TGACACTGGCAAAACAATGCA | GGTCCTTTTCACCAGCAAGCT |
| *ID3* | CTACAGCGCGTCATCGACTA | TGAGCTCGGCTGTCTGGATG |
| *KDM6B* | ACTGCCTCCACCACCATTAC | GCATCTCCATGCAGGGTACT |
| *MAP2* | AGACTGCAGCTCTGCCTTTAG | AGGCTGTAAGTAAATCTTCCTCC |
| MCP | GGCTCTGACTGACCGCGTTA* | TGATCCATTCAGCGATCCCG |
| *MEF2C* | GGACAAGGAATGGGAGGA | TGAGTAGAAGGCAGGGAGAG |
| *MYH3* | TGCTGCATACCCAGAACACC | CCCTGCTGGCATCTTCTACC |
| *MYH11* | CCGGGAAAACCGAAAACACC | AGATGGGCCTTGCGTGATAC |
| *MYOD1* | GCCGCTTTCCTTAACCACAA | CTGAATGCCCACCCACTGTC |
| *MYOG* | AGTGCCATCCAGTACATCGAGC | AGGCGCTGTGAGAGCTGCATTC |
| *NANOG* | CATGAGTGTGGATCCAGCTTG | CCTGAATAAGCAGATCCATGG |
| *OLIG3* | AGGCTGAAGATCAACGGACG | TGGCGATCTTGGAGAGCTTG |
| OPTtetR | CCACCGAGAAGCAGTACGAG | TTTCCTCTTTGGCGACCTGG |
| *PAX7* | ACCCCTGCCTAACCACATC | GCGGCAAAGAATCTTGGAGAC |
| PCP | GGCTCTGACTGACCGCGTTA* | TCGAAGATCTGACGGTCTGC |
| *POU5F1* | AGTGAGAGGCAACCTGGAGA | ACACTCGGACCACATCCTTC |
| *RPLP0* | GGCGTCCTCGTGGAAGTGAC | GCCTTGCGCATCATGGTGTT |
| *SMARCD3* | TTATGGCCCTGACAACCACC | TTGAACTGGGGAGGCTGGTA |
| *SOX2* | TGGACAGTTACGCGCACAT | CGAGTAGGACATGCTGTAGGT |
| *TFRC* | CAGAGCAGACATAAAGGAAATGGG | CCAGAAGACATGTCGGAAAGG |
| *TNNI1* | ACCATGCCGGAAGTCGAGAG | TAGCGCACCTTCTCAGCCTC |
| *TTN* | GTAAAAAGAGCTGCCCCAGTGA | GCTAGGTGGCCCAGTGCTACT |

**Table S12. RT-qPCR primers.** *Common forward primer binding to an exon within the shared CAG promoter; reverse primers bind downstream of an intervening intron, favoring amplification of spliced cDNA over genomic DNA.

| Antigen | Species | Clonality | Dilution | Application |
|---|---|---|---|---|
| $\alpha$-actinin-2 (ACTN2) | Mouse | monoclonal | 1:100 | IF |
| GAPDH | Mouse | monoclonal | 1:1,000 | WB |
| MYOD1 | Rabbit | monoclonal | 1:100 | IF |
| NANOG | Mouse | monoclonal | 1:100 | Flow cytometry |
| POU5F1/OCT4 | Mouse | monoclonal | 1:100 | Flow cytometry |
| Vinculin (VCL) | Mouse | monoclonal | 1:10,000 | WB |
| dCas9 | Mouse | monoclonal | 1:250–1:1,000* | IF, WB |
| TetR | Rabbit | polyclonal | 1:4,000–1:1,000* | IF, WB |
| Isotype IgG1 | Mouse | monoclonal | 1:100 | Flow cytometry |

**Table S13. Antibodies.** *Dilutions are reported in the same order as the corresponding applications.
